# Conserved small nucleotidic elements at the origin of concerted piRNA biogenesis from genes and lncRNAs

**DOI:** 10.1101/2020.02.05.936112

**Authors:** Silke Jensen, Emilie Brasset, Elise Parey, Hugues Roest-Crollius, Igor V. Sharakhov, Chantal Vaury

**Affiliations:** GReD, Université Clermont Auvergne, CNRS, INSERM, Faculté de Médecine, 63000 Clermont-Ferrand, France; Institut de Biologie de l’Ecole Normale Supérieure (IBENS), Ecole Normale Supérieure, CNRS, INSERM, PSL Research University, 75005, Paris, France; Department of Entomology, Virginia Polytechnic Institute and State University, Blacksburg, VA 24061, USA; Department of Cytology and Genetics, Tomsk State University, Tomsk, 634050, Russia

**Keywords:** phased piRNA biogenesis, piRNA, lncRNA, malaria mosquitoes, germline

## Abstract

PIWI-interacting RNAs (piRNAs) target transcripts by sequence complementarity serving as guides for RNA slicing in animal germ cells. The piRNA pathway is increasingly recognized as critical for essential cellular functions such as germline development and reproduction. In the *Anopheles gambiae* ovary, as much as 11% of piRNAs map to protein-coding genes. Here we show that ovarian mRNAs and long non-coding RNAs (lncRNAs) are processed into piRNAs that can direct other transcripts into the piRNA biogenesis pathway. Targeting piRNAs fuel transcripts either into the ping-pong cycle of piRNA amplification or into the machinery of phased piRNA biogenesis, thereby creating networks of inter-regulating transcripts. RNAs of the same network share related genomic repeats. These repeats give rise to piRNAs, which target other transcripts and lead to a cascade of concerted RNA slicing. While ping-pong networks are based on repeats of several hundred nucleotides, networks that rely on phased piRNA biogenesis operate through short ∼40-nucleotides long repeats, which we named snetDNAs. Interestingly, snetDNAs are recurring in evolution from insects to mammals. Our study brings to light a new type of a conserved regulatory pathway, the snetDNA-pathway, by which short sequences can include independent genes and lncRNAs in the same biological pathway.

**AUTHOR SUMMARY:** Small RNA molecules are essential actors in silencing mobile genetic elements in animal germ cells. The 24-29-nucleotide-long Piwi-interacting RNAs (piRNAs) target transcripts by sequence complementarity serving as guides for RNA slicing. Mosquitoes of the *Anopheles gambiae* species complex are the principal vectors of malaria, and research on their germline is essential to develop new strategies of vector control by acting on reproduction. In the *Anopheles gambiae* ovary as much as 11% of piRNAs originate from protein-coding genes. We identified piRNAs which are able to target transcripts from several distinct genes or long non-coding RNAs (lncRNAs), bringing together genic transcripts and lncRNAs in a same regulation network. piRNA targeting induces transcript slicing and production of novel piRNAs, which then target other mRNAs and lncRNAs leading again to piRNA processing, thus resulting in a cascade of RNA slicing and piRNA production. Each network relies on piRNAs originating from repeated genetic elements, present in all transcripts of the same network. Some of these repeats are very short, only ∼40-nucleotides long. We identified similar repeats in all 43 animal species that we analysed, including mosquitoes, flies, arachnidae, snail, mouse, rat and human, suggesting that such regulation networks are recurrent, possibly conserved, in evolutionary history.

## INTRODUCTION

PIWI-interacting small RNAs (piRNAs), 24 to 29 nucleotides (nt) long, are essential actors in controlling transposable elements (TEs) in animal gonads. Specific genomic loci called piRNA clusters, of tens or even hundreds of kilobases (Kb), produce large amounts of piRNAs. Owing to sequence complementarity, piRNAs from piRNA clusters guide PIWI proteins to cleave TE transcripts and induce their processing into new piRNAs. These new piRNAs can themselves act as PIWI guides for the cleavage of complementary transcripts inducing production of piRNAs that are identical to the “initiator” piRNAs that originated from the piRNA clusters. This leads to a piRNA amplification process, which has been called the ping-pong cycle (1, 2), reviewed in (3). piRNAs display specific characteristics related to their biogenesis and to ping-pong amplification. piRNAs that originate from ping-pong amplification show a specific 10-nt 5’-overlap due to the fact that PIWI proteins, slice their targets between nucleotides 10 and 11 of their guide. In *Drosophila melanogaster*, piRNAs bound to the PIWI protein Aubergine (Aub), show a “1U” signature corresponding to a high proportion of uracil at their 5’-end. piRNAs bound to Argonaute 3 (Ago3), often have an adenine at position 10. This “10A” signature corresponds to the nucleotide that faces the 5’-uracil of the complementary piRNA and is attributed to a preference of Aub for an A at this position of the target RNA (1, 4).

Phased piRNA biogenesis in the *D. melanogaster* germline is also operating with Aub and Ago3 as ping-pong does but it doesn’t necessarily lead to an amplification loop (5, 6). It leads to production of piRNAs, initiated by the slicing of a transcript by Ago3 or Aub. In this process, an initial Ago3/Aub-bound piRNA, a “trigger-piRNA” (5) or “initiator” (3), targets a complementary transcript. This results in cleavage of the transcript and production of “responder-piRNAs”, essentially Aub-bound, which have 5’-overlap of 10 nt with the trigger-piRNAs. Then, piRNA biogenesis can spread downstream to new regions in a 3’-directed phased manner from the responder-piRNAs and give rise to so-called “trail-piRNAs” (5) or “trailing piRNAs” (3) (Figure 1). Trail-piRNAs are essentially bound to Piwi protein and present “1U” signature in *D. melanogaster* (5, 6).

**Figure 1.**
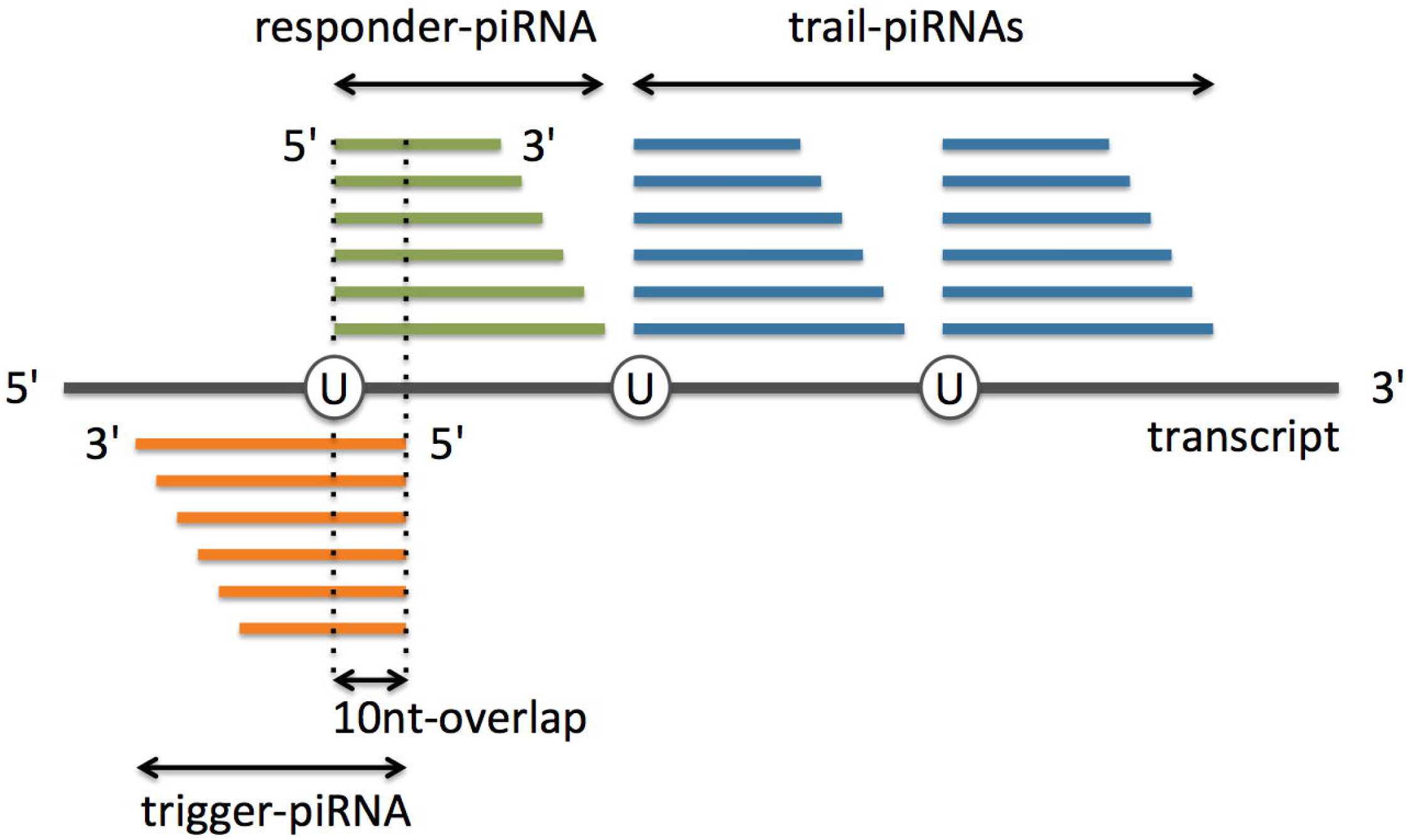
Phased piRNA biogenesis in *Drosophila melanogaster*. Trigger-piRNAs anneal partially complementary transcripts inducing slicing of the latter to responder-piRNAs that have 10-nt 5’-overlap with trigger-piRNAs. Further slicing of the transcript at uridine residues downstream of the responder-piRNA leads to production of trail-piRNAs. 5’-uridine is largely over-represented in trail-piRNAs. Ping-pong between trigger- and responder-piRNAs is marked by over-representation of 1U and 10A. (adapted from (5)).

TE-related sequences are not the only producers of piRNAs. In several organisms, piRNAs originating from the 3’ untranslated regions (UTR) of gene transcripts have been reported (7–10). Several studies indicate that piRNAs play a role beyond TE silencing (reviewed in (8) and can be involved in diverse cellular processes like programmed genome rearrangement and genome-wide surveillance of germline transcripts (11, 12), mRNA or long non-coding RNA (lncRNA) regulation and development ((13–18), reviewed in (19, 20)).

Here, we sought to address whether piRNAs, if produced from protein-coding genes and unrelated to any TE-sequences, may lead to piRNA-guided mRNA cleavage and establish a piRNA regulation network dedicated to gene regulation. To explore this possibility, we studied piRNAs from *Anopheles gambiae* ovaries. The African malaria vector *An. gambiae* particularly suits this analysis because first, piRNAs with a ping-pong signature are produced in the ovarian tissue (21–23) and second, a high proportion of them, 11%, are derived from protein-coding transcripts (21). We here uncover piRNA biogenesis in *An. gambiae* resulting from ping-pong amplification between transcripts from distant genes. We identify a novel “intrinsic ping-pong amplification”, which is achieved with genome-unique piRNAs originating from the same single gene. We also evidence phased piRNA biogenesis from genes and lncRNAs, which is initiated by trigger-piRNAs that are not related to nowadays known TEs and originate from other genes and lncRNAs at distant loci. When examined across distant species, we further bring evidence for a new type of regulatory pathway, named snetDNA-pathway, conserved throughout evolution and based on particularly short genomic sequences, the snetDNAs (“small-RNA-network DNA”), which establish regulatory networks through piRNA targeting and phased piRNA biogenesis.

## RESULTS

### Genes may give rise to piRNAs implicated in ping-pong amplification in *An. gambiae* ovaries

Small RNAs from *An. gambiae* ovaries show a bi-modal length distribution with one peak at 22 nt attributed to microRNAs and a broad peak spanning 24–29 nt. Here we analysed only small RNAs of 24-29 nt length, the putative piRNAs.

While analysing this piRNA population, we focused on piRNAs which map to protein coding genes, without mismatch, and found that 870 mRNAs map more than 3 Reads Per Million (RPM) of piRNAs. piRNAs mapping to these 870 mRNAs make up 10.4% of all genome-mapping piRNAs. Among the mRNA-mapping piRNAs, 93.3 %, are genome-unique, i.e. they map only once in the *An. gambiae* genome. The majority of the 870 mRNAs produce only sense piRNA reads (564 transcripts), four mRNAs produce only antisense, and 302 mRNAs map both sense and antisense piRNA reads (Figure 2A).

**Figure 2.**
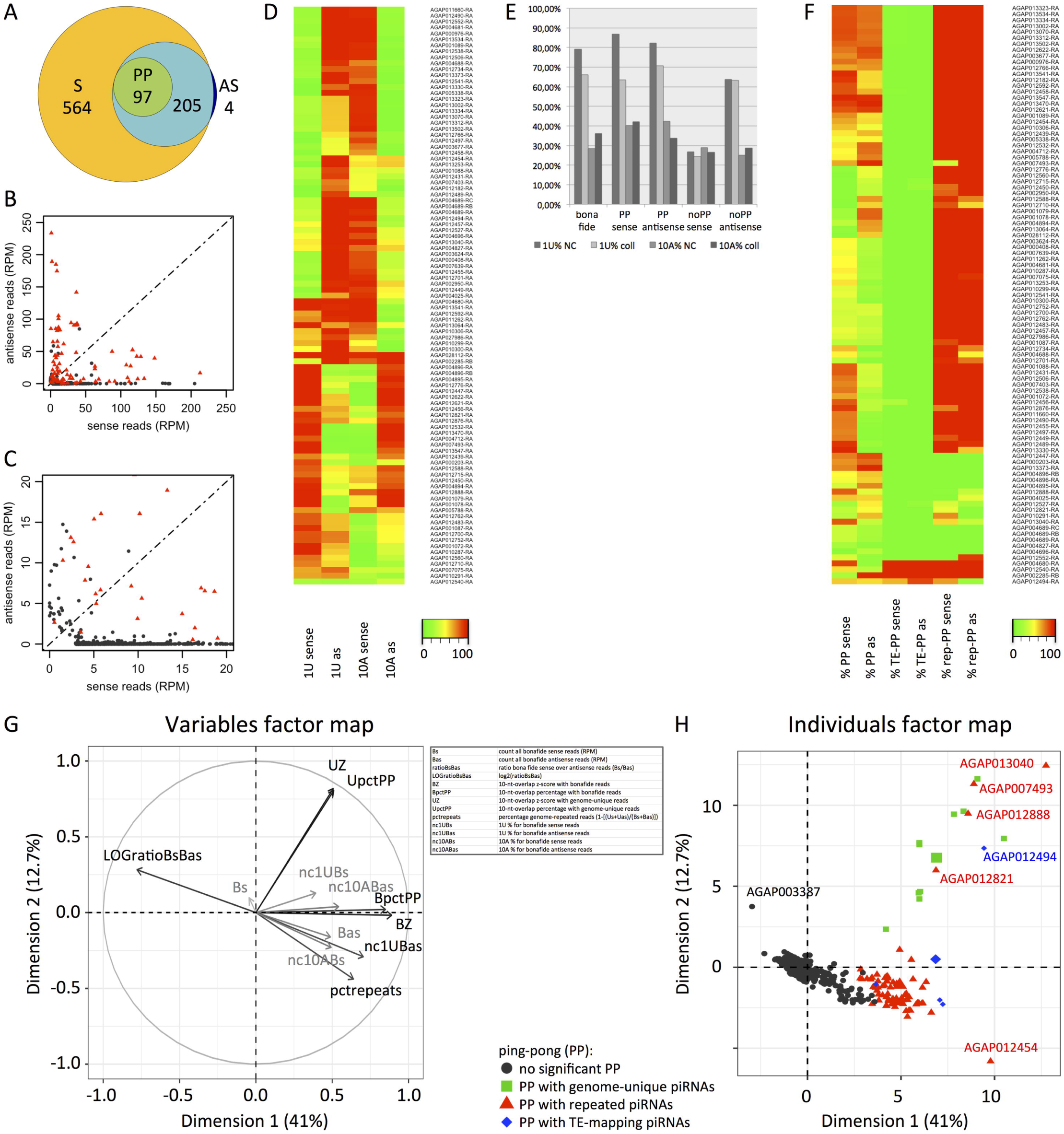
Protein-coding mRNAs matching >3RPM piRNAs. (A) Area-proportional Euler Diagram: mRNAs matching sense piRNAs (S), antisense piRNAs (AS) and piRNAs with significant 10-nt 5’-overlap ping-pong signature (PP). 564 mRNAs match only sense piRNAs, 4 only antisense, and 302 match both sense and antisense piRNAs; 97 of the latter match piRNAs with significant 10-nt 5’-overlap ping-pong signature, 205 mRNAs match piRNAs without it. (B) Scatterplot showing normalized counts of sense and antisense piRNAs (RPM), restricted to up to 250 RPM, for mRNAs matching >3RPM piRNAs. Red spots: mRNAs with significant 10-nt 5’-overlap signature. Grey spots: mRNAs without significant 10-nt 5’-overlap signature (C) Zoom-in from scatterplot in (B). (D) Heat map of 1U and 10A percentages for sense and antisense (as) *bona fide* piRNAs mapping the 97 mRNAs that show significant 10-nt 5’-overlap. (E) Percentages of 1U and 10A signatures from non-collapsed (NC) and collapsed (coll) libraries (see “Methods” section), for *bona fide* and for genic mRNAs from annotated protein-coding genes, AGAP003387 excepted, with or without significant ping-pong signature (“PP” or “noPP,” respectively). (F) For the 97 mRNAs, heat map showing percentages of piRNAs with ping-pong partner among sense and antisense (as) oriented piRNAs (PP), percentages of piRNAs that map insect TEs, allowing 0-3 mismatches, among piRNAs with ping-pong partner (TE-PP), and percentages of piRNAs that map repeated sequences in the genome among piRNAs with ping-pong partner (rep-PP). (G, H) Principal component analysis.

When sense and antisense piRNAs are detected, there is a possibility of ping-pong amplification (1). To test whether genic piRNAs may be amplified by the ping-pong mechanism, we searched sense and antisense piRNAs that have a 5’-overlap of 10 nt (based on their 5’-end positions when mapped to the transcripts). We found a significant over-representation of 10-nt 5’-overlaps (defined in Methods) for 97 transcripts out of the 302 mRNAs mapping both sense and antisense piRNAs (Figure 2A-C, Table S1). The median count of piRNAs is 2-times higher for these 97 transcripts than for the other 773 piRNA-mapping transcripts without enriched 10-nt 5’-overlaps (Supplemental Figure SFig1). The piRNAs from these 97 transcripts also show significant 1U and 10A over-representation (Figures 2D, E). Actually, more than half (55%) of the 97 mRNAs have 1U-biased antisense piRNAs together with 10A-biased sense piRNAs, one third (32%) have 1U-biased sense piRNAs and 10A-biased antisense piRNAs (>50% 1U or 10A respectively, Figure 2D, Table S1). Globally, for the 97 ping-pong positive mRNAs, 1U and 10A signatures are equally present for sense and antisense reads. On the opposite, sense-oriented piRNAs mapping mRNAs without ping-pong signature are neither 1U-nor 10A-enriched and their antisense piRNAs show only significant 1U bias, but no 10A enrichment. This is true for both non-collapsed and collapsed libraries (Figure 2E). The latter unexpected data reveal a new category of small RNAs, which might be specific to mRNAs, 24-29 nt long, without any enriched 1U or 10A.

Until now, ping-pong amplification of endogenous piRNAs was known for TEs. Thus we checked whether the ping-pong observed for 97 mRNAs might be due to the presence of TE-homologous sequences. We tested whether the piRNAs with ping-pong partner, the “ping-pong-piRNAs” (with 10-nt 5’-overlap) map TEs. We found only four transcripts with a high proportion of ping-pong-piRNAs mapping TEs (Figure 2F, Table S1). Consequently, ping-pong for mRNAs is mostly unlinked to the presence of TE-related sequences. We then checked whether the ping-pong-piRNAs map to other, repeated, sequences in the *An. gambiae* genome. The results show that for 82 out of 97 transcripts, the ping-pong-piRNAs map to repeats other than TEs. For the remaining 11 transcripts, all ping-pong-piRNAs map to genome-unique sequences, which are not related to any repeats or TEs.

To extract the important information from our data concerning piRNAs mapping protein-coding transcripts (Table S1), we performed a Principal Component Analysis (PCA) (24) (Figure 2G, Individuals’ coordinates and variables’ contribution are given in Table S2, a correlogram for the PCA variables is shown in Supplemental Figure SFig2, the corresponding raw data are presented in Tables S3 and S4). *Bona fide* piRNAs, defined as all small RNAs, 24-29 nt in length, which do not map miRNAs, rRNAs, snRNAs or tRNAs (see also “Methods”), and genome-unique piRNAs were considered separately. The variables that had the highest contribution to variance were the z-score and the percentage of 10-nt 5’-overlaps, for *bona fide* piRNAs (BZ and BpctPP respectively) and for genome-unique piRNAs (UZ and UpctPP respectively), the ratio of *bona fide* sense over antisense piRNAs (LOGratioBsBas), and the percentage of genome-repeated piRNAs (pctrepeats). The PCA distinguishes essentially three groups of transcripts mapping piRNAs: mRNAs without significant ping-pong, mRNAs mapping repeated or TE-matching piRNAs doing ping-pong, and mRNAs mapping genome-unique piRNAs doing ping-pong (Figure 2H).

Eleven transcripts achieve ping-pong amplification with genome-unique piRNAs (Table S1). These 11 mRNAs originate from 8 different genes. Bar plots of mapping piRNAs illustrate the high proportion of piRNAs with ping-pong partners originating from the same gene (Figure 3). We checked whether piRNAs from other loci may also target these transcripts by mismatched pairing (allowing up to 5 mismatches) and trigger ping-pong amplification. We found that such “trans ping-pong” (4) is possible for these transcripts but piRNAs with ping-pong partners originating from the same gene are the most abundant (Figure 3 and Table S5). Thus, our data provide the first evidence that ping-pong amplification may be achieved with genome-unique piRNAs originating from the same single genomic sequence. We propose to call this ping-pong amplification “intrinsic ping-pong”.

**Figure 3.**
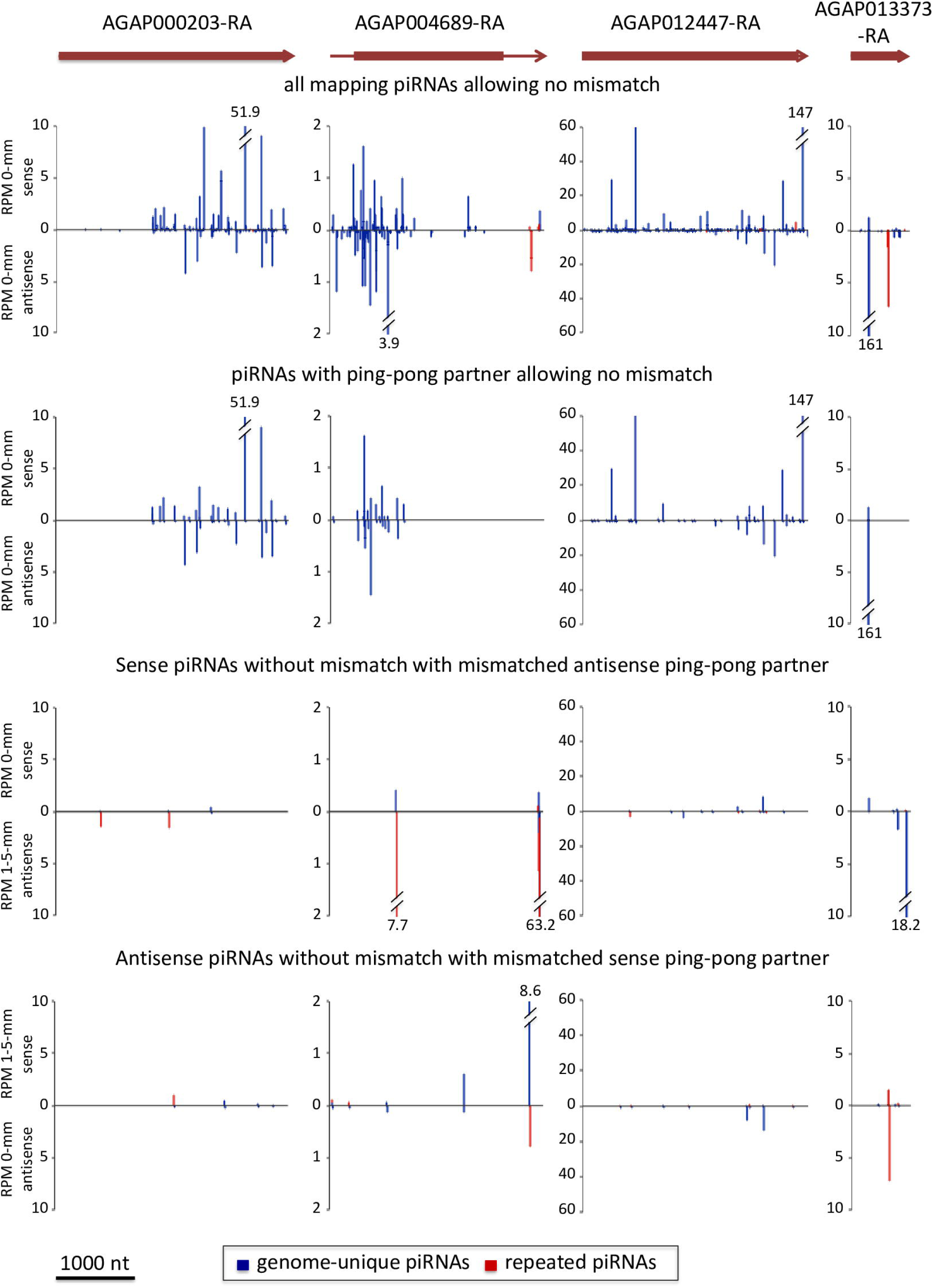
piRNAs mapping transcripts with >95% genome-unique mappers and with significant 10-nt 5’-overlap signature. Barplots of piRNA 5’-positions for four protein-coding transcripts, AGAP000203-RA (inositol-1,4,5-trisphosphate 5-phosphatase), AGAP004689-RA (hypothetical protein without identified protein domains), AGAP012447-RA (ionotropic glutamate receptor GLURIIe), and AGAP013373-RA (homeobox protein). Structure and orientation of the transcripts are shown, coding sequences as a box, untranslated regions (UTR) as a line (Release AgamP4.2). Counts (RPM) of genome-unique (blue) and repeated (red) piRNAs for all mapping piRNAs (top) and for piRNAs having ping-pong partners (second, third and bottom rows). In the first and second row only piRNAs matching the transcripts without any mismatch (“0-mm”) are shown, in the third row sense piRNAs matching the transcripts without mismatch together with mismatched (1 to 5 mismatches, “1-5-mm”) antisense ping-pong partners, in the bottom row antisense piRNAs matching the transcripts without mismatch together with mismatched sense ping-pong partners.

### piRNAs from genomic repeats may fuel mRNAs from distant loci into the ping-pong cycle of piRNA amplification and create “ping-pong-networks”

We focused on the group of mRNAs mapping repeated piRNAs involved in ping-pong. The aim of this study was to find out whether piRNAs originating from one gene may target transcripts of other genes and engage them in ping-pong amplification. We searched for piRNAs with ping-pong signature that may target several genes. We identified 58 mRNAs forming 17 different groups, the mRNAs of each group being targeted by the same ensemble of piRNAs originating from a repeated sequence, which is present in all transcripts of the same group. Actually, these 17 groups represent what we named “ping-pong-networks” (Table S6). To check effective piRNA production for the 58 transcripts of the 17 putative ping-pong-networks, we searched for genome-unique piRNAs mapping these transcripts. There may be spreading to the flanking sequences, or erroneous annotation of the transcripts omitting part of the UTRs as has been observed for AGAP003266-RA (see (25) and below). Thus, we also analysed the flanking sequences, which could in fact produce piRNAs. We found that 32 of the 58 transcripts match >0.5 RPM genome-unique piRNAs, and when including flanking sequences (5 kb up- and downstream), 52 of the 58 transcript loci match >0.5 RPM genome-unique piRNAs. These results indicate that at least 52, i.e. 90%, of the 58 ping-pong-network transcripts correspond indeed to piRNA producing loci (Table S7). To identify the repeats, which are common to the transcripts of each ping-pong-network, we made multiple sequence alignments and built the corresponding sequence logos (supplementary information SI-MSA1). The results evidence different repeats of 130 to 1,130 nucleotides. These repeats and the piRNAs produced from the ping-pong networks do not contain any low complexity sequences (e.g. repeats of di- or trinucleotides). Only one of the two repeats of the ping-pong network 12, repeat A, contains several grouped CAG triplets (see supplementary information SI-MSA1), but these sequences do not match any piRNAs. Consequently, there is no bias of piRNA matching due to low complexity repeats.

Our data show that piRNA biogenesis may occur by ping-pong amplification between mRNAs from a group of genes that contain related repeats. Because mRNAs are sliced when processed to piRNAs, this mechanism may constitute a basis for concerted gene regulation within each ping-pong network.

### piRNAs from genes and lncRNAs may direct other transcripts to phased piRNA biogenesis thus creating networks of inter-regulation

In the *Drosophila* germline, TE-derived piRNAs, the “trigger-piRNAs”, may target genic mRNAs and lead to piRNA-guided slicing of the latter and phased piRNA biogenesis (5, 6). This process is different from ping-pong by the fact that it does not lead to a piRNA amplification loop, but to the production of sense-oriented phased “responder-” and “trail-piRNAs” (5) (Figure 1). We decided to examine whether piRNAs, which originate from genic transcripts or lncRNAs may be complementary to other transcripts and initiate their phased slicing. To achieve this, 24-29 nt reads were mapped against the genome allowing no mismatch, and only genome-unique reads (that map only once to the genome) that match mRNAs or lncRNAs were considered for further analyses. This has the advantage that the origin of these piRNAs is clearly known. To find potential targets, these piRNAs were aligned in a reverse complementary manner, allowing 1 to 5 mismatches, to the 664 mRNAs and 20 lncRNAs that match >3RPM genome-unique sense-oriented piRNAs without mismatch (Table S8). We identified 310 transcripts from 282 genes and 5 different lncRNAs that may be targeted by >3 RPM piRNAs (Table S8). In *Drosophila*, trigger-piRNAs show a precise 10-nt 5’-overlap to the responder-piRNAs that are produced by slicing of the target transcript (Figure 1) (5, 6). Thus, we searched for pairs of putative trigger- and responder-piRNAs showing the typical 10-nt 5’-overlap. We identified a total of 209 RPM trigger-piRNAs, all genome-unique, and 88079 RPM corresponding responder-piRNAs, 88069 RPM of them genome-unique and 10 RPM repeated twice in the genome, originating from five different transcripts (three mRNAs and two lncRNAs). They correspond to 12 trigger/responder pairs, constituting 12 different putative trigger events (Table S9). These trigger events result in a network shown in Figure 4A. It shows where the trigger-piRNAs originate from, and which transcripts they target, together with the amounts of the respective piRNAs. Complementarity of the trigger-piRNAs to the targeted transcripts (mismatched) and of all piRNAs originating from the targeted transcripts (no mismatch), including responder- and trail-piRNAs, are shown in Figure 4B-E and in Supplemental Figure SFig3. Notably, the same trigger-piRNAs may target both different transcripts and several sites within the same transcript.

**Figure 4.**
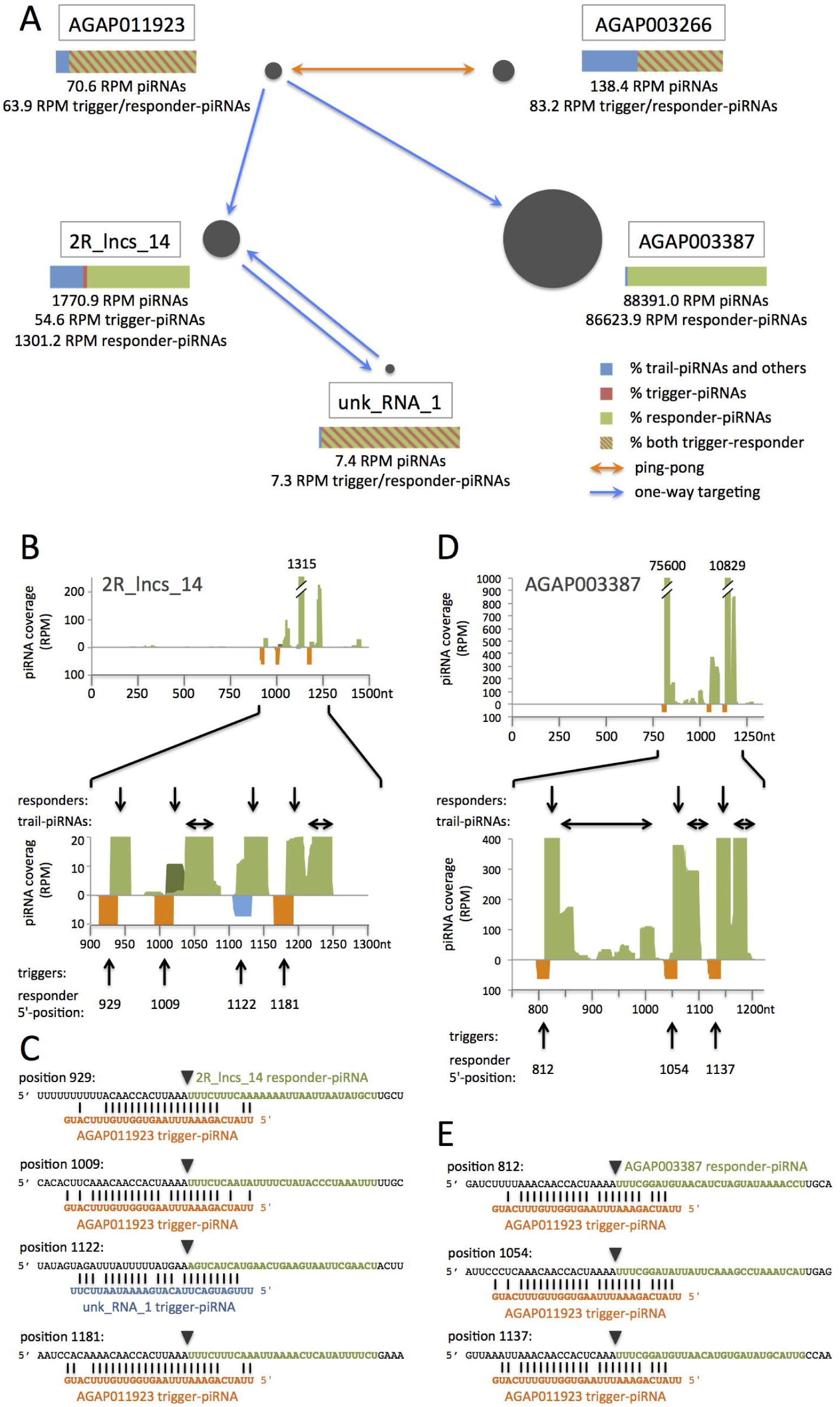
piRNA-based network involving mRNAs and lncRNAs in *An. gambiae*. (A) Schematic representation of the network. Trigger-piRNAs from AGAP011923 target transcripts from AGAP003266 and AGAP003387 and 2R_lncs_14. Resulting responder-piRNAs from AGAP003266 target back AGAP011923 transcripts leading to trans ping-pong amplification (orange arrow). Blue arrows indicate a one-way targeting by trigger-piRNAs when responder-piRNAs do not match the initial trigger-producing transcript, thus excluding possibility of ping-pong amplification. The bars (color legend on the right) indicate percentages of trigger- and responder-piRNAs for each transcript (genome-unique reads). For AGAP011923, AGAP003266, and unk_RNA_1, responder-piRNAs are also trigger-piRNAs that target back the initial trigger-producing transcript. (B,D) Mapping of piRNAs to network transcripts. The entire 2R_lncs_14 and AGAP003387-RA mRNA are shown. Shown are all piRNAs mapping the transcripts without mismatch (in light green when genome-unique, in dark green when repeated twice in the genome) and the putative trigger-piRNAs (orange or blue, mismatched pairing). Trigger-piRNAs, all genome-unique, originate from AGAP011923 (orange) or from unk_RNA_1 (blue). Plus-oriented piRNAs are shown in the upper part, minus-oriented piRNAs in the lower part of each figure. The responder- and trail-piRNAs are indicated, respectively, by vertical and horizontal arrows above the zoom-in figure. Arrows below point to the trigger-piRNAs, together with the 5’-position of the corresponding responder-piRNAs. (C,E) Sequences of the responder-piRNAs (green) and trigger-piRNAs (orange, blue) for each trigger-responder pair. Trigger-piRNAs originate from AGAP011923 (orange) or unk_RNA_1 (blue). Transcript sequences surrounding the responder sequences are in black. The 5’-end position is indicated for each responder-piRNA. The inverted triangles show the slicer cleavage position facing nucleotides 10-11 of the trigger-piRNA. (B,C) Mapping of piRNAs to 2R_lncs_14. (D,E) Mapping of piRNAs to the AGAP003387-RA transcript.

A first set of trigger events involves AGAP011923 and AGAP003266 transcripts (Figure 4A). Trigger-piRNAs from AGAP011923 induce responder-piRNA slicing at 2 different positions within the 3’ UTR of AGAP003266 (reported in ref. (25)) (Supplemental Figure SFig3 C, D). In this particular case, the responder-piRNAs from AGAP003266 may target back the AGAP011923 transcript from which the corresponding trigger-piRNAs originate. This process results in trans ping-pong amplification (4) between the transcripts AGAP011923 and AGAP003266 (Figure 4A). It is noteworthy that the targeting piRNAs from AGAP011923 and AGAP003266 all have 1U and 10A (Table S9). Trail-piRNAs are found downstream of each of the three trigger/responder-piRNA peaks (two for AGAP003266 and one for AGAP011923), indicating that these trigger events induce effective phased piRNA biogenesis (Supplemental Figure SFig3 A, C). These data indicate that responder-piRNAs may actually become trigger-piRNAs targeting back the sequence that gave the initial trigger-piRNAs.

Trigger-piRNAs from AGAP011923 also induce phased piRNA biogenesis by slicing of the lncRNA 2R_lncs_14. The resulting trail-piRNAs from 2R_lncs_14 include 54.6 RPM piRNAs that are complementary to an unannotated RNA that we named unk_RNA_1 (“unknown RNA 1”) (Figure 4A). We aimed to establish whether these trail-piRNAs could act as trigger-piRNAs. We identified 7.3 RPM corresponding responder-piRNAs from unk_RNA_1 (Figure 4A, Supplemental Figure SFig3 E, F). To know whether there may be ping-pong between 2R_lncs_14 and unk_RNA_1, we looked whether unk_RNA_1 responders are complementary to the 2R_lncs_14 transcript. We found them complementary with 5 mismatches over 25 nt, but they do not match at the position of the initial trigger-piRNA within 2R_lncs_14. They match 113 nt downstream of it (Figure 4B). Thus, there is no ping-pong between these two transcripts. The targeting of the downstream position by the unk_RNA_1 piRNAs induces slicing of 2R_lncs_14 and leads to production of 1239.4 RPM responder-piRNA.

These results indicate that both trail-piRNAs and responder-piRNAs can become trigger-piRNAs for piRNA-guided slicing. This leads to an amplification of piRNA processing by a process of back-targeting, which differs from a ping-pong loop if the back-targeted sequence is different from the initial trigger-producing site.

In all cases, targeting by trigger-piRNAs very efficiently induces phased piRNA biogenesis. Relatively few trigger-piRNAs induce slicing of high amounts of responder-piRNAs, especially in the case of AGAP003387. Indeed, AGAP003387 is targeted by 63.9 RPM trigger-piRNAs from AGAP011923, and sliced into 86624 RPM responder-piRNAs and 1767 RPM trail-piRNAs, totalling 9% of all genome-mapping piRNAs in the *An. gambiae* ovary (21) (Figures 4A, D, E).

In conclusion, in *An. gambiae* ovaries, piRNAs from genes can be engaged in a regulation network involving mRNAs and lncRNAs from distant loci. The latter produce trigger-piRNAs inducing piRNA-guided slicing of other unrelated mRNAs or lncRNAs. Resulting responder-piRNAs and trail-piRNAs may themselves become trigger-piRNAs extending the network. Targeting by trigger-piRNAs leads to slicing and thus to the decay of the targeted transcripts.

### Genomic sequences producing piRNAs engaged in networks are repeated and conserved from insects to mammals

To find out whether similar piRNA-networks exist in other species, we aligned sequences of the five network transcripts from *An. gambiae* to 38 disease vector genomes (mosquitoes, flies, arachnidae, snail), to *Drosophila melanogaster* and to mammal genomes (human, mouse, rat) by BLASTN.

We found that the five network transcripts were conserved either over their whole length (2R_lncs_14, AGAP003387, AGAP003266) or at least over 80% of their length (AGAP011923, unk_RNA_1) in six species of the *An. gambiae* complex (Figure 5A-E, Supplemental Figure SFig4, Table S10). In the other species, short motifs encompassing approximately 40 nucleotides were conserved across multiple genomes. Strikingly, the majority of these motifs correspond to the sequences encompassing the trigger-piRNA annealing site and the responder-piRNA sequences, which we identified as snetDNA for “small-RNA-network DNA”. These sequences were found conserved not only in the 21 *Anopheles* species but also in the other 22 analysed species. Moreover, the snetDNA sequences were highly repeated. In *Culex quinquefasciatus*, the snetDNA positioned between 996-1032 in 2R_lncs_14 was found repeated more than 200 times (e-value <1, conservation over >30 nt), 193 copies being clustered together in the same locus of 10.5 kb, organized in tandem repeats of 54-nt length (Table S11). We also found that many of the BLASTN matches hit annotated transcripts (Figure 5F, Table S12). Moreover, as annotated transcripts do not include all non-coding RNAs, the number of hits in transcripts is probably largely underestimated.

**Figure 5.**
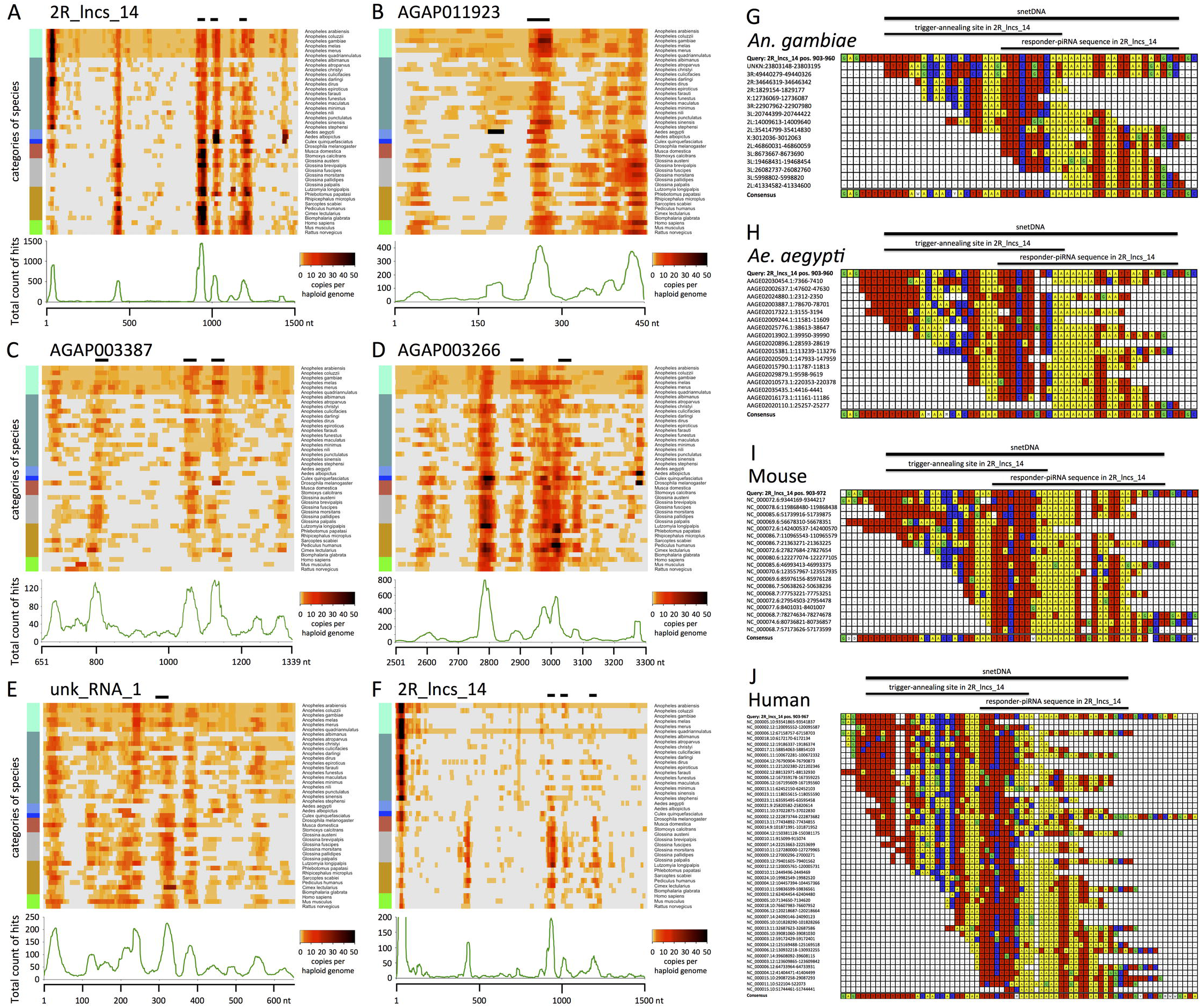
BLASTN results reveal short conserved repeats, the snetDNAs. (A-F) Heat maps illustrating BLASTN results, allowing a maximal e-value of 10.0, for network transcripts 2R_lncs_14 (A), AGAP011923-RA (B), AGAP003387-RA (C), AGAP003266 3’-UTR (D), and unk_RNA_1 (E) aligned to genomes (A-E) and for 2R_lncs_14 aligned to annotated transcripts (F) of 43 different species (detailed BLASTN results in Table S10). The counts of hits are plotted as sliding windows, corresponding each to one similar sequence, along the transcript for each individual species. Zero hit is in light grey, one single hit in the lightest orange. Light orange along the whole transcript indicates existence of a genomic sequence, which is similar all over the transcript length (see also Table S10). This situation is observed in the six species of the *An. gambiae* complex. On the left, a color code for different categories of species is presented. Mosquitoes are in different blue colors: *An. gambiae* complex, light blue; other *Anopheles*, cadet blue; *Aedes*, cornflower blue; and *Culex, blue*; coral: flies; gray: tsetse; green: mammals; dark golden: other species, including snail, body louse, bed bugs, tick, mite, and sand flies. Horizontal bars above the heat map indicate the trigger-responder sites (spanning over the trigger-annealing site and the corresponding responder-piRNAs). Below each heat map is a graphical representation of the totalized counts of hits for all analysed species mapped for each position of the transcript. The detailed BLASTN results with the aligned sequences can be found in Tables S10 (for A-E) and S12 (for F). (G-J) Multiple sequence alignment and consensus of *An. gambiae* (G), *Ae. aegypti* (H), mouse (I) and human (J) snetDNAs matching to 2R_lncs_14 positions 903-960 (“Query”, top sequence in each panel). The resulting consensus sequences are shown in the last line of each panel. Color code: Adenine in yellow, cytosine in blue, guanine in green and thymine in red. Additional alignments and consensus sequences of snetDNAs can be found in Supplementary SI-MSA2.

We determined consensus sequences of the best-conserved snetDNAs in *An. gambiae, Ae. aegypti*, mouse and Human and found them highly similar in these evolutionary very distant species of mosquitoes and mammals (Figure 5G-J, Supplementary SI-MSA2). In all four species, the snetDNAs homologous to 2R_lncs_14 contain a conserved central motif CACTNAAATTTCTTTN(A)_3-6_TTAA. The AGAP011923-RA related snetDNAs have CANANTTATCAGAAATTTAAGTG in their center. In each case the 5’-end of the putative responder-piRNA corresponds to a T-rich sequence, TTTCTTT for 2R_lncs_14-related snetDNAs and TTATCAG for AGAP011923-RA-related snetDNAs, with a highly conserved T corresponding to the 5’ 1U of the putative responder-piRNA. Our data suggest that the piRNA mediated regulation network of transcripts based on repeated snetDNAs is largely conserved in the animal kingdom.

Although well investigated nowadays, lncRNA function remains elusive. It is very interesting to observe that the lncRNA 2R_lncs_14 displays the most conserved sequence of the network. Actually, it is conserved over its whole length in the six species of the *An. gambiae* complex, and its snetDNAs are conserved in all 43 analysed species (Figure 5A, Supplemental Figure SFig4A) with sequence identity ranging from 81% to 97.5% over 35-45 nucleotides (Table S10). Sequences homologous to 2R_lncs_14 snetDNAs are often found clustered together in the same locus with three to nine snetDNAs identified by BLASTN distributed over less than 4 kb (Table S11). This configuration resembles the one of 2R_lncs_14 (Figure 4B). In multiple cases, the conserved sequence at positions 1408-1440 of 2R_lncs_14 was also present in these clustered snetDNA loci, suggesting that it is another conserved snetDNA (Figures 4B and 5A, Table S11). We checked whether this region of 2R_lncs_14 may be targeted by putative trigger-piRNAs and identified such targeting piRNAs as well as the corresponding responder-piRNAs located at positions 1407-1435 (Figure 4B, Table S9). Eleven loci with clustered snetDNAs were identified in *Ae. aegypti*, ten in *Aedes albopictus*. At least one such locus with 2 to 9 snetDNAs could be identified in each of the analysed *Anopheles, Aedes*, and *Culex* species; with the exception of *An. maculatus* in which only non-clustered snetDNAs could be identified by BLASTN. We propose that these loci with clustered snetDNAs are orthologs of 2R_lncs_14.

### SnetDNAs from the *Aedes aegypti* mosquito produce piRNAs and establish networks

An important question is whether the conserved snetDNAs produce piRNAs and establish “snetDNA-networks” in mosquito species other than *An. gambiae*. To answer this question, we analysed small RNAs that had been sequenced from *Ae. aegypti* adults (26).

In *Ae. aegypti*, we found piRNAs similar to AGAP011923 trigger-piRNAs (Supplemental Figure SFig5). They originate from two distant loci, the annotated gene AAEL017228 (AAEL027227 in the AaegL5 assembly) and a region downstream of AAEL009512, that may be in fact its extended 3’-UTR (Figure 6A). These putative trigger-piRNAs differ by only 2 nucleotides and might thus target the same transcripts. Notably, AAEL017228-RA is the transcript, which has the most mapping piRNAs among all annotated transcripts in *Ae. aegypti*. It also maps >3 times more piRNAs than the AAEL017385 gene which contains tapiR1 and tapiR2, two highly abundant piRNAs in *Ae. aegypti*, conserved for approximately 200 million years, with biological function in control of gene expression and embryonic development (27). Interestingly, the putative AAEL009512 3’-UTR has four identical piRNA-producing sequences (Supplemental Figure SFig6). These sequences are unique to this region, i.e., they do not match other genomic regions and they are clustered together over less than 4 kb (Figure 6B), as is observed for snetDNAs within 2R_lncs_14 and its putative orthologs (Figure 4B, Table S11) or for AGAP003387 in *An. gambiae* (Figure 4D).

**Figure 6.**
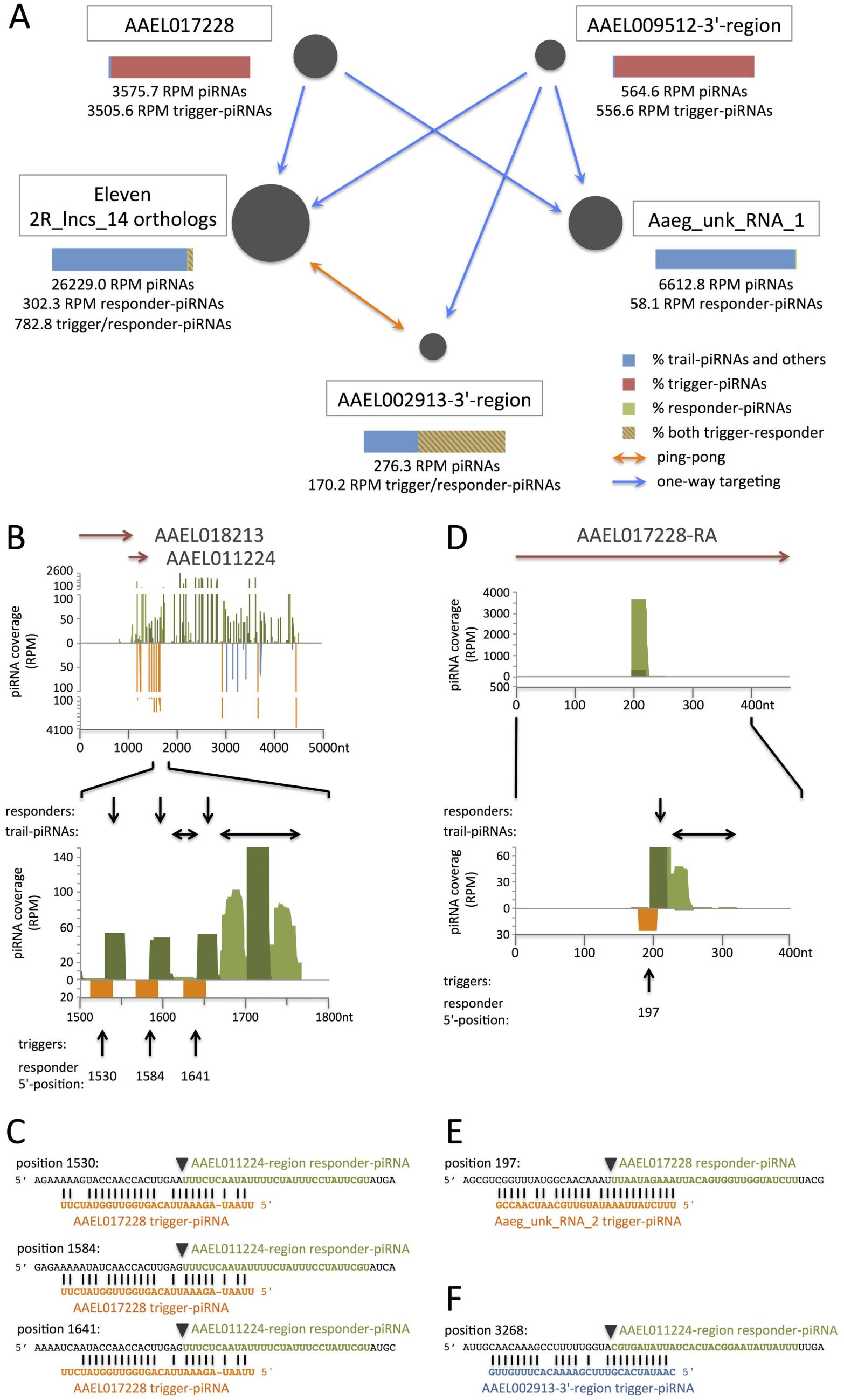
SnetDNA-network involving mRNAs and lncRNAs in *Ae. aegypti*. (A) Schematic representation of the network. Trigger-piRNAs from AAEL017228 and the AAEL009512 3’-region target transcripts from 2R_lncs_14 orthologs, Aaeg_unk_RNA_1 (non-annotated) and the AAEL002913 3’-region. Resulting responder-piRNAs from the AAEL002913 3’-region target back 2R_lncs_14 orthologs leading to trans ping-pong amplification. Arrows indicate one-way targeting (blue) or trans ping-pong (orange). Bars (color legend on the right) indicate percentages of trigger- and responder-piRNAs for each transcript (all mapping reads). Trigger/responder-piRNAs may function both as trigger- or responder-piRNAs. (B,C,F) Mapping of piRNAs to the AAEL011224 region (AaegL3_supercont1.555:368337-373336[-]). This region is the 2R_lncs_14 ortholog with the highest number of clustered snetDNAs and producing the highest amount of genome-unique piRNAs in *Ae. aegypti*. Shown trigger-piRNAs, all genome-unique, originate from AAEL017228 (orange) and from the AAEL002913 3’-region (blue). (D,E) Mapping of piRNAs to the AAEL017228-RA transcript. Trigger-piRNAs, all genome-unique, originate from Aaeg_unk_RNA_2 (orange). (B,D) Mapping of piRNAs to network transcripts. Shown are all piRNAs mapping the transcripts without mismatch (in light green when genome-unique, in dark green when repeated in the genome) and the putative trigger-piRNAs (orange or blue, mismatched pairing). Arrows in the upper part indicate the annotated genes in the AAEL011224 region. The entire AAEL017228-RA mRNA is shown. Plus-oriented piRNAs are shown in the upper part, minus-oriented piRNAs in the lower part of each figure. The responder- and trail-piRNAs are indicated, respectively, by vertical and horizontal arrows above the zoom-in figure. Arrows below point to the trigger-piRNAs, together with the 5’-position of the corresponding responder-piRNAs. (C,E,F) Sequences of the responder-piRNAs (green) and trigger-piRNAs (orange, blue) for each trigger-responder pair. Transcript sequences surrounding the responder sequences are in black. The 5’-end position is indicated for each responder-piRNA. The inverted triangles show the slicer cleavage position facing nucleotides 10-11 of the trigger-piRNA.

We also identified piRNAs similar to *An. gambiae* 2R_lncs_14 responder-piRNAs in *Ae. aegypti*. They have 10-nt 5’-overlap to the putative trigger-piRNAs from AAEL017228 and AAEL009512 and originate from previously identified 2R_lncs_14 orthologs (see above). Actually, the eleven 2R_lncs_14 orthologs identified in *Ae. aegypti* (Table S11) produce a large amount of piRNAs, 2.6% of all piRNAs. Each of these orthologs also gives genome-unique piRNAs in addition to non-unique piRNAs, attesting that they all produce piRNAs. The *Ae. aegypti* snetDNA-network is shown in Figure 6A, alignments of trigger- and responder-piRNAs to the implicated transcripts are shown in Figure 6B-F and Supplemental Figure SFig6. We performed BLASTN of putative trigger- and responder-piRNA sequences against the *Ae. aegypti* genome. We found that trigger-piRNAs from AAEL017228 are genome-unique while trigger-piRNAs from the AAEL009512 3’-region map to 4 sites clustered together and are unique to this region. On the other hand, the piRNA sequences similar to *An. gambiae* 2R_lncs_14 responder-piRNAs were found highly repeated with a total of 74 genomic sites. All except two are localized in the eleven identified 2R_lncs_14 orthologs (Tables S11 and S13). The *Ae. aegypti* 2R_lncs_14 ortholog with 9 snetDNAs identified by BLASTN produces the highest amount of genome-unique piRNAs, 2824.9 RPM. Interestingly, it overlaps two annotated protein-coding genes in the same orientation, AAEL011224 and AAEL018213. When aligned to piRNAs allowing up to 6 mismatches, we identified a total of 17 snetDNAs, targeted by trigger-piRNAs from AAEL017228, AAEL009512 and AAEL002913 3’-regions, in this 2R_lncs_14 ortholog spanning over 4.6 kb (Figure 6B, C). Responder-piRNAs from the AAEL002913 3’-region (annotated as AAEL025618-RA transcript in AaegL5) may target back *Ae. aegypti* 2R_lncs_14 orthologs indicating that in *Ae. aegypti*, as in *An. gambiae*, responder-piRNAs may become trigger-piRNAs (Figure 6A, B, F, Supplemental SFig6 E, F).

### The characteristics of snetDNAs in mouse and human suggest that snetDNAs derive from ancient transposable elements

We analysed small RNAs that had been sequenced from mouse fetal and adult testes (28, 29). We found piRNAs that are similar to *An. gambiae* and *Ae. aegypti* network piRNAs (Supplemental Figure SFig5). These piRNAs match putative mouse snetDNAs determined by BLASTN (in Figure 5A-E, Table S10) either without any mismatch indicating production of piRNAs from these snetDNAs (fetal mouse piRNAs in SFig5), or when allowing up to 4 mismatches indicating possible targeting by piRNAs (adult mouse piRNAs in SFig5).

We analysed the localisation of mouse and human snetDNAs, found by BLASTN analyses (in Figure 5A-E, Table S10), with respect to annotated genes and TE-related sequences. Mouse snetDNAs significantly overlap with genes, long intergenic noncoding RNAs (lincRNAs), LINEs and DNA repeats. Human snetDNAs overlap with genes, protein-coding genes, introns and DNA repeats (Table S14). In mouse and human, the snetDNA containing TEs belong to the haT transposons of the Charlie sub-family in mouse and Charlie/MER and Tigger sub-families in human. haT transposons represent ancient, inactive transposons (30). The significant overlap of mouse and human snetDNAs with TEs suggests that snetDNAs in fact derive from TEs.

Genes containing snetDNAs are mostly expressed in multiple or all tissues but some of these genes are specifically expressed in male gonadal tissue or brain. In fact, from 54 snetDNA containing genes in human, 21 are tissue-specific by TissueEnrich (Table S15). From the 21 tissue-specific genes, 7 are specifically expressed in testes, 8 in cerebral cortex. In mouse, out of 25 snetDNA containing genes, 8 are tissue-specific. Two of the latter are specifically expressed in testes, six in brain tissues. Tissue-specific expression of snetDNA containing genes in testes and brain is remarkable because these tissues express piRNAs (14, 31–33) making piRNA processing from the transcripts of these genes possible.

Our data show that snetDNAs are evolutionarily conserved in quite distant species, from insects to mammals, and probably derive from TEs. SnetDNAs may produce piRNAs and/or be targeted by piRNAs and thus be at the origin of regulation networks by RNA degradation as shown here for *An. gambiae* and *Ae. aegypti*.

## DISCUSSION

Here we extended knowledge on biogenesis of piRNA from genes and lncRNAs in *An. gambiae*. These piRNAs originate not only from ping-pong amplification between genic transcripts but also from phased biogenesis initiated by trigger-piRNAs originating from distant loci. The majority of piRNAs from genes and lncRNAs in *An. gambiae* show no similarity with known TEs. They constitute a basis for networks of concerted gene regulation, each one involving an ensemble of mRNAs that contain specific repeats, which are at the origin of piRNA production. Two different types of networks are observed: (a) “ping-pong-networks” grouping transcripts containing the same repeated sequences, 130 to 1,130 base pairs long, producing piRNAs which integrate the ping-pong amplification cycle and (b) “snetDNA-networks” grouping transcripts containing snetDNAs, short repeats of size between 25 and 42 nucleotides, that produce trigger- and responder-piRNAs which are at the origin of concerted phased piRNA biogenesis. In the latter network, transcripts producing trigger-piRNAs target other complementary transcripts and induce their slicing to responder- and trail-piRNAs. Responder- and trail-piRNAs may then become trigger-piRNAs targeting other complementary transcripts. If the responder-piRNAs are complementary to the initial trigger-producing transcript, they may target back the latter and induce piRNA amplification between these two transcripts either by trans ping-pong or phased piRNA biogenesis.

For the “ping-pong-networks”, we observed piRNA ping-pong amplification between transcripts from distant loci having the same repeats. In addition, for 11 other mRNAs, we identified a novel “intrinsic ping-pong amplification”, which is achieved with genome-unique piRNAs originating from the same single gene. Until now, ping-pong amplification had been observed between piRNAs from different loci, i.e. piRNA clusters on one hand and functional TEs on the other hand. Here we identified 11 transcripts from 8 genes, which produce both sense and antisense ping-pong partners by their own, without need of targeting piRNAs from other loci. An open question now is how these genes become autonomous piRNA producing loci.

In snetDNA-networks, we observed that relatively few trigger-piRNAs may induce slicing of high amounts of responder-piRNAs, especially in the case of AGAP003387. This suggests that, as observed in *C. elegans*, piRNA concentration does not limit targeting but probably correlates better with binding energy than with piRNA abundance (34).

Our data give insights into the mechanisms of target slicing. Ago3-bound piRNAs in *Drosophila* have enriched 10A while Aub-bound piRNAs are enriched for 1U only (1). In the networks described above, 96.5% of the trigger-piRNAs have both 1U and 10A signatures, whereas 98.4% of the responder-piRNAs have 1U but no 10A (Table S9). Assuming that 1U and 10A signatures of Ago3- and Aub-bound piRNAs have the same specificity in *Drosophila* and *Anopheles*, even if this still has to be tested, we hypothesize that trigger-piRNAs of the *An. gambiae* snetDNA-network are essentially Ago3-bound and responder-piRNAs Aub-bound.

In *Drosophila*, it is known that responder-piRNAs are essentially bound onto Aub with a minority of them bound onto Ago3 (5). Han *et al*. (6) observed that RNAs cut by Ago3 produce phased Aub-bound piRNAs, but RNAs cut by Aub do not produce phased Ago3-bound piRNAs. In the *An. gambiae* snetDNA-network, trigger-responder pairs are not necessarily followed by phased trail-piRNAs (AGAP000203 and AGAP013373 in Figure 3, 2R_lncs_14 responder-positions 929 and 1122 in Figure 4B). Also, responder- and trail-piRNAs of the network may become trigger-piRNAs. This raises the question to which proteins these piRNAs may be bound. Our observations suggest that in *An. gambiae*, trigger- and responder-piRNAs may be bound either to Aub or Ago3 orthologs, like in *Drosophila*, and that slicing of phased trail-piRNAs may depend on the protein that charged the trigger-piRNAs.

In the *Drosophila melanogaster* germline, phased piRNA biogenesis depends on the targeting of transcripts by trigger-piRNAs. Pairing between the trigger-piRNAs and the targets supports some mismatches (5). We assumed that this is the same in the *An. gambiae* germline. In mouse, some authors suggested that slicing by PIWI argonautes, like Aub and Ago3, requires perfect matching between the transcript and the trigger-piRNA at positions 2-11, the “seed” (35), while others observed slicing of the targeted transcript even with 3 mismatches over positions 1-20 of the trigger-piRNA (36). It is not known whether the strict pairing within a seed sequence is required for PIWI slicing in *An. gambiae*. In the *An. gambiae* ovary, we observed typical genome-unique responder- and phased trail-piRNAs originating from the network transcripts and identified corresponding putative trigger-piRNAs. From 12 different trigger events implicated in the snetDNA network in *An. gambiae*, only 5 show a perfect match over the whole putative seed, positions 2-11 (SFig3). For the rest of the trigger-piRNAs, perfect matching is observed in all cases around the slicing position 10-11, while 1 to 3 mismatches are observed within the putative seed. We hypothesize that the strict pairing within the seed is not necessarily required for the slicing by the PIWI-protein(s) involved in the network systems of *An. gambiae. An. gambiae* has three PIWI class proteins, Ago3 (AGAP008862), AGAP011204 and AGAP009509 (37). We propose that at least one of these proteins does not need necessarily strict matching within the whole 2-11 seed for slicing activity, but uses a smaller seed or allows some relaxed matching within the seed. We also observed that, even when base pairing starts at position 2 or 3, the slicing of the target always occurs between the nucleotides 10 and 11 relative to the piRNA 5’ end. This is consistent with what has been observed in mouse testes (36).

Our present data suggest that *An. gambiae* mRNAs and lncRNAs are processed into piRNAs that are able to target other transcripts and direct them into the piRNA pathway, thus creating piRNA-mediated regulation networks. Such networks are based on the presence, within the network transcripts, of short repeated sequences, the snetDNAs, which are conserved from insects to mammals, notably in the 38 tested species of disease-vectors (mosquitoes, tsetse flies, sand fly, tick, mite, body louse, bed bug, snail). Due to the production of piRNAs from snetDNAs, networks of interdependent regulation are established between different genes and lncRNAs and may control their transcripts’ stability/slicing within specific tissues where piRNA pathway and ping-pong amplification are known to be active. It is important to note that such regulatory pathways may function with any repeated sequences that produce piRNAs. Still, the conservation of specific snetDNA consensus sequences indicates an evolutionary and functional advantage of these specific sequences.

Targeting of transcripts by trigger-piRNAs and subsequent slicing may interfere with the expression of the targeted transcripts. AGAP003387 transcripts are part of the *An. gambiae* snetDNA-network evidenced here. Remarkably, it has been shown that the level of, AGAP003387 transcripts was increased in piRNA-pathway impaired females, in which PIWI class proteins Ago3, AGAP011204 and AGAP009509 were knocked down. These results obtained by knocking-down independently the three different PIWI class proteins show that AGAP003387 is regulated by the piRNA pathway (23). Our data now explain why this is the case: Knock-down of these PIWI-class proteins impairs piRNA-targeting of the AGAP003387 transcript and the subsequent slicing, thus leading to higher AGAP003387 transcript levels. Taken together, these data provide evidence of the functionality of the *An. gambiae* snetDNA-network reported here.

In mouse testes, piRNA-guided cleavage of mRNAs has been demonstrated by Zhang et al. (36). The functional impact of piRNA-guided mRNA cleavage is further evidenced as these authors report that potential piRNA-targeted mRNAs directly interact with Miwi, the mouse Piwi protein, and show higher expression levels in the testes of Miwi catalytic mutant mice compared to wildtype. Gainetdinov et al. (38) report that the mechanism of piRNA biogenesis possibly is the same in most animals. Together with our data, these observations suggest that similar mechanisms of piRNA-mediated mRNA regulation possibly exist in most animal species.

In this study, we detail networks in *An. gambiae* and in *Ae. aegypti*, but it may be anticipated that many more will be discovered through further analyses. As snetDNAs, very short repeated sequences, are sufficient to establish trigger/responder-piRNA-based regulatory networks in *An. gambiae*, it is likely that other repeats, TE-related or not, inserted within genes may assure additional regulatory networks based on piRNA biogenesis. It is conceivable that target genes with such repeats are repressed in normal environmental conditions because their transcripts are sliced for piRNA production. Their repression may vary in a concerted manner if epigenetic events affect the piRNA pathway, contributing to a possible rapid plasticity in their expression levels.

## METHODS

### Data Access

Small RNA libraries were SRX966734, for *An. gambiae* ovaries (21), available at NCBI SRA database, GSM830467 for *Ae. aegypti* adults (26), GSM1318059 for fetal mouse testes (28) and GSM822760 and GSM822762 for adult mouse testes (29), these datasets being available at NCBI GEO DataSets.

### Mapping small RNA reads

Reads were mapped against AgamP4.2, AaegL3, genome assemblies for *An. gambiae* and *Ae. aegypti* respectively, allowing zero mismatch, using sRNAPipe (39). Small RNAs of 24-29 nt length were considered as putative piRNAs. They were mapped against transcripts or TEs using scripts and pipelines based on BWA (40), Bowtie2 (41), or NucBase (42). RPM (reads per million) for each sequence were calculated as the number of reads per million genome-mapped reads allowing no mismatch. Multi-mapping reads were randomly assigned to the mapping sequences. Transcript-mapping reads were identified by alignment to protein-coding transcripts from genome assemblies AgamP4.2 (*An. gambiae*) and AaegL3.2 (*Ae. aegypti*), available at VectorBase (43), and to lncRNAs (44). TE-matching reads were identified by alignment to all TE sequences for the subphylum Hexapoda that are available on RepBase (http://www.girinst.org) (45), allowing up to 3 mismatches.

*Bona fide* reads were defined as small RNAs which do not map microRNAs (miRNAs), ribosomal RNAs (rRNAs), small nuclear RNAs (snRNAs) or transfer RNAs (tRNAs). “Collapsed” counts were determined from collapsed libraries where all identical reads were reduced to one single read. “Non-collapsed” counts originate from the initial sequencing libraries. “Multiplicity” was calculated as the ratio of the non-collapsed count of piRNAs over the collapsed count (Table S1).

Tablet was used for visual exploration of piRNA mapping (46).

### Analyses of ping-pong signature

Ping-pong signature was analysed essentially as in (47). The putative piRNAs, 24-29 nt in length, were mapped to the transcripts as above. Their 5’-positions on the mapped sequences were determined from the resulting sam files by a dedicated script, available upon request. The 5’-overlaps between sense-oriented piRNAs and antisense-oriented piRNAs were calculated for all opposite piRNAs with a maximal distance of 24 nt between their respective 5’-positions. This maximal overlap of 24 nt was chosen according to the size of the considered piRNAs, 24-29 nt, and corresponds to the maximal overlap which is possible for all considered piRNAs to avoid the bias observed above this limit. We concluded for a significant 10-nt 5’-overlap when its z-score was >1.96, within the window of 5’-overlaps from 1 to 24 nt (including 10-nt 5’-overlaps), at condition that the total number of overlapping piRNA pairs within this window was at least 30 (for any overlap length).

### Repeat Masking

For repeat-masking we used CENSOR at http://www.girinst.org and all insect TEs available at Repbase (48).

### BLASTN analyses

BLASTN alignments were done at VectorBase (43) for disease vectors (https://www.vectorbase.org/blast, E=1 or E=10, Word size: 11, Complexity Masking: Off) or by command-line blastn, with settings matching those of VectorBase, for *Drosophila*, human, mouse and rat (/opt/blast-2.4.0/bin/blastn -db genome -query transcripts.fa -task “blastn-short” -evalue 10 –reward 2 -penalty -3 -outfmt “7 qacc qstart qend sacc sstart send sstrand evalue bitscore length pident mismatch gapopen qseq sseq” -out blastn-results.txt), these genomes being unavailable at VectorBase. In the case of command-line blastn, for the figures with alignments respecting a maximal E-value of 1, the alignments with an E-value >1 were dropped. Redundancy within BLASTN results was suppressed before making heat maps: For overlapping, nested alignments, only the longest alignment was retained.

### SnetDNA consensus sequences

To build snetDNA consensus sequences, the genomic sequences that aligned to 2R_lncs_14 and AGAP011923 snetDNAs and flanking sequences (BLASTN E=10 as described in “BLASTN analyses”) were extracted from the respective genomes (*An. gambiae* AgamP4, *Ae*.*aegypti* AaegL3, mouse GRCm38.p5 and human GRCh38.p8) and aligned using “Muscle” and “ClustalW” multiple sequence alignment tools included in MacVector 17.0.5.

### Bio-informatics

Barplots, scatter plots, heatmaps, Euler diagram, Principal component analysis, and correlograms were made using “R” (R Core Team, 2017. “R: A language and environment for statistical computing.” RFoundation for Statistical Computing, Vienna, Austria. URL https://www.R-project.org/). Principal component analysis was proceeded using “FactoMineR” (49) and “factoextra” (http://www.sthda.com/english/rpkgs/factoextra). Euler diagram was made with eulerr (http://eulerr.co, https://CRAN.R-project.org/package=eulerr) Heatmaps were made using “R” package gplots (https://CRAN.R-project.org/package=gplots). Correlograms were made using “R” packages “corrplot” (https://CRAN.R-project.org/package=corrplot) and “Hmisc” (https://CRAN.R-project.org/package=Hmisc). “ratioBsBas” and “LOGratioBsBas” for correlograms and PCA were computed as follows: “Bs” being the count of all *bona fide* sense reads in RPM, “Bas” being the count of all *bona fide* antisense reads in RPM, “ratioBsBas” was computed as Bs/Bas, or as (Bs+0.06)/Bas when Bs=0, or as Bs/(Bas+0.06) when Bas=0; “LOGratioBsBas” is log(ratioBsBas). Coverage of transcripts by piRNAs was analysed using bedtools genomecov (50).

### snetDNAs and genomic annotations

We extracted gene annotations from GENCODE basic annotations v28 for the human genome (GRCh38.p12), as well as GENCODE basic annotations v19 for the mouse genome (GRCm38.p6) (51) downloaded from GENCODE ftp site. For each gene, we selected its longest transcript to define its exons/introns structure. In addition, we extracted repeat annotations predicted with RepeatMasker (Smit, AFA, Hubley, R & Green, P. RepeatMasker Open-4.0. 2013-2015, http://www.repeatmasker.org) for human and mouse and available from the UCSC portal (52). To better characterize the genomic locations of snetDNAs, we tested the overlap between snetDNAs and several genomic features, using the GenometriCorr package (53). The GenometriCorr package implements spatial tests of independence between two sets of genomic intervals. We used the Jaccard test to compare observed overlaps to a null distribution based on permutations of the snetDNAs intervals across the genome.

### Expression of genes containing snetDNAs

We investigated expression enrichment towards particular tissues amongst protein coding genes containing a snetDNA, using the TissueEnrich package (54). TissueEnrich defines tissue-specific genes using RNA-Seq data in 35 human tissues from the Human Protein Atlas (55) and in 17 mouse tissues from the Mouse ENCODE Dataset (56). The package then provides a test for significant enrichment of tissue-specific genes towards particular tissues, using a hypergeometric test with the Benjamini & Hochberg correction for multiple testing.

The scripts are available upon request.

## Supporting information

Table S13 and multiple sequence alignments SI-MSA1

multiple sequence alignments SI-MSA2

## ACCESSION NUMBERS

All small RNA sequencing data used here are available at NCBI SRA database or NCBI GEO DataSets. They have been published by us or by others and are publicly available. The accession numbers are given in the Methods section together with the corresponding reference.

## SUPPLEMENTARY DATA

Tables S1 to S12 and S14 and S15 can be found in file “Supplementary-1.xlsx”, supplementary figures, Table S13 and multiple sequence alignments SI-MSA1 in file “Supplementary-2.pdf” and multiple sequence alignments SI-MSA2 in file “Supplementary-3.pdf”.

## ACKNOWLEDGEMENTS

We thank Romain Pogorelcnik for creating bioinformatic tools and pipelines, Pierre Pouchin, Yoan Renaud, Franck Court and Nadia Goué for active support concerning bioinformatics and helpful discussions.

## AUTHOR CONTRIBUTIONS

S.J., E.B. and C.V. designed the study. S.J. designed and developed the concept, analysed the data, conducted all bioinformatics and statistics analyses, except the study of snetDNA localization in mouse and human genomes, which was done by E.P. and H.R.-C.. S.J. and E.B. were in charge of all bioinformatic scripts and pipelines. S.J. wrote the paper. I.V.S., E.B. and C.V. gave conceptual advice, commented on and edited the manuscript. All authors discussed the results and implications at all stages.

## REFERENCES

1. Brennecke, J., Aravin, A.A., Stark, A., Dus, M., Kellis, M., Sachidanandam, R. and Hannon, G.J. (2007) Discrete Small RNA-Generating Loci as Master Regulators of Transposon Activity in Drosophila. Cell, 128, 1089–1103.

2. Gunawardane, L.S., Saito, K., Nishida, K.M., Miyoshi, K., Kawamura, Y., Nagami, T., Siomi, H. and Siomi, M.C. (2007) A slicer-mediated mechanism for repeat-associated siRNA 5’ end formation in Drosophila. Science, 315, 1587–1590.

3. Ozata, D.M., Gainetdinov, I., Zoch, A., O’Carroll, D. and Zamore, P.D. (2019) PIWI-interacting RNAs: small RNAs with big functions. Nat. Rev. Genet., 20, 89–108.

4. Wang, W., Yoshikawa, M., Han, B.W., Izumi, N., Tomari, Y., Weng, Z. and Zamore, P.D. (2014) The Initial Uridine of Primary piRNAs Does Not Create the Tenth Adenine that Is the Hallmark of Secondary piRNAs. Mol. Cell, 56, 708–716.

5. Mohn, F., Handler, D. and Brennecke, J. (2015) piRNA-guided slicing specifies transcripts for Zucchini-dependent, phased piRNA biogenesis. Science, 348, 812–817.

6. Han, B.W., Wang, W., Li, C., Weng, Z. and Zamore, P.D. (2015) piRNA-guided transposon cleavage initiates Zucchini-dependent, phased piRNA production. Science, 348, 817–821.

7. Gan, H., Lin, X., Zhang, Z., Zhang, W., Liao, S., Wang, L. and Han, C. (2011) piRNA profiling during specific stages of mouse spermatogenesis. RNA, 17, 1191–1203.

8. Sarkar, A., Volff, J.-N. and Vaury, C. (2017) piRNAs and their diverse roles: a transposable element-driven tactic for gene regulation? FASEB J., 31, 436–446.

9. Yamtich, J., Heo, S.-J., Dhahbi, J., Martin, D.I. and Boffelli, D. (2015) piRNA-like small RNAs mark extended 3’UTRs present in germ and somatic cells. BMC Genomics, 16, 462.

10. Robine, N., Lau, N.C., Balla, S., Jin, Z., Okamura, K., Kuramochi-Miyagawa, S., Blower, M.D. and Lai, E.C. (2009) A Broadly Conserved Pathway Generates 3′UTR-Directed Primary piRNAs. Curr. Biol., 19, 2066–2076.

11. Bouhouche, K., Gout, J.-F., Kapusta, A., Bétermier, M. and Meyer, E. (2011) Functional specialization of Piwi proteins in Paramecium tetraurelia from post-transcriptional gene silencing to genome remodelling. Nucleic Acids Res., 39, 4249–4264.

12. Lee, H.-C., Gu, W., Shirayama, M., Youngman, E., Conte, D. and Mello, C.C. (2012) C. elegans piRNAs Mediate the Genome-wide Surveillance of Germline Transcripts. Cell, 150, 78–87.

13. Roovers, E.F., Rosenkranz, D., Mahdipour, M., Han, C.-T., He, N., Chuva de Sousa Lopes, S.M., van der Westerlaken, L.A.J., Zischler, H., Butter, F., Roelen, B.A.J., et al. (2015) Piwi Proteins and piRNAs in Mammalian Oocytes and Early Embryos. Cell Rep., 10, 2069–2082.

14. Rajasethupathy, P., Antonov, I., Sheridan, R., Frey, S., Sander, C., Tuschl, T. and Kandel, E.R. (2012) A Role for Neuronal piRNAs in the Epigenetic Control of Memory-Related Synaptic Plasticity. Cell, 149, 693–707.

15. Saito, K., Inagaki, S., Mituyama, T., Kawamura, Y., Ono, Y., Sakota, E., Kotani, H., Asai, K., Siomi, H. and Siomi, M.C. (2009) A regulatory circuit for piwi by the large Maf gene traffic jam in Drosophila. Nature, 461, 1296–1299.

16. Barckmann, B., Pierson, S., Dufourt, J., Papin, C., Armenise, C., Port, F., Grentzinger, T., Chambeyron, S., Baronian, G., Desvignes, J.-P., et al. (2015) Aubergine iCLIP Reveals piRNA-Dependent Decay of mRNAs Involved in Germ Cell Development in the Early Embryo. Cell Rep., 12, 1205–1216.

17. Rouget, C., Papin, C., Boureux, A., Meunier, A.-C., Franco, B., Robine, N., Lai, E.C., Pelisson, A. and Simonelig, M. (2010) Maternal mRNA deadenylation and decay by the piRNA pathway in the early Drosophila embryo. Nature, 467, 1128–1132.

18. Watanabe, T., Cheng, E., Zhong, M. and Lin, H. (2015) Retrotransposons and pseudogenes regulate mRNAs and lncRNAs via the piRNA pathway in the germline. Genome Res., 25, 368–380.

19. Ku, H.-Y. and Lin, H. (2014) PIWI proteins and their interactors in piRNA biogenesis, germline development and gene expression. Natl. Sci. Rev., 1, 205–218.

20. Rojas-Ríos, P. and Simonelig, M. (2018) piRNAs and PIWI proteins: regulators of gene expression in development and stem cells. Dev. Camb. Engl., 145.

21. George, P., Jensen, S., Pogorelcnik, R., Lee, J., Xing, Y., Brasset, E., Vaury, C. and Sharakhov, I.V. (2015) Increased production of piRNAs from euchromatic clusters and genes in Anopheles gambiae compared with Drosophila melanogaster. Epigenetics Chromatin, 8, 50.

22. Castellano, L., Rizzi, E., Krell, J., Di Cristina, M., Galizi, R., Mori, A., Tam, J., De Bellis, G., Stebbing, J., Crisanti, A., et al. (2015) The germline of the malaria mosquito produces abundant miRNAs, endo-siRNAs, piRNAs and 29-nt small RNAs. BMC Genomics, 16, 100.

23. Biryukova, I. and Ye, T. (2015) Endogenous siRNAs and piRNAs derived from transposable elements and genes in the malaria vector mosquito Anopheles gambiae. BMC Genomics, 16, 278.

24. Abdi, H. and Williams, L.J. (2010) Principal component analysis. Wiley Interdiscip. Rev. Comput. Stat., 2, 433–459.

25. Gomez-Diaz, E., Rivero, A., Chandre, F. and Corces, V.G. (2014) Insights into the epigenomic landscape of the human malaria vector Anopheles gambiae. Front. Genet., 5, 277.

26. Arensburger, P., Hice, R.H., Wright, J.A., Craig, N.L. and Atkinson, P.W. (2011) The mosquito Aedes aegypti has a large genome size and high transposable element load but contains a low proportion of transposon-specific piRNAs. BMC Genomics, 12, 606.

27. Halbach, R., Miesen, P., Joosten, J., Taşköprü, E., Pennings, B., Chantal B.F. Vogels, Sarah H. Merkling, Constantianus J. Koenraadt, Louis Lambrechts and Ronald P. van Rij (2020) An ancient satellite repeat controls gene expression and embryonic development in Aedes aegypti through a highly conserved piRNA. bioRxiv, http://dx.doi.org/10.1101/2020.01.15.907428.

28. Ichiyanagi, T., Ichiyanagi, K., Ogawa, A., Kuramochi-Miyagawa, S., Nakano, T., Chuma, S., Sasaki, H. and Udono, H. (2014) HSP90α plays an important role in piRNA biogenesis and retrotransposon repression in mouse. Nucleic Acids Res., 42, 11903–11911.

29. Reuter, M., Berninger, P., Chuma, S., Shah, H., Hosokawa, M., Funaya, C., Antony, C., Sachidanandam, R. and Pillai, R.S. (2011) Miwi catalysis is required for piRNA amplification-independent LINE1 transposon silencing. Nature, 480, 264–267.

30. Arensburger, P., Hice, R.H., Zhou, L., Smith, R.C., Tom, A.C., Wright, J.A., Knapp, J., O’Brochta, D.A., Craig, N.L. and Atkinson, P.W. (2011) Phylogenetic and Functional Characterization of the *hAT* Transposon Superfamily. Genetics, 188, 45–57.

31. Aravin, A., Gaidatzis, D., Pfeffer, S., Lagos-Quintana, M., Landgraf, P., Iovino, N., Morris, P., Brownstein, M.J., Kuramochi-Miyagawa, S., Nakano, T., et al. (2006) A novel class of small RNAs bind to MILI protein in mouse testes. Nature, 442, 203–207.

32. Girard, A., Sachidanandam, R., Hannon, G.J. and Carmell, M.A. (2006) A germline-specific class of small RNAs binds mammalian Piwi proteins. Nature, 442, 199–202.

33. Lee, E.J., Banerjee, S., Zhou, H., Jammalamadaka, A., Arcila, M., Manjunath, B.S. and Kosik, K.S. (2011) Identification of piRNAs in the central nervous system. RNA, 17, 1090–1099.

34. Shen, E.-Z., Chen, H., Ozturk, A.R., Tu, S., Shirayama, M., Tang, W., Ding, Y.-H., Dai, S.-Y., Weng, Z. and Mello, C.C. (2018) Identification of piRNA Binding Sites Reveals the Argonaute Regulatory Landscape of the C. elegans Germline. Cell, 172, 937–951.e18.

35. Goh, W.S.S., Falciatori, I., Tam, O.H., Burgess, R., Meikar, O., Kotaja, N., Hammell, M. and Hannon, G.J. (2015) piRNA-directed cleavage of meiotic transcripts regulates spermatogenesis. Genes Dev., 29, 1032–1044.

36. Zhang, P., Kang, J.-Y., Gou, L.-T., Wang, J., Xue, Y., Skogerboe, G., Dai, P., Huang, D.-W., Chen, R., Fu, X.-D., et al. (2015) MIWI and piRNA-mediated cleavage of messenger RNAs in mouse testes. Cell Res., 25, 193–207.

37. Campbell, C.L., Black, W.C., Hess, A.M. and Foy, B.D. (2008) Comparative genomics of small RNA regulatory pathway components in vector mosquitoes. BMC Genomics, 9, 425.

38. Gainetdinov, I., Colpan, C., Arif, A., Cecchini, K. and Zamore, P.D. (2018) A Single Mechanism of Biogenesis, Initiated and Directed by PIWI Proteins, Explains piRNA Production in Most Animals. Mol. Cell, 71, 775–790.e5.

39. Pogorelcnik, R., Vaury, C., Pouchin, P., Jensen, S. and Brasset, E. (2018) sRNAPipe: a Galaxy-based pipeline for bioinformatic in-depth exploration of small RNAseq data. Mob. DNA, 9, 25.

40. Li, H. and Durbin, R. (2010) Fast and accurate long-read alignment with Burrows–Wheeler transform. Bioinformatics, 26, 589–595.

41. Langmead, B. and Salzberg, S.L. (2012) Fast gapped-read alignment with Bowtie 2. Nat. Methods, 9, 357–359.

42. Dufourt, J., Pouchin, P., Peyret, P., Brasset, E. and Vaury, C. (2013) NucBase, an easy to use read mapper for small RNAs. Mob. DNA, 4, 1.

43. Giraldo-Calderon, G.I., Emrich, S.J., MacCallum, R.M., Maslen, G., Dialynas, E., Topalis, P., Ho, N., Gesing, S., the VectorBase Consortium, Madey, G., et al. (2015) VectorBase: an updated bioinformatics resource for invertebrate vectors and other organisms related with human diseases. Nucleic Acids Res., 43, D707–D713.

44. Jenkins, A.M., Waterhouse, R.M. and Muskavitch, M.A. (2015) Long non-coding RNA discovery across the genus anopheles reveals conserved secondary structures within and beyond the Gambiae complex. BMC Genomics, 16, 337.

45. Bao, W., Kojima, K.K. and Kohany, O. (2015) Repbase Update, a database of repetitive elements in eukaryotic genomes. Mob. DNA, 6, 11.

46. Milne, I., Stephen, G., Bayer, M., Cock, P.J.A., Pritchard, L., Cardle, L., Shaw, P.D. and Marshall, D. (2013) Using Tablet for visual exploration of second-generation sequencing data. Brief. Bioinform., 14, 193–202.

47. Grentzinger, T., Armenise, C., Brun, C., Mugat, B., Serrano, V., Pelisson, A. and Chambeyron, S. (2012) piRNA-mediated transgenerational inheritance of an acquired trait. Genome Res., 22, 1877–1888.

48. Kohany, O., Gentles, A.J., Hankus, L. and Jurka, J. (2006) Annotation, submission and screening of repetitive elements in Repbase: RepbaseSubmitter and Censor. BMC Bioinformatics, 7, 474.

49. Lê, S., Josse, J. and Husson, F. (2008) FactoMineR: An R Package for Multivariate Analysis. J. Stat. Softw., 25, 1–18.

50. Quinlan, A.R. and Hall, I.M. (2010) BEDTools: a flexible suite of utilities for comparing genomic features. Bioinformatics, 26, 841–842.

51. Frankish, A., Diekhans, M., Ferreira, A.-M., Johnson, R., Jungreis, I., Loveland, J., Mudge, J.M., Sisu, C., Wright, J., Armstrong, J., et al. (2019) GENCODE reference annotation for the human and mouse genomes. Nucleic Acids Res., 47, D766–D773.

52. Kent, W.J., Sugnet, C.W., Furey, T.S., Roskin, K.M., Pringle, T.H., Zahler, A.M. and Haussler, D. (2002) The human genome browser at UCSC. Genome Res., 12, 996–1006.

53. Favorov, A., Mularoni, L., Cope, L.M., Medvedeva, Y., Mironov, A.A., Makeev, V.J. and Wheelan, S.J. (2012) Exploring massive, genome scale datasets with the GenometriCorr package. PLoS Comput. Biol., 8, e1002529.

54. Jain, A. and Tuteja, G. (2019) TissueEnrich: Tissue-specific gene enrichment analysis. Bioinformatics, 35, 1966–1967.

55. Uhlen, M., Fagerberg, L., Hallstrom, B.M., Lindskog, C., Oksvold, P., Mardinoglu, A., Sivertsson, A., Kampf, C., Sjostedt, E., Asplund, A., et al. (2015) Tissue-based map of the human proteome. Science, 347, 1260419–1260419.

56. Shen, Y., Yue, F., McCleary, D.F., Ye, Z., Edsall, L., Kuan, S., Wagner, U., Dixon, J., Lee, L., Lobanenkov, V.V., et al. (2012) A map of the cis-regulatory sequences in the mouse genome. Nature, 488, 116–120.

