## Supplementary material for "Conserved small nucleotidic elements at the origin of concerted piRNA biogenesis from genes and lncRNAs": Table S13 and multiple sequence alignments SI-MSA1

**Supplemental Figures:**

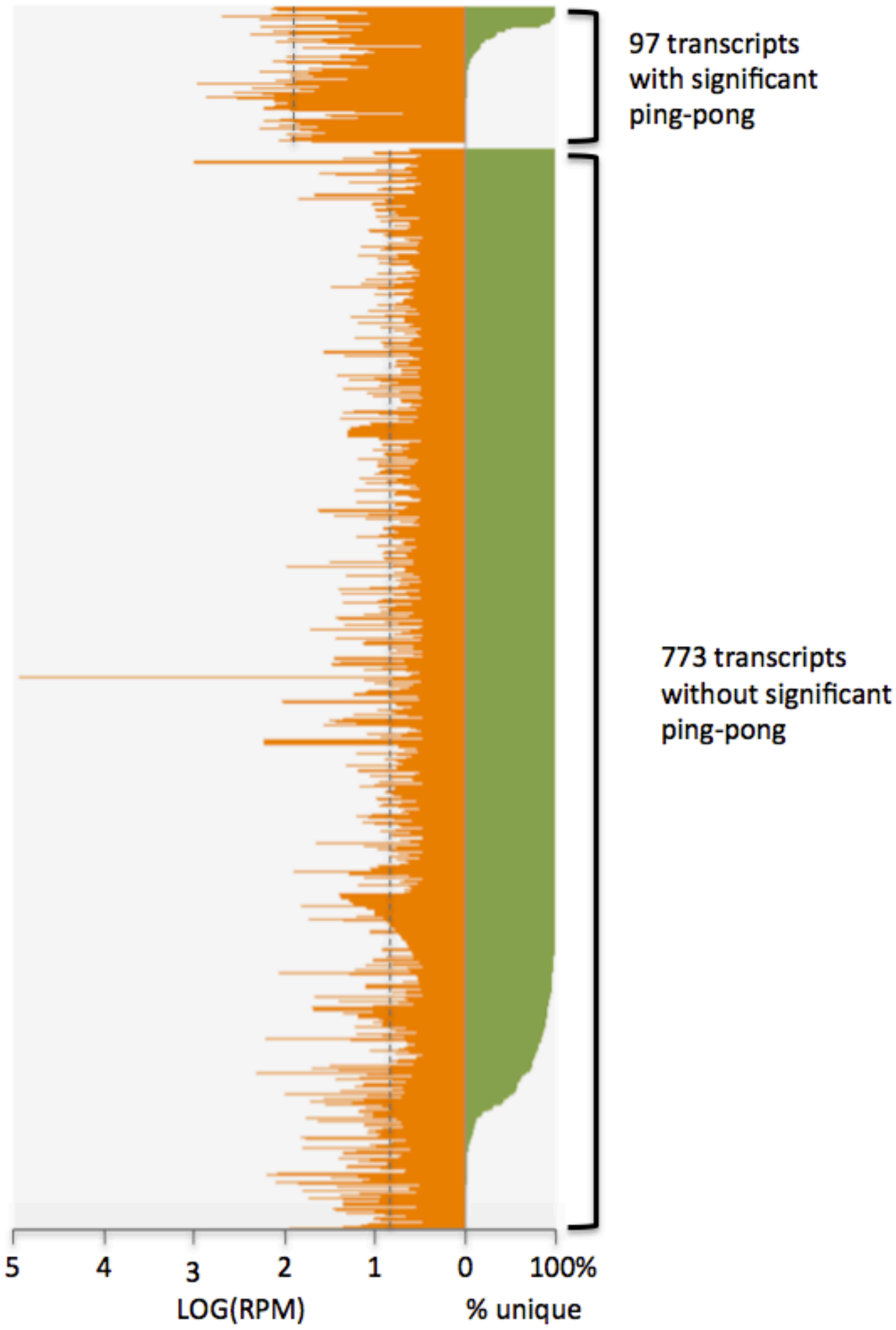


**SFig1 - Amounts of piRNAs and percentages of genome-unique piRNAs.**

Shown are the amounts of piRNAs (LOG of piRNA count in RPM, orange) and percentages of genome-unique piRNAs (green) for each of 870 mRNAs matching >3 RPM piRNAs. For amounts of piRNAs, medians for each category of transcripts are indicated by dotted lines.

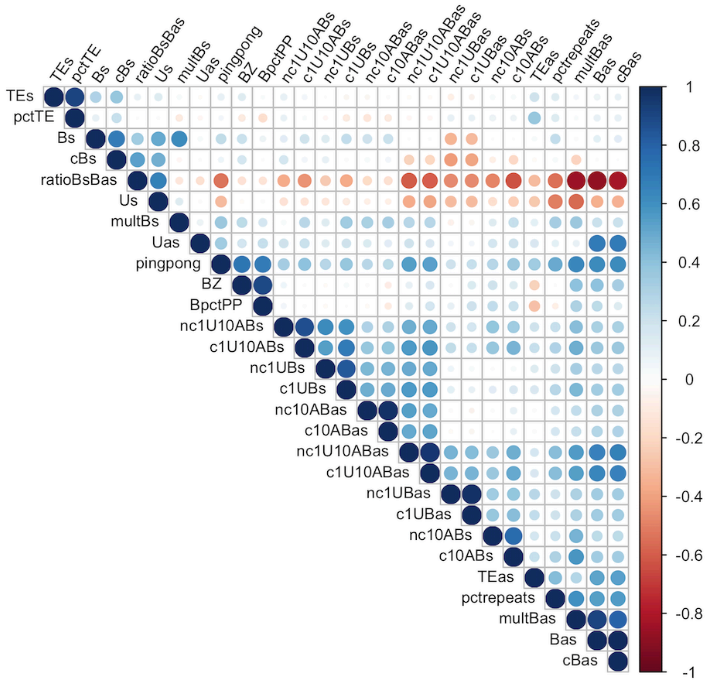


| | |
|---|---|
| pingpong | significant 10-nt-overlap for bona fide piRNAs (yes, no) |
| Bs | count all bonafide sense reads (RPM) |
| Bas | count all bonafide antisense reads (RPM) |
| ratioBsBas | ratio bona fide sense over antisense reads (Bs/Bas) |
| LOGratioBsB | log2(ratioBsBas) |
| BZ | 10-nt-overlap z-score with bonafide reads |
| BpctPP | 10-nt-overlap percentage with bonafide reads |
| Us | count genome-unique sense reads (RPM) |
| Uas | count genome-unique antisense reads (RPM) |
| pctrepeats | percentage genome-repeated reads (1-[(Us+Uas)/(Bs+Bas)]) |
| TEs | count insect-TE-matching sense reads (RPM) |
| TEas | count insect-TE-matching antisense reads (RPM) |
| pctTE | percentage insect-TE-matching piRNAs [(TEs+TEas)/(Bs+Bas)] |
| nc1UBs | 1U % for bonafide sense reads |
| nc1UBas | 1U % for bonafide antisense reads |
| nc10ABs | 10A % for bonafide sense reads |
| nc10ABas | 10A % for bonafide antisense reads |
| nc1U10ABs | 1U-10A % for bonafide sense reads |
| nc1U10ABas | 1U-10A % for bonafide antisense reads |
| cBs | count bonafide sense reads collapsed (RPM) |
| cBas | count bonafide antisense reads collapsed (RPM) |
| multBs | multiplicity for bonafide sense reads (Bs/cBs) |
| multBas | multiplicity for bonafide antisense reads (Bas/cBas) |
| c1UBs | 1U % for bonafide sense reads collapsed |
| c1UBas | 1U % for bonafide antisense reads collapsed |
| c10ABs | 10A % for bonafide sense reads collapsed |
| c10ABas | 10A % for bonafide antisense reads collapsed |
| c1U10ABs | 1U-10A % for bonafide sense reads collapsed |
| c1U10ABas | 1U-10A % for bonafide antisense reads collapsed |
| UZ | 10-nt-overlap z-score with genome-unique reads |
| UpctPP | 10-nt-overlap percentage with genome-unique reads |

**SFig2 - Correlogram obtained by Spearman method on 869 mRNAs matching >3RPM piRNAs (without AGAP003387).**

There is a weak correlation between ping-pong and the percentage of repeated piRNAs for bonafide and non-TE reads, but not for genome-unique reads. But there is no correlation between ping-pong and the percentage of TE-related piRNAs (Note that there are only very few TE-related reads). There is correlation between the ping-pong parameters themselves: z-scores, percentage of 10nt-overlap and the resulting yes/no record, and a correlation between bonafide antisense reads (Bas) and ping-pong parameters (z-score, etc.). There also is correlation between bonafide antisense reads (Bas) and TE-matching antisense reads and the percentage of repeated piRNAs.

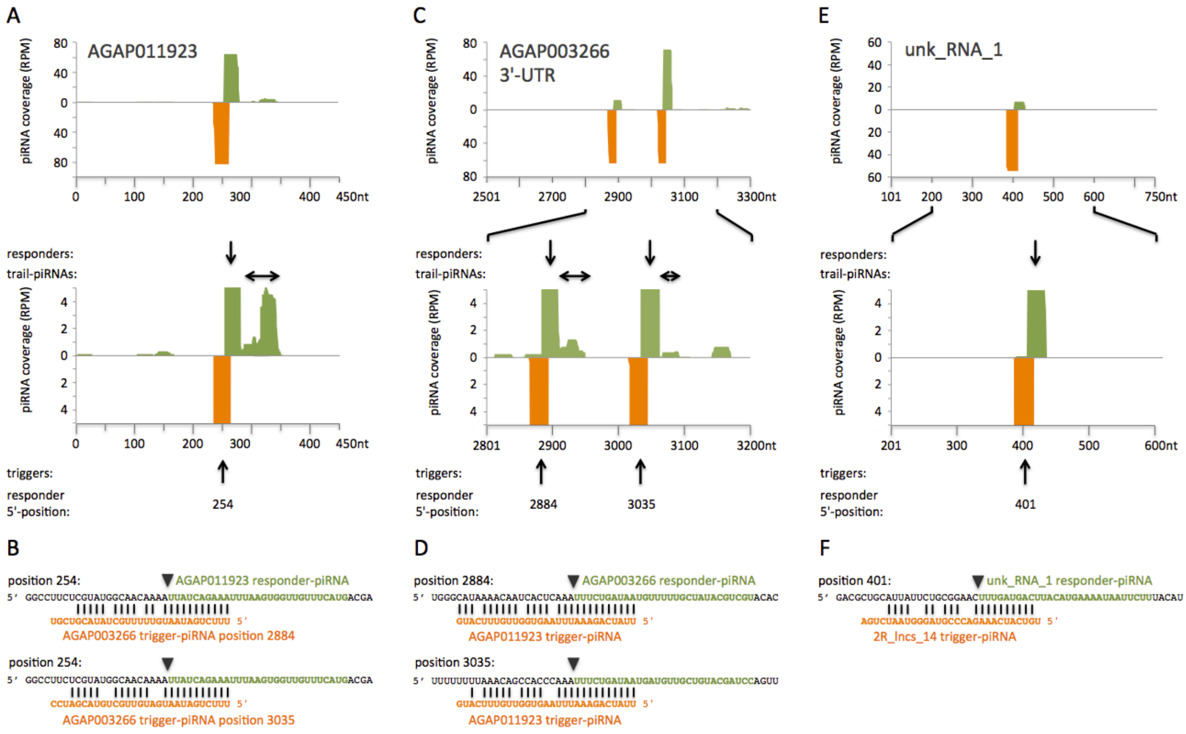


**SFig3 - piRNA-based network involving mRNAs and lncRNAs in *An. gambiae* (Complement to Figure 4).**

(A,C,E) Mapping of piRNAs to network transcripts. The entire AGAP011923-RA mRNA is represented (A), AGAP003266 3'-UTR (2R:34610936-34611735[+]) (C) and unk_RNA_1 (2R:21592981-21593730[+]) (E). Shown are all piRNAs mapping the transcripts without mismatch (green) and the putative trigger-piRNAs (orange, mismatched pairing). All these piRNAs are genome-unique. Plus-oriented piRNAs are shown in the upper part, minus-oriented piRNAs in the lower part of each figure. The responder- and trail-piRNAs (green) are indicated, respectively, by vertical and horizontal arrows above the zoom-in figure. Arrows below point to the trigger-piRNAs (orange), together with the 5'-position of the corresponding responder-piRNAs. (B,D,F) Sequences of the responder-piRNAs (green) and trigger-piRNAs (orange) for each trigger-responder pair. Transcript sequences surrounding the responder sequences are in black. The 5'-end position is indicated for each responder-piRNA. The inverted triangles show the slicer cleavage position facing nucleotides 10-11 of the trigger-piRNA. (A,B) Mapping of piRNAs to the AGAP011923-RA transcript. Trigger-piRNAs originate from the AGAP003266 3'-UTR. (C,D) Mapping of piRNAs to the AGAP003266 3'-UTR. Trigger-piRNAs originate from AGAP011923. (E,F) Mapping of piRNAs to unk_RNA_1. Trigger-piRNAs originate from 2R_lncs_14.

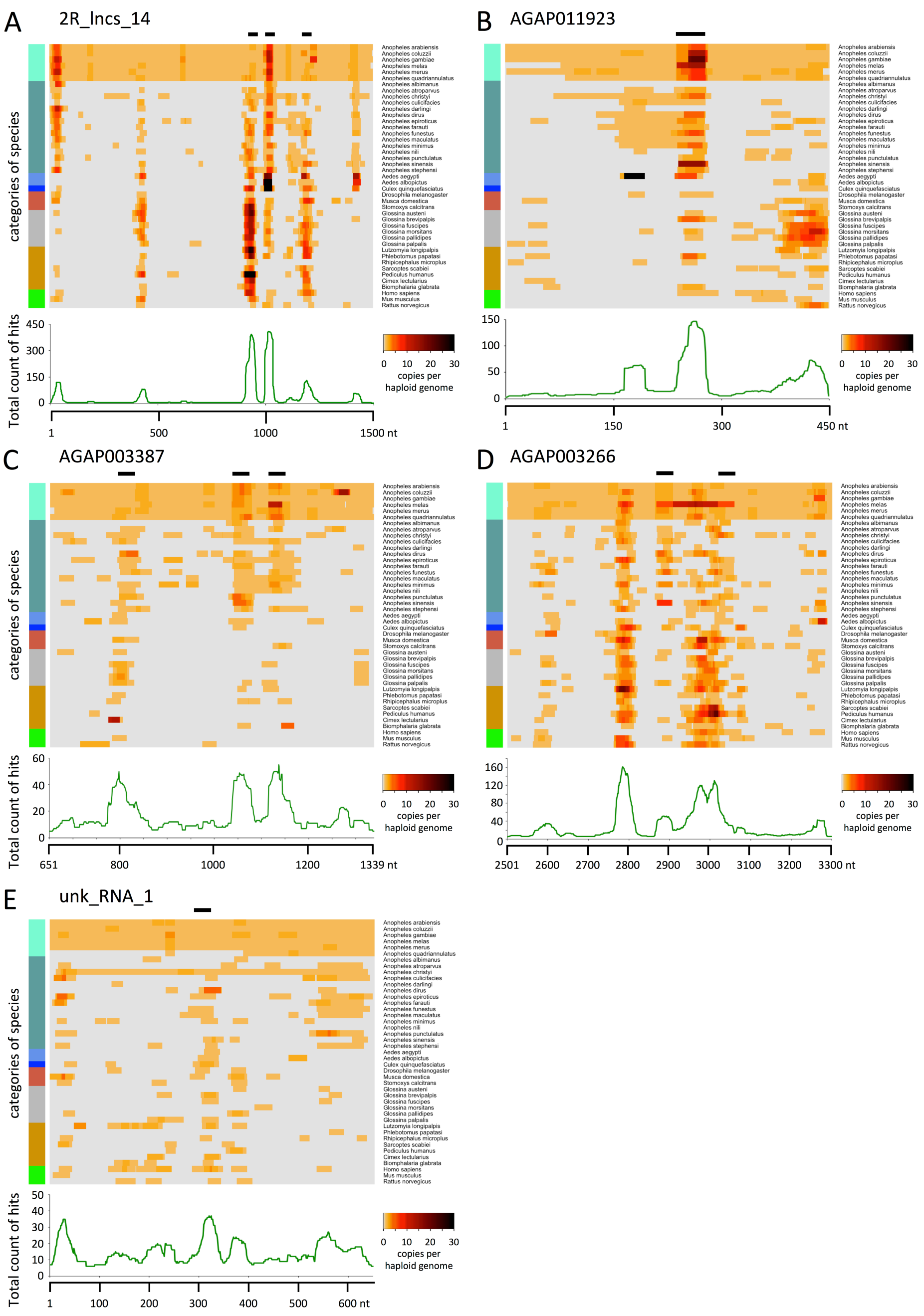


**SFig4: BLASTN results reveal short conserved repeats, the snetDNAs (e-value ≤1). (Complement to Figure 5)**

Heat maps illustrating Blastn results, with a maximal e-value of 1.0, for network transcripts 2R_lncs_14 (A), AGAP011923-RA (B), AGAP003387-RA (C), AGAP003266 3'-UTR (D), and unk_RNA_1 (E) aligned to genomes of 43 different species. The counts of hits are plotted along each transcript for each individual species. Zero hit is in light grey. A color code for different categories of species is presented on the left. Mosquitoes are in different blue colors: *An. gambiae* complex, light blue, other *Anopheles*, cadet blue, *Aedes*, cornflower blue, and *Culex, blue*; coral: flies; gray: tsetse; green: mammals; dark golden: other species including snail, body louse, bed bugs, tick, mite, sand flies. Horizontal bars above the heat map indicate the trigger-responder sites (spanning over the trigger-annealing site and the corresponding responder-piRNAs). Below each heat map, a graphical representation of the totalized counts of hits for all analyzed species mapped for each position of the transcript. The detailed BLASTN results with the aligned sequences can be found in Supplemental Table S10.

**Trigger-piRNAs:**

| | | |
|---|---|---|
| *An. gambiae* AGAP011923: | UUAUCAGAAAUUUAAGUGGUUGUUUCAUG | 63.9 RPM |
| *Ae. aegypti* AAEL017228: | UUAAUAGAAAUUACAGUGGUUGGUAUCUU | 3505.6 RPM |
| *Ae. aegypti* AAEL009512-3'-region: | UUAAUAGAAAUUACAGUGGUUGGUCUUUU | 556.6 RPM |
| Mouse fetal testes piRNA: | UUAUAUGUAAGUACACUG–UAGCUGUCU | 25.6 RPM |
| Mouse adult testes piRNA: | UUAUAUGUAAGUACACUG–UAGCGGUCUU | 5.0 RPM |

**Responder-piRNAs:**

| | | |
|---|---|---|
| *An. gambiae* 2R_lncs_14 position 1009: | UUUCUCAAUAU–UUUCUAUACCCUAAAUUU | 11.6 RPM |
| *Ae. aegypti* 2R_lncs_14 ortholog: | UUUCUCAAUAU–UUUCUAUUUCCUAUUCGU | 270.3 RPM |
| *Ae. aegypti* 2R_lncs_14 ortholog: | UUUCUCUAUAUAUUUCUAUUUAC–AAUUCG | 32.0 RPM |
| *An. gambiae* 2R_lncs_14 position 1407: | UGUGAUAUUAUCACUA–CGGAAAUAUUUUC | 8.0 RPM |
| *Ae. aegypti* 2R_lncs_14 ortholog: | CGUGAUAUUAUCACUA–CGGAA–UAUUAUUU | 782.8 RPM |
| Mouse adult testes piRNA: | UCUGUCAACA–GACUAUCAGAAAU–UUCAAAGC | 4.1 RPM |

**SFig5 - SnetDNA-network piRNAs in *Ae. aegypti* and similar piRNAs matching snetDNAs in mouse fetal and adult testes.**

Shown are network piRNAs found in *Ae. aegypti* and piRNAs matching snetDNAs, drawn from data of Figure 5, in mouse fetal and adult testes. Dissimilarities with respect to the corresponding *An. gambiae* piRNAs are highlightened in red. Nucleotides that are identical between mouse and *Ae. aegypti* piRNAs are in green. 1U and 10A are in bold and pointed by black dots. On the right, the respective amounts of piRNAs (RPM).

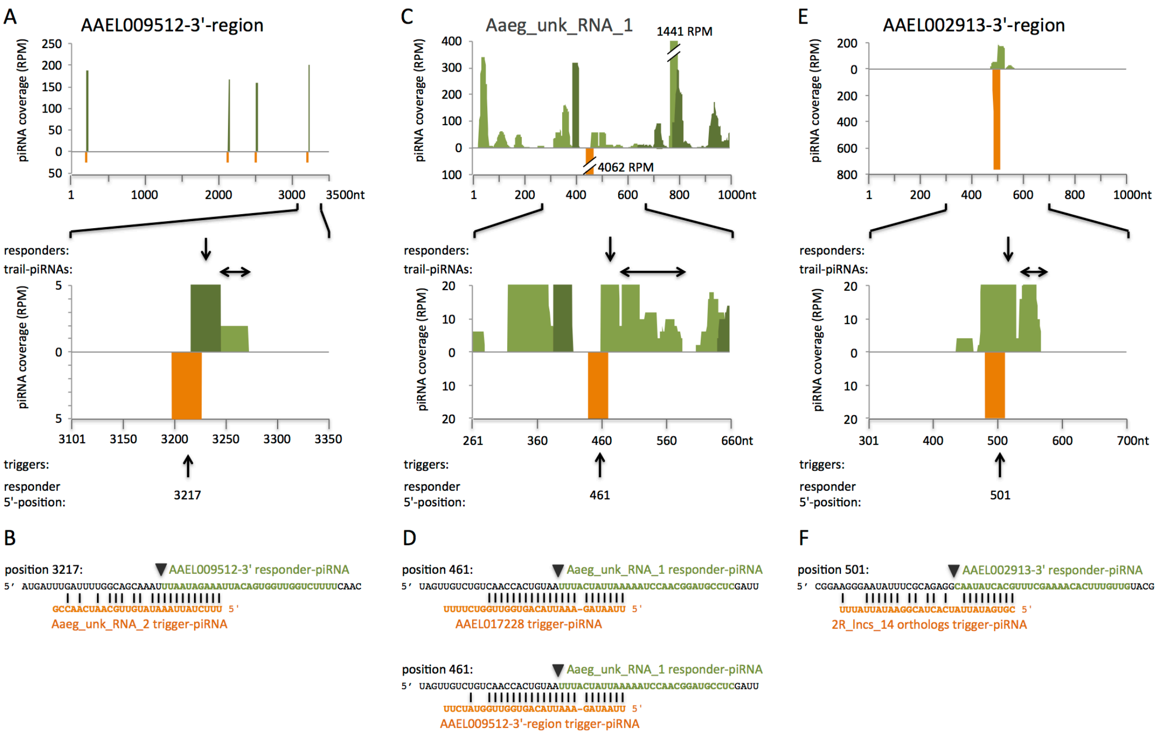


**SFig6 - SnetDNA-network involving mRNAs and lncRNAs in *Ae. aegypti* (Complement to Figure 6)**

(A,C,E) Mapping of piRNAs to network transcripts. Shown are all piRNAs mapping the transcripts without mismatch (in light green when genome-unique, in dark green when repeated in the genome) and the putative trigger-piRNAs (orange, mismatched pairing). Plus-oriented piRNAs are shown in the upper part, minus-oriented piRNAs in the lower part of each figure. The responder- and trail-piRNAs (green) are indicated, respectively, by vertical and horizontal arrows above the zoom-in figure. Arrows below point to the trigger-piRNAs (orange), together with the 5'-position of the corresponding responder-piRNAs. (B,D,F) Sequences of the responder-piRNAs (green) and trigger-piRNAs (orange) for each trigger-responder pair. Transcript sequences surrounding the responder sequences are in black. The 5'-end position is indicated for each responder-piRNA. The inverted triangles show the slicer cleavage position facing nucleotides 10-11 of the trigger-piRNA. (A,B) Mapping of piRNAs to the AAEL009512 3'-region (AaegL3_supercont1401:478162-481661[-]). Trigger-piRNAs originate from Aaeg_unk_RNA_2. (C,D) Mapping of piRNAs to Aaeg_unk_RNA_1 (AaegL3_supercont1478_556717-557716[+]). Trigger-piRNAs originate from AAEL017228 and the AAEL009512 3'-region. (E,F) Mapping of piRNAs to the AAEL002913 3'-region (AaegL3_supercont1-71_1676720-1677719[-]). Trigger-piRNAs originate from 2R_lncs_14 orthologs.

### Table S13

| piRNA nb. | *Ae. aegypti* locus | corresponding *An. gambiae* locus | nature of piRNA | piRNA-sequence *Ae. aegypti* (corresponding piRNA-sequence in *An. gambiae*) | mapnum vs. genome | mapnum vs. 2R_lncs_14 orthologs | RPM |
|---|---|---|---|---|---|---|---|
| 1 | AAEL017228-RA | AGAP011923-RA | trigger/ responder | UUAAUAGAAAUUACAGUGGUUGGUAUCUU (UUAUCAGAAAUUUAAGUGGUUGUUUCAUG) | 1 | 0 | 3505.6 |
| 2 | AAEL009512-3'-region | AGAP011923-RA | trigger/ responder | UUAAUAGAAAUUACAGUGGUUGGUCUUUU (UUAUCAGAAAUUUAAGUGGUUGUUUCAUG) | 1 | 0 | 556.6 |
| 3 | Aaeg_unk_RNA_1 | none | responder | UUUACUAUUAAAAAUCCAACGGAUGCCUC | 1 | 0 | 58.1 |
| 4 | 2R_lncs_14 orthologs | 2R_lncs_14 position 1009 | responder | UUUCUCAAUAUUUUCUAUUUCCUAUUCGU (UUUCUCAAUAUUUUCUAUACCCUAAAUUU) | 27 | 25 | 270.3 |
| 5 | AAeg_2R_lncs_14-J (2R_lncs_14 ortholog) | 2R_lncs_14 position 1009 | responder | UUUCUCUAUAUAUUUCUAUUUAC-AAUUCG (UUUCUCAAUAU-UUUCUAUACCCUAAAUUU) | 5 | 5 | 32.0 |
| 6 | 2R_lncs_14 orthologs | 2R_lncs_14 position 1407 | responder | CGUGAUAUUAUCACUACGGAA-UAUUAUUU (UGUGAUAUUAUCACUACGGAAAUAUUUUC) | 42 | 42 | 782.8 |
| 7 | AAEL002913-3'-region | none | trigger/ responder | CAAUAUCACGUUUCGAAAACACUUUGUUG | 1 | 0 | 170.2 |

#### SI-MSA1

#### Supplemental Information: Multiple Sequence Alignment Ping-pong networks

```
Ping-pong network 1:
CLUSTAL O(1.2.4) multiple sequence alignment (https://www.ebi.ac.uk/Tools/msa/clustalo/)

AGAP012458-RA      ------------------------------------------------------------  0
AGAP001088-RA      ------------------------------------------------------------  0
AGAP012431-RA      ------------------------------------------------------------  0
AGAP012766-RA      ------------------------------------------------------------  0
AGAP003677-RA      AGGTTGTGTGTGTAAAAAAACGTAAACAGTGTTGTTGGCTGATCAA-TTTTGGTAAGTTT  59
AGAP012489-RA      --------------------------------------------CAAGTGCGGTAAGTTT  16
AGAP007403-RA      AAGTTGTGTGTGTAAAAAAACCTAAACAGTGTTGTTGGCTGATCAAATTTTGGTAAGTTT  60
AGAP012182-RA      ------------------------------------------------------------  0


AGAP012458-RA      ------------------------------------------------------------  0
AGAP001088-RA      -------------GTCTTAATTTTACGTAAGTAAGTGCCATTTTAGCAAAAGTTAGTGGT  47
AGAP012431-RA      -------------GTCTTAATTTTACGTAAGTAAGTGCCATTTTAGCAAAAGTTAGTGGT  47
AGAP012766-RA      -------------------------------------------------------GTGGT  5
AGAP003677-RA      TTTAATAATTAAGTTCTTAATTTTACGTAAGTAAGTGCCATTTTAGCTAAATTAAGTGGT  119
AGAP012489-RA      TTTAATAATTATGTTCTTAATTTTACGTAAGTAAGTGCCATTTTAGCAAAAGAAAGTGGT  76
AGAP007403-RA      TTTAATAATTTAGTTCTTAATTTTACGTAAGTAAGTGCCATTTTAGCAAAAGTAAGTGGT  120
AGAP012182-RA      -------------------------------------------------------GTGGT  5


AGAP012458-RA      ------------------------------------------------------------  0
AGAP001088-RA      AGAAGTCCTGCCCAATGCGTGCTGGCTCCGTCGAAAGCGTTACGTGCATGTACCGGCGTA  107
AGAP012431-RA      AGAAGTCCTGCCCAATGCGTGCTGGCTCCGTCGGAAGCGTTACGTGCATGTACCGGCGTA  107
AGAP012766-RA      AGAAGTCCTGCCTAATGTGTGCTGGCTCCGTCGGAAGCGTTACGTGCATGTACCGGCGTA  65
AGAP003677-RA      AGAAATCCTGCCCAATGCGTGCTGGCTCCGTCGGAAGCGGTACGTGTATGTACCGGCGTA  179
AGAP012489-RA      AGAAGTCCTGCCCAATGTGTGCTGGCTCCGTCGGAAGCGTTACGTGCATGTACCGGCGTA  136
AGAP007403-RA      AGAAGTCCTGCCCAATGCGTGCTGGCTCCGTCGGAAGCGGTACGTGCATGTACCGGCGTA  180
AGAP012182-RA      AGAAGTCCTGCCCAATGCGTGCTGGCTCCGTCGGAAGCGGTACGTGCATGTACTGGCGTA  65


AGAP012458-RA      ------------------------------------------------------------  0
AGAP001088-RA      GGAAAGTGATGTGGTGAACAACACAGCACAAGACAGCGGATATAATGCA-----------  156
AGAP012431-RA      GGAAAGTGATGTGGTGAACAACACAGCACAAGACAGCGGATATAATGCA-----------  156
AGAP012766-RA      GGAAAGTGGTGTGGTGAACAACACAGCACAAGACAACG----------------------  103
AGAP003677-RA      GAAAAGTGGCGTGATGAACAACACAGCACAAGACAGCA----------------------  217
AGAP012489-RA      GGAAAGTGGTGTGGTGAACAACACAGCACAAGACAGCG----------------------  174
AGAP007403-RA      GGAAAGTGGTGTGGTGAACAACACAGCACAAGACAACGGGTAAGTGTTATCCAGCGATTA  240
AGAP012182-RA      GGAAAGTGGTGTGATGAACAACACAGCAT-------------------------------  94


AGAP012458-RA      ------------------------------------------------------------  0
AGAP001088-RA      ------------------------------------------------------------  156
AGAP012431-RA      ------------------------------------------------------------  156
AGAP012766-RA      ---------------------------------------------------GATATAATG  112
AGAP003677-RA      ---------------------------------------------------GATATAATG  226
AGAP012489-RA      ---------------------------------------------------GATATAATT  183
AGAP007403-RA      TCATGATTAAGAAACATTATTTTAATAGAATATGGCCACTTTTAACTTGCAGATATAATG  300
AGAP012182-RA      ------------------------------------------AAGACAACGGATATAATG  112


AGAP012458-RA      ------------------------------------------------------------  0
AGAP001088-RA      --GAAATGATA------TGCGGCAGGTGTTTGCGACTGTA-----CGTTGCGTTGTATAT  203
AGAP012431-RA      --GAAATGATA------TGCGGCAGGTGTTTGCGACTGTA-----CGTTGCGTTGTATAT  203
AGAP012766-RA      CAGAAATGATATGGGCCTGCCTCAGGTGTTTGCGACTGTGCGCTGTGTATGATTGTGTAT  172
AGAP003677-RA      CAGAAACGAAATGGGCCTGCCGCAGGTGATTGCCACTGTGCGCTGTGTATGATTGTGTAT  286
AGAP012489-RA      CAGAAACGATATGGGCCTGCGGCAGGTGTTTGCGACTGTGCGCTGTGTATGATTGTGTAT  243
AGAP007403-RA      CAGAAACGATATGGGCCTGCGGCAGGTGTTTGCGACTGTGCGCTGTGTATGATTGTGTAT  360
AGAP012182-RA      CAGAAACGATATGGGCCTGCGGCAGGTGTTTGCGACTGTGCGCTGTGTATGATTGTGTAT  172


AGAP012458-RA      ------------------------------------------------------------  0
AGAP001088-RA      GATTCGCCAATTTTCCTCTATCCT-GGAACGTGGGCTTACGAGGAACCATTTAAGAACAT  262
AGAP012431-RA      GATTCGCCAATTTTCCTCTATCCT-GGAACGTGGGCTTACGAGGAACCATTTAAGAACAT  262
AGAP012766-RA      GATTCGCCAATTTTCCTCTATCCGTGGAACGTGGGGTTACGAGGAACCGTTCAAGAGGAT  232
AGAP003677-RA      GATTCGCCAATTTTCCTCTATCCT-GGAACGTGGGCTTACGAGGAACCATTTAAGAACAT  345
AGAP012489-RA      GATACGCCAATTTTCCTCTATCCT-GGAACGTGGGCTTACGAGGAACCGTTCAAGAAGAT  302
AGAP007403-RA      GATACGCCAATTTTCCTCTATCCT-GGAACGTGGACTTACGAGGAACCATTTAAGAACAT  419
AGAP012182-RA      GATACGCCAATTTTCCTCTATTCT-GGAACGTGGTGTTACGATGAACCATTAAAGAACAT  231
```

```
AGAP012458-RA      --------------------------------------------CACGCTAGAGGCCCAG  16
AGAP001088-RA      CATCCGGAAGCTGTACGGCGACGAGTCGGGCTCTACCAGCACAACACGCTGA-GGCCCAG  321
AGAP012431-RA      CATCCGGAAGCTGTACGGCGACGAGTCGGGCTCTACCAGCACAACACGCTGA-GGCCCAG  321
AGAP012766-RA      CCTCCGGAAGCTGTACGGCGACGAGTCGGGCTCTACCAGCACAACACGCTAGAGGCCCAG  292
AGAP003677-RA      CCTCCGGAAGCTGTACAGCGACGAGTCGGGCTCTACCAGCACAACACGCTAGAGGCCCAG  405
AGAP012489-RA      CCTCCGGAAGCTGTACGGCGACGTGTCGGGCTCTACCAGCACAACACACTAGAGGCCCAG  362
AGAP007403-RA      CCTCCGGAAGCTGTACGGCGACGAGTCGGGCTCTACCAGCACAACACGCTAGAGACCCAG  479
AGAP012182-RA      CCTCCGGGAGCTGTACGGCGACGAATCGGGCTCTACCAGCACAACACGCTAGAGACCCAG  291
                                                               *** **   * *****

AGAP012458-RA      CCGGCACCGGGTGCCCCGATTGCGCCGCCAAACGGTGTCGAGCGCGCTCGTTCAAAACGT  76
AGAP001088-RA      CCGGCACCGGGTGCCCCGATTGCGCCGCCGGACGGTGTCGAGGAAAAAACGTTGAAACGT  381
AGAP012431-RA      CCGGCACCGGGTGCCCCGATTGCGCCGCCGGACGGTGTCGAGGAAAAAACGTTGAAACGT  381
AGAP012766-RA      CCGGCACCGGGTGCCCCGATTGCGCCGTCAGACGGTGTAGAG--CGCTCGTTCACAACGT  350
AGAP003677-RA      CCGGTACCGGGTGCCCCGATTGCGCCGTGAGACGGTGTCGAGCGCGCTCGTTAACAACGT  465
AGAP012489-RA      CCGGTACCGGGTGCCCCGATTGCGCCGCCAGACGGTGTCGAGCGCGCTCGTTAACAACGT  422
AGAP007403-RA      CCGGCACCGGGTGCCCCGATTGCGCCGCCAGACGTTGTCGAGCGCGCTCGTTCACAACGT  539
AGAP012182-RA      CCGGCACCGGGTGCCCCGATTGCGCCGTCAGACGGTGTCGAGCGCGCTCGTTCACAACGT  351
                   **** **********************    *** *** ***         *   *****

AGAP012458-RA      TCGGGGCAGGAATTCAACCCTTCGGATCATT---ATCCTCTGCCTCACAATAGCTGCAGC  133
AGAP001088-RA      TCGGGGCAGGAATTCAACCCTTCGGATCCTCATTATCCTCTGCCCCACAATAGCTGCAGC  441
AGAP012431-RA      TCGGGGCAGGAATTCAACCCTTCGGATCCTCATTATCCTCTGCCCCACAATAGCTGCAGC  441
AGAP012766-RA      TCGGGACAGAAATACAACCCTTCGGATCATT---ATCCTCTGCCTCACAATAGCTGCAGC  407
AGAP003677-RA      TCGGGGCAGGAATTCAACCCTTCGGATCATTAT---CCTCTGCCTCACAATAGCTGCAGC  522
AGAP012489-RA      TCGGGGCAGGAATTCAACCCTTCGGATCTGCATTATCCTCTGCCTCACAATAGCTGCAGC  482
AGAP007403-RA      TCGGGGCAGGAATTGAACCCTTCGGATCATT---ATCCTCTGGCTCACAATAGCTGCAGC  596
AGAP012182-RA      TCGGGGCAGGAATTCAACCCTTCGGATCTGCATTATCCGCTGTCCCACAATAGCTGCAGC  411
                   ***** *** ***  *************        ** *** * ***************

AGAP012458-RA      AGCAACAGGACCGAACCGAC---CGCACTACTGATTGCGGTCAAATTTCCTGGCAGGAGA  190
AGAP001088-RA      AGCAACAGGACCGAACCGAC---CGCACTACTGATTGCGGTCAAATTTCCTGGCAGGAGA  498
AGAP012431-RA      AGCAACAGGACCGAACCGAC---CGCACTACTGATTGCGGTCAAATTTCCTGGCAGGAGA  498
AGAP012766-RA      AGCAACAGGACTGAACCGAC---CGCACTACTGATTGCGGTCAAATTTCCTGGCAGGAGA  464
AGAP003677-RA      AGCATCAGGACCGAACCGACCACCGCACTACTGATTGCGGTCAAATTTCGTG--------  574
AGAP012489-RA      AGCAACAGGACCGAACCGAC---CGCACTA------------------------------  509
AGAP007403-RA      AGCAACAGGACCGAACCGAC---CGCACTACGGATTGCGGTCAAATTTTCTGGCAGGAGA  653
AGAP012182-RA      AGCAACAGGACCGAACCGAC---CGCAGTACTGATTGCGGTCAAATTTCCTG--------  460
                   **** ****** ********   **** **

AGAP012458-RA      AAAAACCCCACAAGCGTATCGATTTCATCGACGGCCGCTCGCTGACGGCACCAACATGCC  250
AGAP001088-RA      AAAAACCCCACGAGTGTATCGATTTCATCGACGGTCGCTCGCTGGCAGCACCGACAAACA  558
AGAP012431-RA      AAAAACCCCACGAGTGTATCGATTTCATCGACGGTCGCTCGCTGGCAGCACCGACAAACA  558
AGAP012766-RA      AAAAATCCCACAAGTGTATCGATTTCATCGACGGCCGCTCGCTGGCGGCACCAACATACC  524
AGAP003677-RA      ------------------------------------------------------------  574
AGAP012489-RA      ------------------------------------------------------------  509
AGAP007403-RA      -AAAACCCCACGAGTGTATCGATTTCATGGACGGCCGCTCGCTGACGGCACCAACATGCC  712
AGAP012182-RA      ------------------------------------------------------------  460


AGAP012458-RA      ACTGAATAACCCCGGAAAGGCGACAAGCACTCATTTCCAGCGGTAAATTGTTAACATCAA  310
AGAP001088-RA      CTCATCACAGTGGCGACAGTGGCAAATTGTTAACGTGAACCCACTTCCGCC-CATCTACA  617
AGAP012431-RA      CTCATCACAGTGGCGACAGTGGCAAATTGTTAACGTGAACCCACTTCCGCC-CATCTACA  617
AGAP012766-RA      ACTGGATAACCTTGGAAAGGCGACAAACATCACGACATCACGTGAATAGTGTTATTGAGT  584
AGAP003677-RA      ------------------------------------------------------------  574
AGAP012489-RA      ------------------------------------------------------------  509
AGAP007403-RA      ACTGAATAACCCCGGAAAGACGACAAGCACTCATCTCCAGCGGCAAATTGTTAACATCAA  772
AGAP012182-RA      ------------------------------------------------------------  460


AGAP012458-RA      TCCTCTTCCGCACATCTACACGACATCATGATAATAGTGTTATTGTGTAAATTGTAAAAA  370
AGAP001088-RA      CGACATCGCGT-------------------------------------------------  628
AGAP012431-RA      CGACATCGCGT-------------------------------------------------  628
AGAP012766-RA      TTATTGAACAA-------------------------------------------------  595
AGAP003677-RA      ------------------------------------------------------------  574
AGAP012489-RA      ------------------------------------------------------------  509
AGAP007403-RA      TCCTCTTCCGC-------------------------------------------------  783
AGAP012182-RA      ------------------------------------------------------------  460


AGAP012458-RA      AAAATACGTTAATGAGTTAAGATTAATCTTAAGTGCGCTTTAAGGCATACATCGATATGC  430
AGAP001088-RA      TAATAGTGTTATTGTGTTAATTGGAAAAATAACTTTATG-------TTAACGAGTTAAGT  681
AGAP012431-RA      TAATAGTGTTATTGTGTTAATTGGAAAAATAACTTTATG-------TTAACGAGTTAAGT  681
AGAP012766-RA      ACTTGATGTTAACGAGTTAAGTTTAATTTTATATGCGCTTTAGGGCATAAATCGATATGC  655
AGAP003677-RA      ------------------------------------------------------------  574
AGAP012489-RA      ------------------------------------------------------------  509
AGAP007403-RA      ACATCTAC----------------------------------------------------  791
AGAP012182-RA      ------------------------------------------------------------  460
```

```
AGAP012458-RA      CTAATTTTAGGTTAAGTGTTTTGAAATGCGCGTGTGTGCGAAGGGTCGAAATGAGCGCTA  490
AGAP001088-RA      TTAATCTTAAG-------------------------------------------------  692
AGAP012431-RA      TTAATCTTAAG-------------------------------------------------  692
AGAP012766-RA      CTAATTTTAAGTTAAGTGTTTTGACATGTGCGTGTGTGTGAAGGGGCAAAATAAGCGCTA  715
AGAP003677-RA      ------------------------------------------------------------  574
AGAP012489-RA      ------------------------------------------------------------  509
AGAP007403-RA      ------------------------------------------------------------  791
AGAP012182-RA      ------------------------------------------------------------  460


AGAP012458-RA      AGCAAACTAAGCGGCCGCGTAGGGCCCCGTGGTCTTGAGGGGCCCCAAAAATGGAACCCA  550
AGAP001088-RA      ------------------------------------------------------------  692
AGAP012431-RA      ------------------------------------------------------------  692
AGAP012766-RA      AGCAAACTAAGCGGCCATGTAGGGCCCCGTGGTCTTGAGGGGCCCCAAAAATGGAACCCA  775
AGAP003677-RA      ------------------------------------------------------------  574
AGAP012489-RA      ------------------------------------------------------------  509
AGAP007403-RA      ------------------------------------------------------------  791
AGAP012182-RA      ------------------------------------------------------------  460

AGAP012458-RA      CCCCTCGCTCGCTAGTAAAATTTACATTTGTGCTTGTAAACC------------------  592
AGAP001088-RA      ------------------------------------------------------------  692
AGAP012431-RA      ------------------------------------------------------------  692
AGAP012766-RA      CCCTCCGCTCGCAAGTAAAATTGCTACTCAAACCTTCTCTATACATTTACCTGCTAAACG  835
AGAP003677-RA      ------------------------------------------------------------  574
AGAP012489-RA      ------------------------------------------------------------  509
AGAP007403-RA      ------------------------------------------------------------  791
AGAP012182-RA      ------------------------------------------------------------  460

AGAP012458-RA      -------------TACATTTA--------TGAGACTGCCTCCACTCGTATGTGCGGGTTT  631
AGAP001088-RA      ------------------------------------------------------------  692
AGAP012431-RA      ------------------------------------------------------------  692
AGAP012766-RA      CTTCAACCAAACCTACATTTATGTGACTGATTATGGGCCTCCACTCGTATGTGCGGATTT  895
AGAP003677-RA      ------------------------------------------------------------  574
AGAP012489-RA      ------------------------------------------------------------  509
AGAP007403-RA      ------------------------------------------------------------  791
AGAP012182-RA      ------------------------------------------------------------  460

AGAP012458-RA      TATACAACATTTAAGATAAATAAATCTAGTTTATATTCTCAT      673
AGAP001088-RA      ------------------------------------------      692
AGAP012431-RA      ------------------------------------------      692
AGAP012766-RA      TACACAACATTTTAGATAAATAAATCTAGTTTA---------      928
AGAP003677-RA      ------------------------------------------      574
AGAP012489-RA      ------------------------------------------      509
AGAP007403-RA      ------------------------------------------      791
AGAP012182-RA      ------------------------------------------      460
```

Ping-pong network 1:
Sequence Logo 3.5.0 (toolshed.g2.bx.psu.edu/repos/devteam/weblogo3/rgweblogo3/3.5.0)

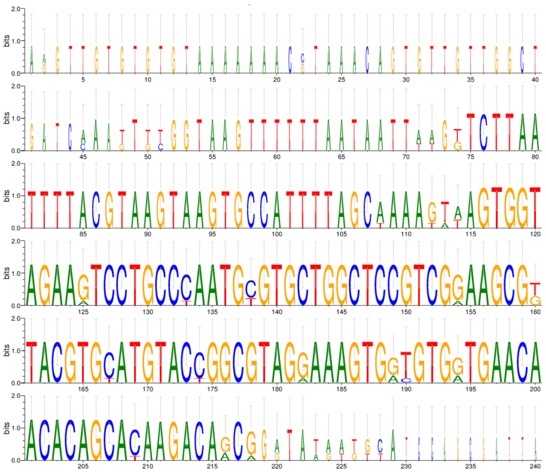

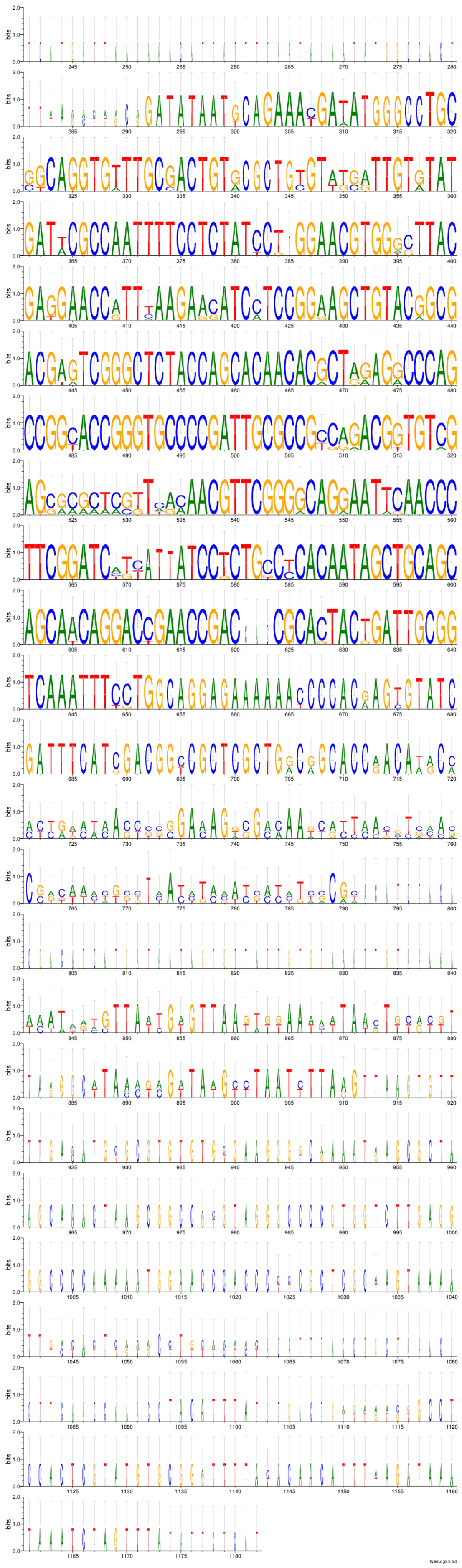
WebLogo 3.5.0

Ping-pong network 2:
CLUSTAL 2.1 multiple sequence alignment

```
AGAP012483-RA      GGACCGGTCGTCGATGCACTCGAGGTG-CACCGCATTGACGAGCGGCAGTAGCAGGATGC 59
AGAP012762-RA      GGACCGGTCGTCGATGCACTCGAGGTG-CACCGCATTGACGAGCGGCAGTAGCAGGATGC 59
AGAP012700-RA      GGACCGGTCGTCGATGCACTCGAGTTG-CACCGCATTGACGAGCGGCAGTAGCAGGATGC 59
AGAP012752-RA      GGACCGGTCGTCGATGCACTCGAGGTGTCACCGCATTGACGAGCGGCAGTAGCAGGATGC 60
AGAP001087-RA      GGACCGGTCGTCGATGCACTCGAGGTG-CACCGCATTGACGAGCGGCAGTAGCAGGATGC 59
                   ************************ ** ********************************

AGAP012483-RA      AATCGGCGCGGCCGAGCTGGCAGCGAAGCCGGTTCTCGGTGAGCCATTCACGCCGAATGA 119
AGAP012762-RA      AATCGGCGCGGCCGAGCTGGCAGCGAAGCCGGTTCTCGGTGAGCCATTCACGCCGAATGA 119
AGAP012700-RA      AATCGGCGCGGCCGAGCTGGCAGCGAAGCCGGTTCTCGGTGAGCCATTCACGCCGAATGA 119
AGAP012752-RA      AATCGGCGCGGCCGAGCTGGCAGCGAAGCCGGTTCTCGGTGAGCCATTCACGCCGAATGA 120
AGAP001087-RA      AATCGGCGCGGCCGAGCTGGCAGCGAGGCCGTTTCTCGGTGAGCCATTCACGCCGAATGA 119
                   ************************** **** ****************************

AGAP012483-RA      GGGCGGTCAAGAGCCGGCTGCATACGTTCGAT------------ATATTCGCTACGCGCA 167
AGAP012762-RA      GGGCGGTCAAGAGCCGGCTGCATACGTTCGAT------------ATATTCGCTACGCGCA 167
AGAP012700-RA      GGGCGGTCAAGAGCCGGCTGCATACGTTCGAT------------ATATTCGCTACGCGCA 167
AGAP012752-RA      GGGCGGTCAAGAGCCGGCTGCATACGTTCGAT------------ATATTCGCTACGCGCA 168
AGAP001087-RA      GGGCGGTCAAGAGCCGGCTGCATACGTTCGATTCGTTCGATTCGATATTCGTTACGCGCA 179
                   ********************************            ******* ********

AGAP012483-RA      CGATCAAACTGTGTAACGATGCCGAGTGTGTCCGGCCATCGCGACTAGAAGCATAGCGGA 227
AGAP012762-RA      CGATCAAACTGTGTAACGATGCCGAGTGTGTCCGGCCATCGCGACTAGAAGCATAGCGGA 227
AGAP012700-RA      CGATCAAACTGTGTAACGATGCCGAGTGTGTCCGGCCATCGCGACTAGAAGCATAGCGGA 227
AGAP012752-RA      CGATCAAACTGCGTAACGATGCCGAGTGTGTCCGGCCATCGCGACTAGAAGCATAGCGGA 228
AGAP001087-RA      CGATCAAACTGTGTAACGATGCCGAGTGTGTCCGGCCATCGCGACTAGAAGCATAGCAAA 239
                   *********** *********************************************  *

AGAP012483-RA      AGTTAAGTTGCGTCGTCTATTGTGGTGCGATGCGCTGTATGTTCCGGGTGTGCGATTGAG 287
AGAP012762-RA      AGTTAAGTTGCGTCGTCTATTGTGGTGCGATGCGCTGTATGTTCCGGGTGTGCGATTGAG 287
AGAP012700-RA      AGTTAAGTTGCGTCGTCTATTGTGGTGCGATGCACTGTATGTTCCGGGTGTGCGATTGAG 287
AGAP012752-RA      AGTTAAGTTGCGTCGTCTATTGTGGTGCGATGCACTGTATGTTCCGGGTGTGCGATTGAG 288
AGAP001087-RA      AGTTAAGTTGCGTCGTCTATTGTGGTACGATGCGCTGTATGTTCCGGGTGTGCGATTGAG 299
                   ************************** ****** **************************

AGAP012483-RA      TTTCATGTTTGTTTCTTTCTTGTTCACAGACACCTTTTCATCAACAACTGGCATGACTAT 347
AGAP012762-RA      TTTCATGTTTGTTTCTTTCTTGTTCACAGACACCTTTTCATCAACAACTGGCATGACTAT 347
AGAP012700-RA      CTTCATGTTTGTTTCTTTTTTGTTCACAGACACCTTTTCATCAACAACTGGCATGACTAT 347
AGAP012752-RA      CTTCATGTTTGTTTCTTTTTTGTTCACAGACACCTTTTCATCAACAACTGGCATGACTAT 348
AGAP001087-RA      TTTCATGTTTGTTTCTTTCTTGTTCACAGACACCTTTTCATCAACAACTGGCATGACTAT 359
                    ***************** *****************************************

AGAP012483-RA      CGGCGCACGGTCGGTGGCTCTAGCGAAGGGGAGGGTGCGTTCCAATCGGCTGGCAGGACA 407
AGAP012762-RA      CGGCGCACGGTCGGTGGCTCTAGCGAAGGGGAGGGTGCGTTCCAATCGGCTGGCAGGACA 407
AGAP012700-RA      CGGCGCACGGTCGGTGGCTCTAGCGAAGGGGAGGGTGCGTTCCAATCGGCTGGCAGGACA 407
AGAP012752-RA      CGGCGCACGGTCGGTGGCTCTAGCGAAGGGGAGGGTGCGTTCCAATCGGCTGGCAGGACA 408
AGAP001087-RA      CGGCGCTCGGTCGGTGGCTCTAGCGAAGGGGAGGGTGCGTTCCAATCGGCTGGCAGGACA 419
                   ****** *****************************************************

AGAP012483-RA      CAGATCGGATCGGTTAGGACAACGATCGGCATAGTACGGCTCGCGACCCGCATGTGGCTA 467
AGAP012762-RA      CAGATCGGATCGGTTAGGACAACGATCGGCATAGTACGGCTCGCGACCCGCATGTGGCTA 467
AGAP012700-RA      CAGATCGGATCGGTTAGGACAACGATCGGCATAGTACGGCTCGCGACCCGCATGTGGCTA 467
AGAP012752-RA      CAGATCGGATCGGTTAGGACAACGATCGGCATAGTACGGCTCGCGACCCGCATGTGGCTA 468
AGAP001087-RA      CAGATCGGATCGGTTAGAACAGCGATCGGCATAGTACGGCTCGCGACCCGCTTGTGGCTA 479
                   ***************** *** ***************************** ********

AGAP012483-RA      TTTGTTGACTAGTTGCCTCTTCAAGTGCAAATCTACGAACATGTTTAGAAGATTTGGCTC 527
AGAP012762-RA      TTTGTTGACTAGTTGCCTCTTCAAGTGCAAATCTACGAACATGTTTAGAAGATTTGGCTC 527
AGAP012700-RA      TTTGTTGACTAGTTGCCTCTTCAAGTGCAAATCTACGAACATGTTTAGAAGATTTGGCTC 527
AGAP012752-RA      TTTGTTGACTAGTTGCCTCTTCAAGTGCAAATCTACGAACATGTTTAGAAGATTTGGCTC 528
AGAP001087-RA      TTTGTTGACTAGTTGCCTCTTCAAGTGCAAATCTACGAACATGTTTAGAAGACTTGGCTC 539
                   **************************************************** *******

AGAP012483-RA      TTTCGGTCGCAACGTTTGCCGAGTCGCTGGGGATATTGGATTGTAAGGAATGAATATACT 587
AGAP012762-RA      TTTCGGTCGCAACGTTTGCCGAGTCGCTGGGGATATTGGATTGTAAGGAATGAATATACT 587
AGAP012700-RA      TTTCGGTCACAACGTTTGCCGAGTCGCTGGGGATATTGGATTGTAAGGAATGAATATACT 587
AGAP012752-RA      TTTCGGTCGCAACGTTTGCCGAGTCGCTGGGGATATTGGATTGTAAGGAATGAATATACT 588
AGAP001087-RA      TTTCGGTCGCAACGTTTGCCGAGTCGCTGGGGATATTGGATTGTAAGGAATGAATATACT 599
                   ******** ***************************************************

AGAP012483-RA      ATCCGCAGGAACCAGATGATGACATCACCAAGGAGCTAATGCGCTAAAGCTTATCAGAAG 647
AGAP012762-RA      ATCCGCAGGAACCAGATGATGACATCACCAAGGAGCTAATGCGCTAAAGCTTATCAGAAG 647
AGAP012700-RA      GTCCGCAGGAACCAGATGATGACATCACCAAGGAGCTAATGCGCTAAAGCTTATCAGGAG 647
AGAP012752-RA      GTCCGCAGGAACCAGATGATGACATCACCAAGGAGCTAATGCGCTAAAGCTTATCAGGAG 648
AGAP001087-RA      GTCCGCAGGAACCAGATGATGACATCACCAAGGAGCTAATGCGCTAAAGCTTATCAGGAG 659
                    ******************************************************** **
```

```
AGAP012483-RA       GAGGCGAGAATGGTTTGATGT 668
AGAP012762-RA       GAGGCGAGAATGGTTTGATGT 668
AGAP012700-RA       GAGGCGAGAATGGTTTGATGT 668
AGAP012752-RA       GAGGCGAGAATGGTTTGATGT 669
AGAP001087-RA       TAGGCGAGAATGGTTTGATGT 680
                     ********************
```

Ping-pong network 2:
Sequence Logo 3.5.0 (toolshed.g2.bx.psu.edu/repos/devteam/weblogo3/rgweblogo3/3.5.0)

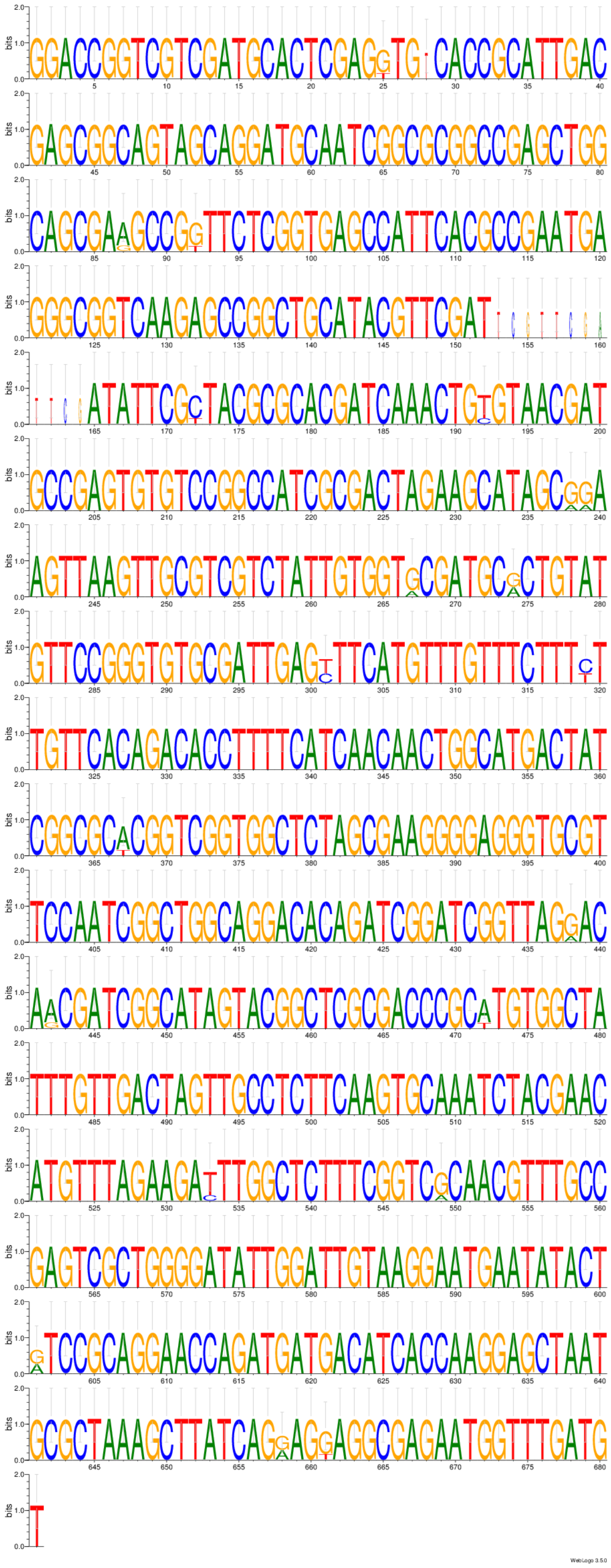

Ping-pong network 3:
CLUSTAL 2.1 multiple sequence alignment

```
AGAP013070-RA      ------------------------------------------------------------
AGAP013323-RA      ------------------------------------------------------------
AGAP013502-RA      ------------------------------------------------------------
AGAP013002-RA      ------------------------------------------------------------
AGAP013534-RA      ------------------------------------------------------------
AGAP013334-RA      ------------------------------------------------------------
AGAP013312-RA      TAGCTTTTGAATACAAACTGAAAAGGAAACACAAACCATCAAAAAGAAAAAACGATTGGT 60
AGAP000976-RA      ------------------------------------------------------------


AGAP013070-RA      ------------------------------------------------------------
AGAP013323-RA      ------------------------------------------------------------
AGAP013502-RA      ------------------------------------------------------------
AGAP013002-RA      ------------------------------------------------------------
AGAP013534-RA      ------------------------------------------------------------
AGAP013334-RA      ------------------------------------------------------------
AGAP013312-RA      GATATCACGAAAACCAACACACTGTGCATCAATCAGAAAGATCTCAACCACAACATATAA 120
AGAP000976-RA      ------------------------------------------------------------


AGAP013070-RA      ------------------------------------------------------------
AGAP013323-RA      ------------------------------------------------------------
AGAP013502-RA      ------------------------------------------------------------
AGAP013002-RA      ------------------------------------------------------------
AGAP013534-RA      ------------------------------------------------------------
AGAP013334-RA      ------------------------------------------------------------
AGAP013312-RA      ACATCAACGAAACAGCAATATAAACATCAAAGGAGAATGGATGATTTCCCCTCCAAAAAC 180
AGAP000976-RA      ------------------------------------------------------------


AGAP013070-RA      --------------------------------------------------------ATGT 4
AGAP013323-RA      --------------------------------------------------------ATGT 4
AGAP013502-RA      ------------------------------------------------------------
AGAP013002-RA      ------------------------------------------------------------
AGAP013534-RA      ------------------------------------------------------------
AGAP013334-RA      ------------------------------------------------------------
AGAP013312-RA      TGGTAGTCAAACCTCTGATTGTACCTTCAAAAACGGATGGTATAGGAGGCCTCAACCTAC 240
AGAP000976-RA      ------------------------------------------------------------


AGAP013070-RA      CAAAAAGTTCGTCAAGCTTCGTGTGTGATGAAAGGCGACAAGAAAAGCAAGCAGCCGCAA 64
AGAP013323-RA      CAAAAAGTTCGTCAAGCTTCGTGTGTGATGAAAGGCGACAAGAAAAGCAAGCAGCCGCAA 64
AGAP013502-RA      ---------CGTCAAGCTTCGTGTGTGATGAAAGGCGACAAGAAAAGCAAGCAGCCGCAA 51
AGAP013002-RA      ---------CGTCAAGCTTCGTGTGTGATGAAAGGCGACAAGAAAAGCAAGCAGCCGCAA 51
AGAP013534-RA      ---------CGTCAAGCTTCGTTTGTGATGAAAGGCCACAAGAAAAGCAAGCATCCGCAA 51
AGAP013334-RA      ---------CGTCAAGCTTCGTGTGTGATGAAAGGCGACAAGAAAAGCAAGCAGCCGCAA 51
AGAP013312-RA      CAACCCTATTAAGAAGAAGAAGAAGTGATGAAAGGCGACAAGAAAAGCAAGCAGCCGCAA 300
AGAP000976-RA      ---------------------------------GGCGACAAGAAAAGCAAGCAGCCTCAA 27
                                                    *** **************** ** ***

AGAP013070-RA      ATTTTACAGCTTGTAATTTGTGTATTATCAAGCTTGTACAACAGCATTTAAACTCGGTGG 124
AGAP013323-RA      ATTTTACAGCTTGTAATTTGTGTATTATCAAGCTTGTACAACAGCATTTAAACTCGGTGG 124
AGAP013502-RA      ATTTTACAGCTTGTAATTTGTGTATTATCAAGCTTGTACAACAGCATTTAAACTCGGTGG 111
AGAP013002-RA      ATTTTACAGCTTCTAATTTGTATATTATCAAGCTTGTACAACAGCATTTAAACTCGGTGG 111
AGAP013534-RA      ATTTTACAGCTTCTAATTTGTATATTATCAAGCTTGTACAACAGCATTTAAACTCGGTGG 111
AGAP013334-RA      ATTTTACAGCTTGTAATTTGTATATTATCAAGCTTGTACAACAGCATTTAAACTCGGTGG 111
AGAP013312-RA      ATTTTACAGCTTGTAATTTGTGTATTATCAAGCTTGTACAACAGCATTTAAACTCGGTGG 360
AGAP000976-RA      ATTTAACAGCTTGTAATTTGTGTATTATCAAGCTTATACAACAGCATTTAAACTCGGTGG 87
                   **** ******* ******** ************* ************************

AGAP013070-RA      TAGAGTTCCGCCCGCTCCATCAAATACAGAGAATGACGCCGTCCAATATTGTTCCTGCCT 184
AGAP013323-RA      TAGAGTTCCGCCCGCTCCATCAAATACAGAGAATGACGCCGTCCAATATTGTTCCTGCCT 184
AGAP013502-RA      TAGAGTTCCGCCCGCTCCATCAAATACAGAGAATGACGCCGTCCAATATTGTTCCTGCCT 171
AGAP013002-RA      TAGAGTTCCGCCCGCTCCATCAAATACAGAGAATGACGCCGTCCAATATTGTTCCTGCCT 171
AGAP013534-RA      TAGAGTTCCGCCCGCTCCATCAAATACAGAGAATGACGCCGTCCAATATTGTTCCTGCCT 171
AGAP013334-RA      TAGAGTTCCGCCCGCTCCATCAAATACAGAGAATGACGCCGTCCAATATTGTTCCTGCCT 171
AGAP013312-RA      TAGAGTTCCGCCCGCTCCATCAAATACAGAGAATGACGCCGTCCAATATTGTTCCTGCCT 420
AGAP000976-RA      TAGAGTTCCGCCCGCTCCATCAAATGTAGAGAATGACGCTGTCCAATATTGTTCCTGCCT 147
                   *************************  ************ ********************

AGAP013070-RA      CGCGGTCGTGCACTACCGTCAGCACTGCACTAGCCGCAACAAAACATGCGCCCCTTGTCC 244
AGAP013323-RA      CGCGGTCGTGCACTACCGTCAGCACTGCACTAGCCGCAACAAAACATGCGCCCCTTGTCC 244
AGAP013502-RA      CGCGGTCGTGCACTACCGTCAGCACTGCACTAGCCGCAACAAAACATGCGCCCCTTGTCC 231
AGAP013002-RA      CGCGGTCGTGCACTACCGTCAGCACTGCACTAGCCGCAACAAAACATGCACCCCTTGTCC 231
AGAP013534-RA      CGCGGTCGTGCACTACCGTCAGCACTGCACTAGCCGCAACAAAACATGCGCCCCTTGTCC 231
AGAP013334-RA      CGCGGTCGTGCACTACCGTCAGCACTGCACTAGCCGCAACAAAACATGCGCCCCTTGTCC 231
AGAP013312-RA      CGCGGTCGTGCACTACCGTCAGCACTGCACTAGCCGCAACAAAACATGCGCCCCTTGTCC 480
AGAP000976-RA      CGCGGTCGTGCACTACCGTCAGCACTGCAATAGCCGCAGCAAAACATGCGCCCCATGTCC 207
                   ***************************** ******** ********** **** *****
```

```
AGAP013070-RA      CACATTTCCATAATCTCGACGGTGGTCAACAGCTGTACGACGGCCGTCTACAGCCACCGG 304
AGAP013323-RA      CACATTTCCATAATCTCGACGGTGGTCAACAGCTGTACGACGGCCGCCTACAGCCACCGG 304
AGAP013502-RA      CACATTTCCATAATCTCGACGGTGGTCAACAGCTGTACGACGGCCGCCTACAGCCACCGG 291
AGAP013002-RA      CACATTTCCATAATCTCGACGGTGGTCAACAGCTGTACGACGGCCGCCTACAGCCACCGG 291
AGAP013534-RA      CACATTTCCATAATCTCGACGGTGGTCAACAGCTGTACGACGGCCGCCTACAGCCACCGG 291
AGAP013334-RA      CACATTTCCATAATCTCGACGGTGGTCAACAGCTGTACGACGGCCGCCTACAGCCACCGG 291
AGAP013312-RA      CACATTTCCATAATCTCGACGGTGGTCAACAGCTGTACGACGGCCGCCTACAGCCACCGG 540
AGAP000976-RA      CACATTTCCATCATCTCGACGGTGCTCAACAGCTGTACGCCTGCTGCCTACAGCCACCGG 267
                   *********** ************ ************** * ** * *************

AGAP013070-RA      GATAGCCTCGCTCTCTAGTGCGATACTGATGAACCTGCGGCGCAAGAGCACGCAAGCGTT 364
AGAP013323-RA      GATAGCCTCGCTCTCTAGTGCGATACTGATGAACCTGCGGCGCAAGAGCACGCAAGCGTT 364
AGAP013502-RA      GATAGCCTCGCTCTCTAGTGCGATACTGATGAACCTGCGGCGCAAGAGCACGCAAGCGTT 351
AGAP013002-RA      GATAGCCTCGCTCTCTAGTGCGATACTGATGAACCTGCGGCGCAAGAGCACGCAAGCGTT 351
AGAP013534-RA      GATAGCCTCGCTCTCTAGTGCGATACTGATGAACCTGCGGCGCAAGAGCACGCAAGCGTT 351
AGAP013334-RA      GATAGCCTCGCTCTCTAGTGCGATACTGATGAACCTGCGGCGCAAGAGCACGCAAGCGTT 351
AGAP013312-RA      GATAGCCTCGCTCTCTAGTGCGATACTGATGAACCTGCGGCGCAAGAGCACGCAAGCGTT 600
AGAP000976-RA      GATAGCCTCGCTCTCTAGTGTGATACTGACGAACCTGCGGCGCAAGAGCACGTAAGCGTT 327
                   ******************** ******** ********************** *******

AGAP013070-RA      TGGTGGCGGCGGCTGTGGAATGATGGTGCACCTCGGTAAATATGGTGCACGAGAAGAATT 424
AGAP013323-RA      TGGTGGCGGCGGCTGTGGAATGATGGTGCACCTCGGTAAATATGGTGCACGAGAAGAATT 424
AGAP013502-RA      TGGTGGCGGCGGCTGTGGAATGATGGTGCACCTCGGTAAATATGGTGCACGAGAAGAATT 411
AGAP013002-RA      TGGTGGCGGCGGCTGTGGAATGATGGTGCACCTCGGTAAATATGGTGCACGAGAAGAATT 411
AGAP013534-RA      TGGTGGCGGCGGCTGTGGAATGATGGTGCACCTCGGTAAATATGGTGCACGAGAAGAATT 411
AGAP013334-RA      TGGTGGCGGCAGCTGTGGAATGATGGTGCACCTCGGTAAATATGGTGCACGAGAAGAATT 411
AGAP013312-RA      TGGTGGCGGCGGCTGTGGAATGATGGTGCACCTCAGTAAATATGGTGCACGAGAAGAATT 660
AGAP000976-RA      TGGTGGCGGCGGCTGTGGAATGATGGTGCACCTCGGTAAATATGGTGCACGAGAAGAATT 387
                   ********** *********************** *************************

AGAP013070-RA      CTTCCCGCAAAGGGCATCCTTTTTTGGGCACTGAAATCGAACGACGCCTACGGAGACAGT 484
AGAP013323-RA      CTTCCCGCAAAAGGCATCCTTTT--GGGCACTGAAATCGAACGACGCCTACGGAGACAGT 482
AGAP013502-RA      CTTCCCGCAAAAGGCATCCGTTT--GGGCACTGAAATCGAACGACGCCTACGGAGACAGT 469
AGAP013002-RA      CTTCCCGCAAAAGGCATCCGCTT--GGGCACTGAAATCGAACGACGCCTACGGAGACAGT 469
AGAP013534-RA      CTTCCCGCAAAAGGCATCCGCTT--GGGCACTGAAATCGAACGACGCCTACGGAGACAGT 469
AGAP013334-RA      CTTCCCGCAAAAGGCATCCGCTT--GGGCACTGAAATCGAACGACGCCTACGGAGACAGT 469
AGAP013312-RA      CTTCCTGCAAAAGGCATTCGCTT--GGGCACTGAAATCGAACGACGCCTACGGAGACAGT 718
AGAP000976-RA      CTTCCCGCAAAAGGCATCCGTTT--GGGCACTGAAATCGAACGACGTCTACGGAGACAGT 445
                   ***** ***** ***** *  **  ******************** *************

AGAP013070-RA      AACAACAACAACCATCCGGACAGGAGACGGATCTCAGTCGGTTGTATGAAAACAGCA--- 541
AGAP013323-RA      AACAACAACAACCATCCGGACAGGAGACGGATCTCAGTCGGTTGTATGAAAACAGCA--- 539
AGAP013502-RA      AACAACAACAACCATCCGGACAGGAGACGGATCTCAGTCGGTTGTATGATAACAGCA--- 526
AGAP013002-RA      AACAACAACAACCATCCGGACAGGAGACGGATCTCAGTCGGTTGTATGAAAACAGCA--- 526
AGAP013534-RA      AACAACAACAACCATCCGGACAGGAGACGGATCTCAGTCGGTTGCATGAAAACAGCA--- 526
AGAP013334-RA      AACAACAACAACCATCCGGACAGGAGACGGATCTCAGTCGGTTGTATGAAAACAGCA--- 526
AGAP013312-RA      AACAACAACAACCATCCGGACAGGAGACGGATCTCAGTCGGTTGTATGAAAACAGCA--- 775
AGAP000976-RA      AACAACAACAACCATCCGGACAGGAGGTGGATCTCAGTCGGTTGTATAAAAACAGCATCA 505
                   **************************  *************** ** * *******

AGAP013070-RA      AGGTGCACTAGATCGTGAAACACTGGCCGTCCAGTACGACACAGGCCTTATCAGCAGATG 601
AGAP013323-RA      AGGTGCACTAGATCGTGAAACACTGGCCGTCCAGTACGACACAGGCCTTATCAGCAGATG 599
AGAP013502-RA      AGGTGCACTAGATCGTGAAACACTGGCCGTCCAGTACGACACAGGCCTTATCAGCAGATG 586
AGAP013002-RA      AGGTGCACTAGATCGTGAAACACTGGCCGTCCAGTACGACACAGGCCTTATCAGCAGATA 586
AGAP013534-RA      AGGTGCACTAGATCGTGAAACACTGGCCGTCCAGTACGACACAGGCCTTATCAGCAGATA 586
AGAP013334-RA      AGGTGCACTAGATCGTGAAACACTGGCCGTCCAGTACGACACAGGCCTTATCAGCAGATA 586
AGAP013312-RA      AGGTGCACTAGATCGTGAAACACTGGCCGTCCAGTACGACACAGGCCTTATCAGCAGATA 835
AGAP000976-RA      AGGTGCACCAGATCGCGAAACACGGGCCGTCCAGTACGACACAGGCCTTATCGGCAG--- 562
                   ******** ****** ******* **************************** ****

AGAP013070-RA      GCATGGAAATGCCGCGTACAATGAGGCCGCACCGTGACGCCCCGCGCTGTAGCCCTCGTT 661
AGAP013323-RA      GCATGGAAATGCCGCGTACAATGAGGCCGCACCGTGACGCCCCGCGCTGTAGCCCTCGTT 659
AGAP013502-RA      GCATGGAAATGCCGCGTACAATGAGGCCGCACCGTGACGCCCCGCGCTGTAGCCCTCGTT 646
AGAP013002-RA      GCATGGAAATGCCGCGTACAATGAGGCCGCACCGTGACGCCCCGCGCTGTAGCCCTCGTT 646
AGAP013534-RA      GCATGGAAATGCCGCGTACAATGAGGCCGCACCGTGACGCCCCGCGCTGTAGCCCTCGTT 646
AGAP013334-RA      GCATGGAAATGCCGCGTACAATGAGGCCGCACCGTGACGCCCCGCGCTGTAGCCCTCGTT 646
AGAP013312-RA      GCATGGAAATGCCGCGTACAATGAGGCCGCACCGTGACGCCCCGCGCTGTAGCCCTCGTT 895
AGAP000976-RA      -CCTGGAG--GCCTTAT----TGGAGTCCCGCC-TTATGTCCTG--CCGTAGGATCCAAC 612
                    * ****   ***    *    **  * * * ** * * * ** *  * ****    *

AGAP013070-RA      TACGCACCGAAGTTAAGCAGTTCCATCGGGATGTCTTCACCGCGCCGGATCATTTGCGGG 721
AGAP013323-RA      TACGCACCGAAGTTAAGCAGTTCCATCGGGATGTCTTCACCGCGCCGGATCATTTGCGGG 719
AGAP013502-RA      TACGCACCGAAGTTAAGCAGTTCCATCGGGATGTCTTCACCGCGCCGGATCATTTGCGTG 706
AGAP013002-RA      TACGCACCGAAGTTAAGCAGTTCCATCGGGATGTCTTCACCGCGCCGGATCATTTGCGGG 706
AGAP013534-RA      TACGCACCGAAGTTAAGCAGTTCCATCGGGATGTCTTCACCGCGCCGGATCATTTGCGGG 706
AGAP013334-RA      TACGCACCGAAGTTAAGCAGTTCCATCGGGATGTCTTCACCGCGCCGGATCATTTGCGGG 706
AGAP013312-RA      TACGCACCGAAGTTAAGCAGTTCCATCGGGATGTCTTCACCGCGCCGGATCATTTGCGGG 955
AGAP000976-RA      CAA--------------------------------------------------------- 615
                    *
```

```
AGAP013070-RA      ACGATACCCA-------------------------------------------------- 731
AGAP013323-RA      ACGATAGCGACAAATGTCCCGCCGTAGGATCCAACCAATATGGTGGATAGTTAATATAAA 779
AGAP013502-RA      ACGATAGCGACAAATGTCCCGCCGTAGGATCCAACCAATATAGTGGATAATTAATATAAA 766
AGAP013002-RA      ACGATAGCGACAAATGTCCCGCCGTAGGATCCAACCAATATGGTGGATAATTAATATAAA 766
AGAP013534-RA      ACGATAGCGACAAATGTCCCGCCGTAGGATCCAACCAATATGGTGGATAATTAATATAAA 766
AGAP013334-RA      ACGATAGCGACAAATGTCCCGCCGTAGGATCCAACCAATATGGTGGATAATTAATATAAA 766
AGAP013312-RA      ACGATAGCGACAAATGTTCCGCCGTAGGATCCAACCAATATGGTGGATAATTAATATAAA 1015
AGAP000976-RA      ------------------------------------------------------------

AGAP013070-RA      ---------------------
AGAP013323-RA      AAATACAAATTACATTAATTT 800
AGAP013502-RA      AAATACAAATTACATTAATTT 787
AGAP013002-RA      AAATACAAATTACATTAATTT 787
AGAP013534-RA      AAATACAAATTACATTAATTT 787
AGAP013334-RA      AAATACAAATTACATTAATTT 787
AGAP013312-RA      AAATACAAATTTCAATAATTT 1036
AGAP000976-RA      ---------------------
```

Ping-pong network 3:
Sequence Logo 3.5.0 (toolshed.g2.bx.psu.edu/repos/devteam/weblogo3/rgweblogo3/3.5.0)

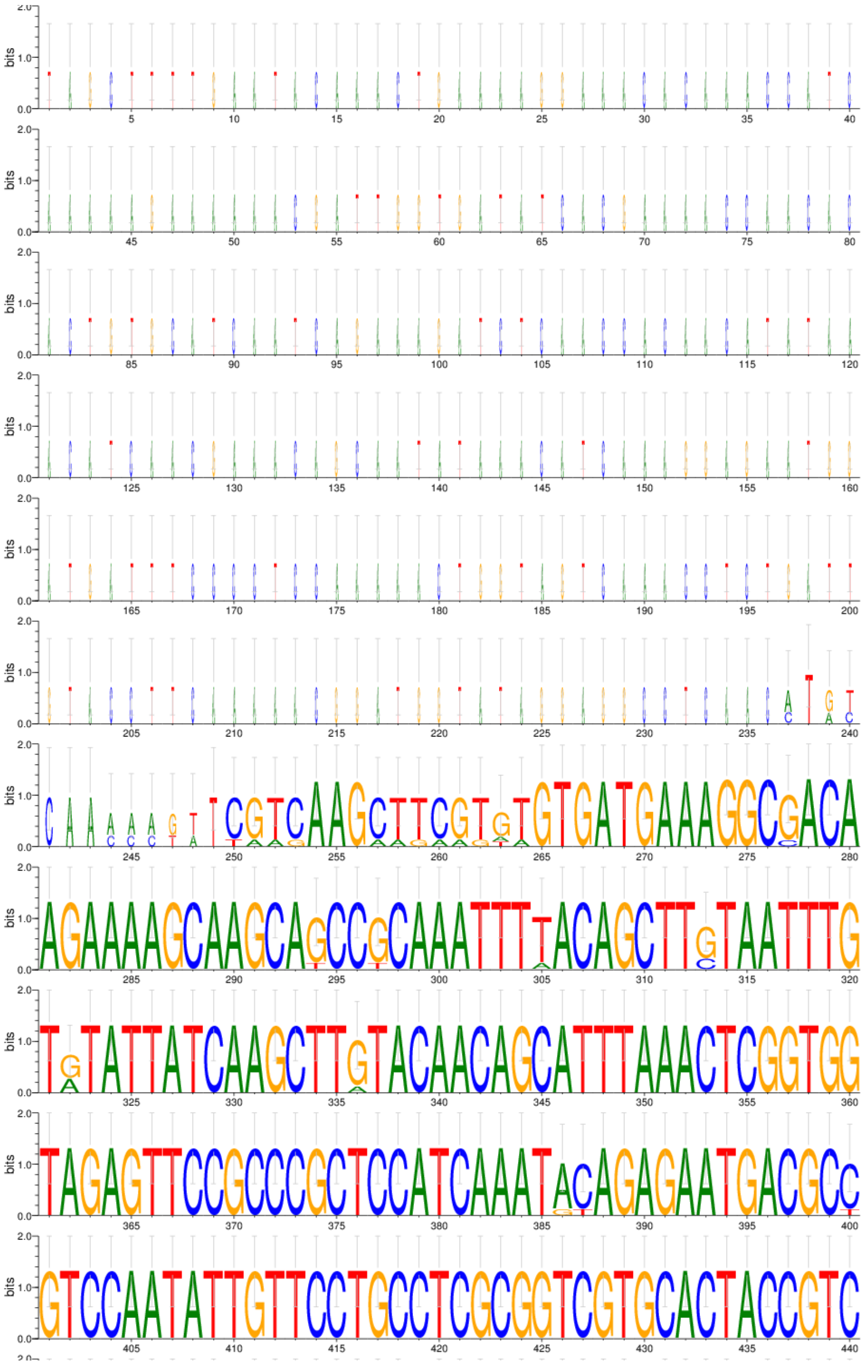

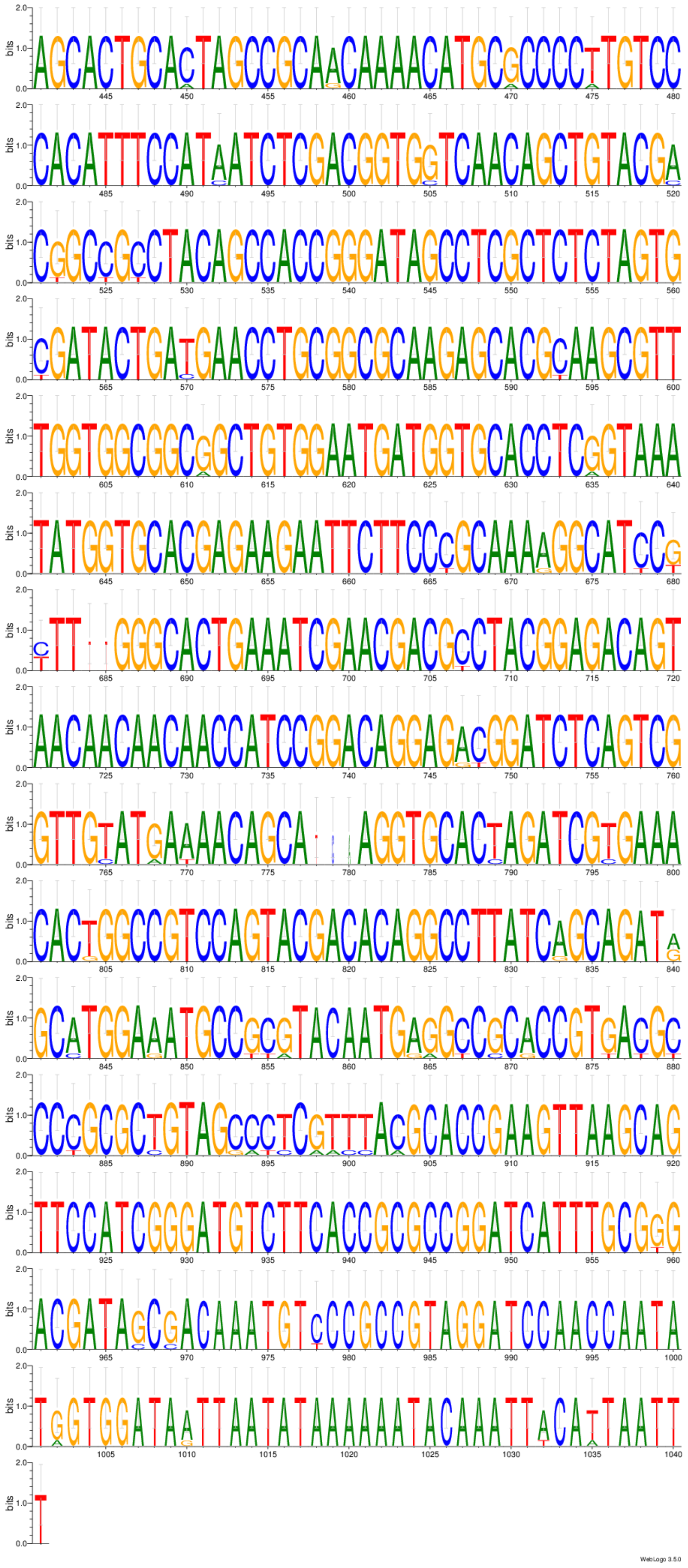
WebLogo 3.5.0

Ping-pong network 4:
CLUSTAL 2.1 multiple sequence alignment

```
AGAP012506-RA      TGATTTCTTTTGCTTCGAACTTAAAGTTGTTTTTCTTATTTGTTTTGCATAAAACCAAGT 60
AGAP012538-RA      TGATTTCTTTTGCTTCGAACTTAAAGTTGTTTTTCTTATTTGTTTTGCATAAAACCAAGT 60
AGAP011660-RA      TGATTTCTTTTGCTTCGAACTTAAAGTTGCTTTTTTTATTTGTTTTGCATAAAACCAAGT 60
AGAP012490-RA      TGATTTCTTTTGCTTCGAACTTAAAGTTGTTTTTTTTATTTGTTTTGCATAAAACCAAGT 60
                   ***************************** **** *************************

AGAP012506-RA      GAAAATGGATCCTTCCAAACGCTTGAAGAAAAGTGGAGTGGCGAGCAAGCGTGCCAATAG 120
AGAP012538-RA      GAAAATGGATCCTTCCAAACGCTTGAAGAAAAGTGGAGTGGCGAGCAAGCGTGCCAA-AG 119
AGAP011660-RA      GAAAATGGATACGTCCAAACGTTTGAAGAAAAGTGGAGTGGCGAGCAAGCGTTCCAA-AG 119
AGAP012490-RA      GAAAATGGATACGTCTAAACGTTTGAAGAAAAGTGGAGTGGCGAGCAAGCGTACCAA-AG 119
                   ********** * ** ***** ****************************** **** **

AGAP012506-RA      CACATTTGGGAGCGCTTTGAAAAGGAATGGGAAGAGGAAAATCTTCCGGGAACAAGCAGC 180
AGAP012538-RA      CA-ATTTGGGAGCGCTTTGAAAAGGAATGGGAAGAGGAAAATCTTCCGGGAACAAGCAGC 178
AGAP011660-RA      CA-ATTTGGGAGCGCTTTGAAAAGGAATGGGAAGAGGAAAATCTTCCGGGAACAAGCAGC 178
AGAP012490-RA      CA-ATTTGGGAGCGCTTTGAAAAGGAATGGGAAGAGGAAAATCTTCCGGGAACAAGCAGC 178
                   ** *********************************************************

AGAP012506-RA      GGTGTCGATCAACAAGCATCATCATACGGGAATGTGCAGCCGGATGCCGTGGAAGTTGCC 240
AGAP012538-RA      GGTGTCGATCAACAAGCATCATCATACGGGAATGTGCAGCCGGATGCCGTGGAAGTTGCC 238
AGAP011660-RA      GGTGTCGATCAACAAGCTTCATCATACGGGAATGTGCAGCCGGATGCCGTGGAAGTTGCC 238
AGAP012490-RA      GATGTCGATCAACAAGCTTCATCATACGGGAATGTGCAGCCGGATGCCGTGGAAGTTGCC 238
                   * *************** ******************************************

AGAP012506-RA      GATATGCCCTTGGAATCAGAAGCAGCGGATTACGTTTCTGATGACGATGAAGACGCTGAT 300
AGAP012538-RA      GATATGCCCTTGGAATCAGAAGCAGCGGATTACGTTTCTGATGACGATGAAGACGCTGAT 298
AGAP011660-RA      GATATACCCATGGAATCAGAAGCAGCGGATTACGTTTCTGATGACGATGAAGACGCTGAT 298
AGAP012490-RA      GATATACCCATGGAATCAGAAGCAGCGGATTACGTTTCTGATGACGATGAAGACGCTGAT 298
                   ***** *** **************************************************

AGAP012506-RA      TGCTCGGTGTTGGAAGACGATTGGAGTGAAGGCGTATTGGAGGACGAAGTCAATGAGGAT 360
AGAP012538-RA      TGCTCGGTGTTGGAAGACGATTGGAGTGAAGGCGTATTGGAGGACGAAGTCAATGAGGAT 358
AGAP011660-RA      TGCTCGGTGTTGGAAGACGATTGGAGTGAAGGCGAATTGGAGGACGAAGTCAATGAGGAT 358
AGAP012490-RA      TGCTCGGTGTTGGAAGACGATTGGAGTGAAGGCGAATTGGAGGACGAAGTCAATGAGGAT 358
                   ********************************** *************************

AGAP012506-RA      GAGTATTTTGATGCTGATGAAGCCCCCGAAGGCAATGCTTATGCTAGCAGGCTTAGGATA 420
AGAP012538-RA      GAGTATTTTGATGCTGATGAAGCCCCCGAAGGCAATGCTTATGCTAGCAGGCTTAGGATA 418
AGAP011660-RA      GAGTATTTTGATGCTGATGAAGCCCCCGAAGGCAATGCTTATGCTAGCAGGCTTAGGATA 418
AGAP012490-RA      GAGTATTTTGATGCTGATGAAGCCCCCGAAGGCAATGCTTATGCTAGCAGGCTTAGGATA 418
                   ************************************************************

AGAP012506-RA      TGGGCTTTAACTCATAAAATAACGCATTCTGCATTGAGTGATTTGCTGGTGTTGACTCGT 480
AGAP012538-RA      TGGGCTTTAACTCATAAAATAACGCATTCTGCATTGAGTGATTTGCTGGTGTTGACTCGT 478
AGAP011660-RA      TGGGCTTTAACTCATAAAATAACGCATTCTGCATTGAGTGATTTGCTGGTGTTGACTCGT 478
AGAP012490-RA      TGGGCTTTAACTCATAAAATAACGCATTCAGCATTGAGTGATTTGCTGGTGTTGACTCGT 478
                   ***************************** ******************************

AGAP012506-RA      GAAACTACCAATATTTCTCTTCCTAGGTGTGCAAAGACGCTACTGAAGACTCCGAAACAG 540
AGAP012538-RA      GAAACTACAAATATTTCTCTTCCTAGGTGTGCAAAGACGCTACTGAAGACTCCGAAACAG 538
AGAP011660-RA      GAAACTACAAATATTTCTCTTCCTAGGTGTGCAAAGACGTTACTGAAGACTCCGAAACAG 538
AGAP012490-RA      GAAACTACAAATATTTCTCTTCCTAGGTGTGCAAAGACGTTACTGAAGACTCCGAAACAG 538
                   ******** ****************************** ********************

AGAP012506-RA      GTGGAGAGACATTTTACGGCCGTTGGAGAAGGGCAGCTTTGGTACCAAGGAATACAAAGC 600
AGAP012538-RA      GTGGAGAGACATTTTACGGCCGTTGGAGAAGGGCAGCTTTGGTACCAAGGAATACAAAGC 598
AGAP011660-RA      GTGGAGAGACATTTTACGGCCGTTGGAGAAGGGCAGCTTTGGTACCAAGGAATACAAAGC 598
AGAP012490-RA      GTGGAGAGACATTTTACGGCCGTTGGAGAAGGGCAGCTTTGGTACCAAGGAATACAAAGC 598
                   ************************************************************

AGAP012506-RA      ACTCTCCAACAATATTATCGATCTGTACCACCAGTCATCAGATTGATAGAAGCTGATATC 660
AGAP012538-RA      ACTCTCCAACAATATTATCGATCTGTACCACCAGTCATCAGATTGATAGAAGCTGATATC 658
AGAP011660-RA      ACTCTCCAACAATATTATCGATCTGTACCACCAGTCATCAGATTGATAGAAGCTAATATC 658
AGAP012490-RA      ACTCTCCAACAATATTATCGATCTGTACCACCAGTCATCAGATTGATAGAAGCTAATATC 658
                   ****************************************************** *****

AGAP012506-RA      GCGGTAGATGGACTTCCGATGCATAATAGTGGACCAACGCAGCTATGGCCTATATTAATG 720
AGAP012538-RA      GCGGTAGATGGACTTCCGATGCATAATAGTGGACCAACGCAGCTATGGCCTATATTGATG 718
AGAP011660-RA      GCGGTAGATGGACTTCCGATGCATAATAGTGGACCAACGCAGCTATGGCCTATATTAATG 718
AGAP012490-RA      GCGGTAGATGGACTTCCGATGCATAATAGTGGACCAACGCAGCTATGGCCTATATTAATG 718
                   ******************************************************** ***

AGAP012506-RA      CACATAGTTAATCTACCAGCACTGCCAATCATG 753
AGAP012538-RA      CACATAGTTAATCTACCAGCACTGCCAATCATG 751
AGAP011660-RA      CACATAGTTAATCTACCAGCACTGCCAATCATG 751
AGAP012490-RA      CACATAGTTAATCTACCAGCACTGCCAATCATG 751
                   *********************************
```

Ping-pong network 4:
Sequence Logo 3.5.0 (toolshed.g2.bx.psu.edu/repos/devteam/weblogo3/rgweblogo3/3.5.0)

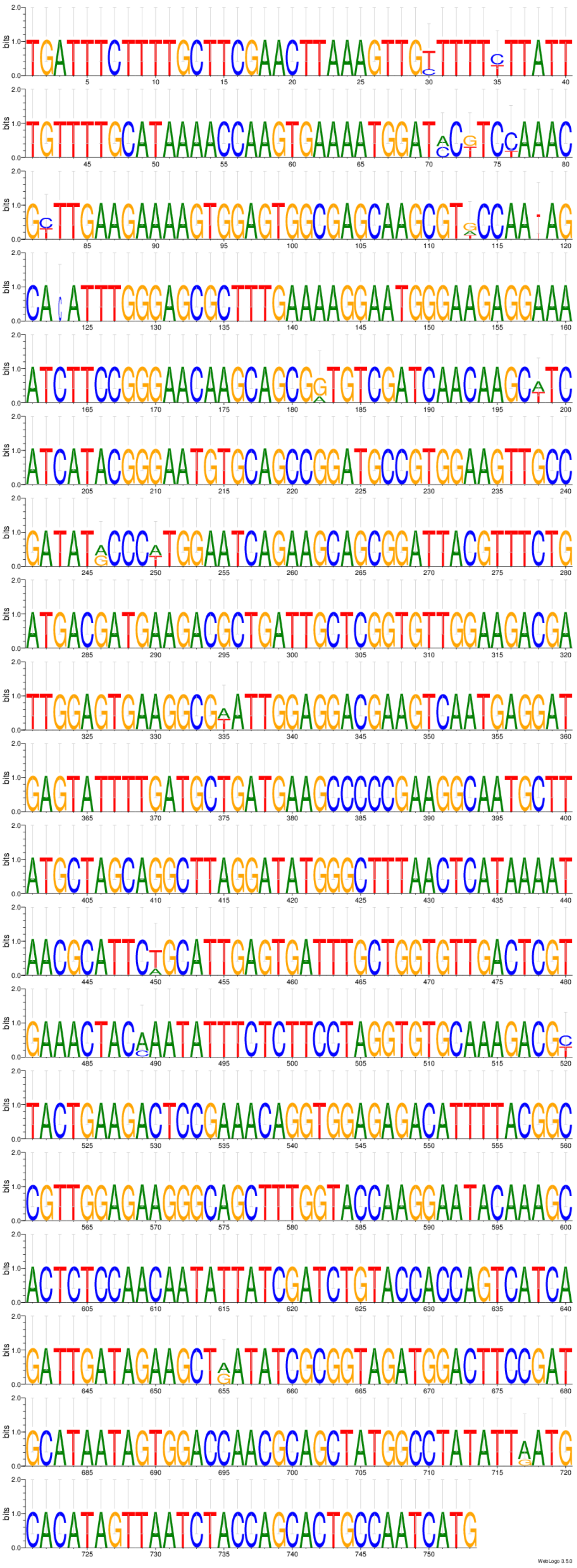

Ping-pong network 5:
CLUSTAL O(1.2.4) multiple sequence alignment (https://www.ebi.ac.uk/Tools/msa/clustalo/)

```
AGAP013064-RA      ------------------------------------------------------------  0
AGAP001078-RA      CGATCTGAATGATGCGAAACAACAAGAAACAAATTAAGTTAACATTTTCCGTAAATTTAA  60
AGAP001079-RA      CGATCTGAATGATGCGAAACACCAAGAAACAAATTAAGTTAACATTTTCCGTAAATTTAA  60
AGAP004894-RA      CGATCTGAATGATGCGAAACAACAAGAAGGAAATTAAGTTAACATTTTCCGTAAATTTAA  60
AGAP012776-RA      ----CTGAATGATGCGAAACAACAAGAAGGAAATTAAGTTAACATTTTCCGTAAATTTAA  56

AGAP013064-RA      ------------------------------------------------------------  0
AGAP001078-RA      GATAAAATCATTGTTCAGTGATTTGTGATCACATAAATTATTCATCGCGCAATATCTGTA  120
AGAP001079-RA      GATAAAATCATTGTTCAGTGATTTGTGATCACATAAATTATTCATCGCGCAATATCTGTA  120
AGAP004894-RA      GATAAAATCATTGTTCAGTGATTTGTGATTACATAAATTATTCATCGCGCAATATCTGTA  120
AGAP012776-RA      GATAAAATCATTGTTCAGTGATTTGTGATCACATAAATTATTCATCGCGCAATATCTGTA  116

AGAP013064-RA      ------------------------------------------------------------  0
AGAP001078-RA      CTATTACAATCTAATGTTCGTGGCTCATTTTATCATCTGGAAAGTGGTGCCTCCTAGACT  180
AGAP001079-RA      CTATTACAATCTAATGTTCGTGGCTCATTTTATCATCTGGAAAGTGGTGCCTCCTAGACT  180
AGAP004894-RA      CTATTACAATCTAATGTTCGTGGCTCATTTTATCATCTGGAAAGTGGTACCTCCTAGACT  180
AGAP012776-RA      CTATTACAATCTAATGTTCGTGGCTCATTTTATCATCTGGAAAGTGGTGCCTCCTAGACT  176

AGAP013064-RA      ------------------------------------------------------------  0
AGAP001078-RA      AGACCGTATATATCAAACCCAATTCGATGAAATTGAATAAACGCGCCAACAGTGAGTTTG  240
AGAP001079-RA      AGACCGTATATATCAAACCCAATTCGATGAAATTGAATAAACGCGCCAACAGTGAGTTTG  240
AGAP004894-RA      AGACCGTATATATCAAACCCAATTCGATGAAATTGAATAAACGCGCCAACAGTGAGTTTG  240
AGAP012776-RA      AGACCGTATATATCGAACCCAATTCGATGAAATTGAATAA--------------------  216

AGAP013064-RA      ------------------------------------------------------------  0
AGAP001078-RA      GTCACCCGTCCTAATTATATGTAGCCATTTGATTGTGGTTTCCGTGTCGGATGGTGGCAT  300
AGAP001079-RA      GTCACCCGTCCTAATTATATGTAGCCATTTGATTGTGGTTTCCGTGTCGGATGGTGGCAT  300
AGAP004894-RA      GTCACCCGTCCTAATTATATGTAGCCATTTGGTTGTGGTTTCCGTGTCGGATGGTGGTAT  300
AGAP012776-RA      ------------------------------------------------------------  216

AGAP013064-RA      ATGATTCTCGTACACACGGCGGAAGCAAAATCACTTGTTATGATTCTATTATTTGGCGCT  60
AGAP001078-RA      ATGATTCTTGTACACATGGCGGAAGCAAATTCACTTGTTATGATTCTATTATTTGGCGCT  360
AGAP001079-RA      ATGATTCTTGTACACATGGCGGAAGCAAATTCACTTGTTATGATTCTATTATTTGGCGCT  360
AGAP004894-RA      ATGATTCTTGTACACATGGCGGAAGCAAAATCACTTGTTATGATTCTATTATTTGGCGCT  360
AGAP012776-RA      ------------------------------------------------------------  216

AGAP013064-RA      TCCAGATTTCATCCATCCAGATCCAGAGAAACATTACAGAACCAACGTGATTTAGTTTTT  120
AGAP001078-RA      TCCAGATTCCATCCATCCAGATCCAGAAAAACATTACAGAACCAACGTGATTTAGTTTTT  420
AGAP001079-RA      TCCAGATTCCATCCATCCAGATCCAGAAAAACATTACAGAACCAACGTGATTTAGTTTTT  420
AGAP004894-RA      TCCAGATTCCATCCATCCAGATCCAGAAAAACATTACAGAACCAACGTGATTTAGTTTTT  420
AGAP012776-RA      ------------------------------------------------------------  216

AGAP013064-RA      GAGCTAGCGTTATATACAATTCCGGAAGTGATGTTTTCGACGTACCGTCCGAACTACTCA  180
AGAP001078-RA      GAGCTAGCGTTATATACAATTCCGGAAGTGATGTTTTCGACGTACGGGTCCGAACTACTC  480
AGAP001079-RA      GAGCTAGCGTTATATACAATTCCGGAAGTGATGTTTTCGACGTACGGGTCCGAACTACTC  480
AGAP004894-RA      GAGCTAGCGTTATATACAATTCCGGAAGTGATGTTTTCGACGTAG-GGTCCGAACTACTC  479
AGAP012776-RA      ------------------------------------------------------------  216

AGAP013064-RA      ATCCTGATAAAGGGCTGTATAAGCGAATTACACTACTCTTACAAGCTGAACCAAACATAT  240
AGAP001078-RA      AATCCTGATATGGGTTGTATAAGCGAATTACACTACTCTTACAAGCTGAACCAAACATAT  540
AGAP001079-RA      AATCCTGATATGGGTTGTATAAGCGAATTACACTACTCTTACAAGCTGAACCAAACATAT  540
AGAP004894-RA      AATCCTGATATGAGTTGCATAAGCGAATTACACTACTCTTATAAGCTGAACCAAACATAT  539
AGAP012776-RA      ------------------------------------------------------------  216

AGAP013064-RA      TTTATAAACATTATTCGTGCGACTTAA---------------------------------  267
AGAP001078-RA      TTTATAAACATTATTCGTGCGACTTAATGCAAAAATAGAATAGCTACAGTTAGTGAGTGA  600
AGAP001079-RA      TTTATAAACATTATTCGTGCGACTTAATGCAAAAATAGAATAGCTACAGTTAGTGAGTGA  600
AGAP004894-RA      TTTATAAACATTATTCGTGCGACTTAATGCAAAAATAGAACAGCTACAGTTAGTGAGTGA  599
AGAP012776-RA      ------------------------------------------------------------  216

AGAP013064-RA      ------------------------------------------------------------  267
AGAP001078-RA      AACGATATCACTATCAACAGTTTAATCATCTCTTAAAACATTTAAAAGGAAATTCTACAT  660
AGAP001079-RA      AACGATATCACTATCAACAGTTTAATCATCTCTTAAAACATTTAAAAGGAAATTCTACAT  660
AGAP004894-RA      AACGATATCACTATCAACAGTTTAATCATCTCTTAAAACATTTAAAAGGAAATTGTACAT  659
AGAP012776-RA      ------------------------------------------------------------  216
```

```
AGAP013064-RA      -----------------------------------------------------------  267
AGAP001078-RA      CACAGTTCATGTATTCTACGCTTAAGCACATGTACGCATTATTAAAGAACGTATATAGAT  720
AGAP001079-RA      CACAGTTCATGTATTCTACGCTTAAGCACATGTACGCATTATTAAAGAACGTATATAGAT  720
AGAP004894-RA      CACAGTTCATGTATTCTACGCTTAAGCACATGTACGCATTACTAAAGAACGTATATAGAT  719
AGAP012776-RA      -----------------------------------------------------------  216


AGAP013064-RA      -------    267
AGAP001078-RA      TAATTAG    727
AGAP001079-RA      TAATTAG    727
AGAP004894-RA      TAATTAG    726
AGAP012776-RA      -------    216
```

Ping-pong network 5:
Sequence Logo 3.5.0 (toolshed.g2.bx.psu.edu/repos/devteam/weblogo3/rgweblogo3/3.5.0)

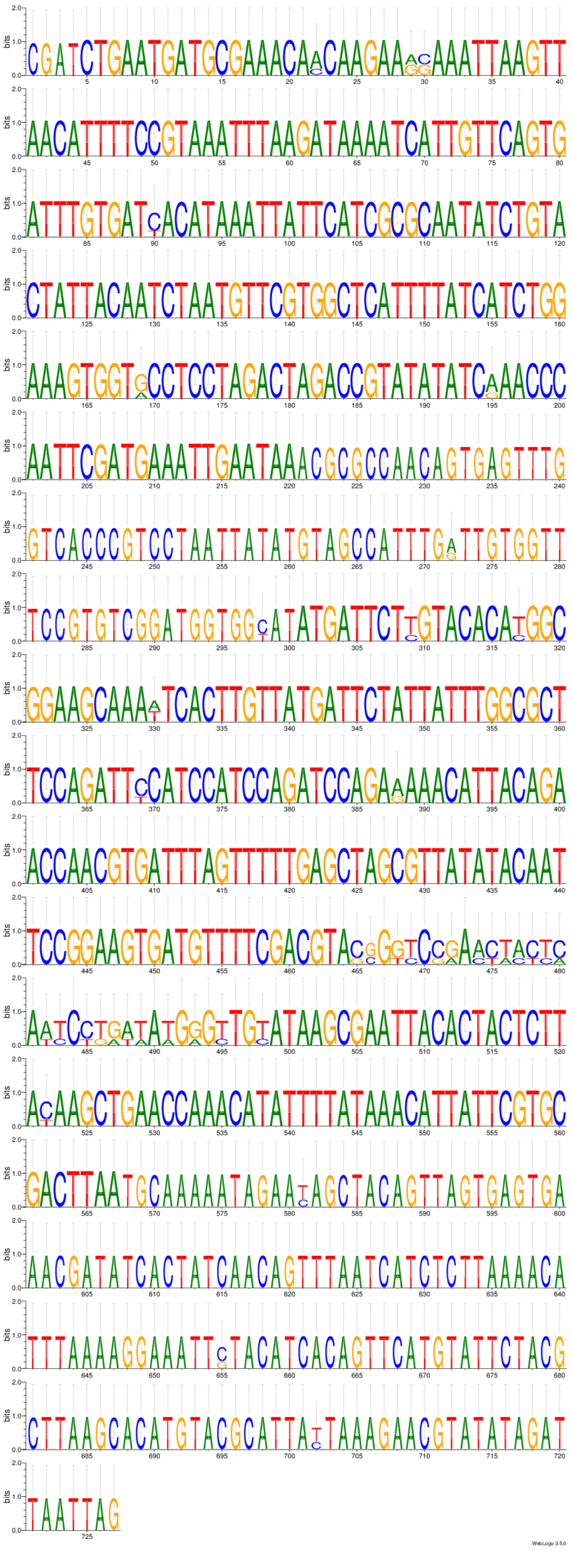

Ping-pong network 6:
CLUSTAL 2.1 multiple sequence alignment

```
AGAP012621-RA      TACCTGCAACAGTAGGGTTACCAAGTAATAATCTTTATTAGCTTTCTAGACTCCATGACG 60
AGAP012622-RA      TACCTGCAACAGTAGGGTTACCAAGTAATAATCTTTATTAGCTTTCTAGACTCCATGACG 60
                   ************************************************************

AGAP012621-RA      GTTTTATGTGTTTGCAGAAAGTGAAGTAATAGAAAATGTTATTTTTTAAAGTATTGAAAC 120
AGAP012622-RA      GCTTTATGTGTTTGCAGAAAGTGAAGTAATAGAAAATGTTATTTTTTAAAGTATTGAAAC 120
                   * **********************************************************

AGAP012621-RA      ATCCTTTAGCGCGCGGATGGTTACGTATTCCGTCTTATGGCCCACAGCCAATTTAGTGC 180
AGAP012622-RA      ATCCTTTAGCGCGCGGATGGTTACATATTCCGTCTTATGGCCCACAGCCAATTTAGTGC 180
                   ************************ ***********************************

AGAP012621-RA      AGCAAAGTTTGGATGGACGAAGCTATGATGCTTTAGACTTTGTTCAGAGTTTGAGGTATG 240
AGAP012622-RA      AGCAAAGTTTGGATGGACGAAGCTATGATGCTTTAGACTTTGTTCAGAGTTTGAGGTATG 240
                   ************************************************************

AGAP012621-RA      GATTATACGGTACATTTTACGTTGCCCCGACGATCTATGGATGGGTAAAGATAACCAGCA 300
AGAP012622-RA      GATTATACGGTACATTTTACGTTGCCCCGACGATCTATGGATGGGTAAAGATAACCAGCA 300
                   ************************************************************

AGAP012621-RA      TCATGTGGCCAAAAATTAATT-ACGTAACTGCCATGATCAAAGCTATTATCGAGCAAGCT 359
AGAP012622-RA      TCATGTGGCCAAAAATTAATTTACGTA-CTGCCATGATCAAAGCTATTATCGAGCAAGCT 359
                   ********************* ***** ********************************

AGAP012621-RA      ACATATGGGCCATTTGCTGGCATTAGTTTTTTATATATTATGTCATTGACCGAAGGAAAG 419
AGAP012622-RA      ACATATGGGCCATTTGCTGGCATTAGTTTTTTATATATTATGTCATTGACCGAAGGAAAG 419
                   ************************************************************

AGAP012621-RA      ACAGCTGTAGAAGCAGTAAAAGAAGTAAAATTGAAATTCCCCACTACATATACACCCATA 479
AGAP012622-RA      ACAGCTGTAGAAGCAGTAAAAGAAGTAAAATTGAAATTCCCCACTACATATACA---GTA 476
                   ******************************************************    **

AGAP012621-RA      GGTCTTGCATTTTGGCCTTTTATTCAAACAATCAATTTTGCTTGTATACCCGAACGGAAT 539
AGAP012622-RA      GGTCTTGCATTTTGGCCTTTTATTCAAACAATCAATTTTGCTTGTATACCCGAACGGAAT 536
                   ************************************************************

AGAP012621-RA      CGTGTGCCATTTGTTGCAACCTGCAGTTTCGTATGGACAGTTTTTCTAGCTTCTATCAAA 599
AGAP012622-RA      CGTGTGCTATTTGTTGCAACCTGCAGTTTCGTATGGACAGTTTTTCTAGCTTCTATCAAA 596
                   ******* ****************************************************

AGAP012621-RA      AATAACTGTATTTCGAATCAAAACTAA--------------------------------- 626
AGAP012622-RA      AATAACTGTATTTCGAATCAAAACTAACATCTGACGAAATAAACAGAACTGCAACGTAGC 656
                   ***************************

AGAP012621-RA      --
AGAP012622-RA      AA 658
```

Ping-pong network 6:
Sequence Logo 3.5.0 (toolshed.g2.bx.psu.edu/repos/devteam/weblogo3/rgweblogo3/3.5.0)

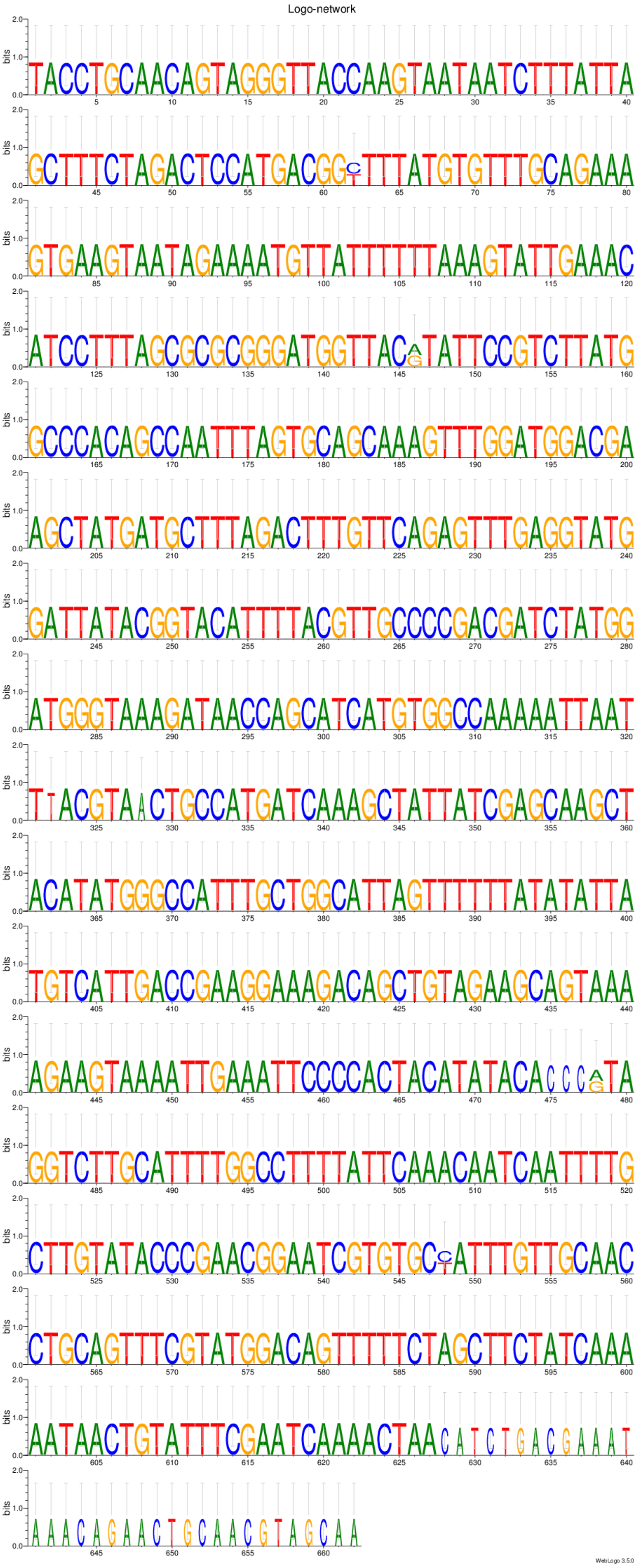

Ping-pong network 7:
CLUSTAL 2.1 multiple sequence alignment

```
AGAP000408-RA      ATGGATAATTTACAGCGTCTGGACAACATCAAGTATTCCGAACAAATTAGAAAGACTGGA 60
AGAP007639-RA      ATGGATAATTTACAGCGTCTGGACAACATCAAGTATTCCGAACAAATTAGAAAGACTGGA 60
AGAP003624-RA      ATGGATAATTTACAGCGTCTGGACAACATCAAGTATTCCGAACAAATTAGAAAGACTGGA 60
                   ************************************************************

AGAP000408-RA      ATGCTCTATAAGCAAAGGGATAAATTGAGGCTAATTGAGAAAGACGCAGCAGATCATGGA 120
AGAP007639-RA      ATGCTCTATAAGCAAAGGGATAAATTGAGGCTAATTGAGAAAGACGCAGCAGATCATGGA 120
AGAP003624-RA      ATGCTCTATAAGCAAAGGGATAAATTGAGGCTAATTGAGAAAGACGCAGCAGATCATGGA 120
                   ************************************************************

AGAP000408-RA      TCAACGTCAGTTGATGACGCTGGATATTCGAATATATCTCATGCGGATTGCAGCATAGCT 180
AGAP007639-RA      TCAACGTCAGTTGATGACGCTGGATATTCGAATATATCTCATGCGGATTGCAGCATAGCT 180
AGAP003624-RA      TCAACGTCAGTTGATGACGCTGGATATTCGAATATATCTCATGCGGATTGCAGCATAGCT 180
                   ************************************************************

AGAP000408-RA      GATCAGAGTGATAATATTGTATCCGATGAATCGACTGTATCCATCGAAGATTTTTTGGAA 240
AGAP007639-RA      GATCAGAGTGATAATATTGTATCCGATGAATCGACTGTATCCATCGAAGATTTTTTGGAA 240
AGAP003624-RA      GATCAGAGTGATAATATTGTATCCGATGAATCGACTGTATCCATCGAAGATTTTTTGGAA 240
                   ************************************************************

AGAP000408-RA      GATGAATCGGAAGAAGAGGTTAGTGGAGACAGTGATGCTGAAGAAGGAGATACCGAACGC 300
AGAP007639-RA      GATGAATCGGAAGAAGAGGTTAGTGGAGACAGTGATGCTGAAGAAGGAGATACCGAACGC 300
AGAP003624-RA      GATGAATCGGAAGAAGAGGTTAGTGGAGACAGTGATGCTGAAGAAGGAGATACCGAACGC 300
                   ************************************************************

AGAP000408-RA      TGTTCATCAAGTTTGCCGTTTGCTGATCGTATAAGAGGATGGGCATTGAAAGCAAATCTT 360
AGAP007639-RA      TGTTCATCAAGTTTGCCGTTTGCTGATCGTATAAGAGGATGGGCATTGAAAGCAAATCTT 360
AGAP003624-RA      TGTTCATCAAGTTTGCCGTTTGCTGATCGTATAAGAGGATGGGCATTGAAAGCAAATCTT 360
                   ************************************************************

AGAP000408-RA      TCACACTACAGCCTCAATCAGCTTCTGCAAATTATTAATACGTCAAAGGTCGATAAACTA 420
AGAP007639-RA      TCACACTACAGCCTCAATCAGCTTCTGCAAATTATTAATACGTCAAAGGTCGATAAACTA 420
AGAP003624-RA      TCACACTACAGCCTCAATCAGCTTCTGCAAATTATTAATACGTCAAAGGTCGATAAACTA 420
                   ************************************************************

AGAP000408-RA      CCCAAGGATGCCAGAACGTTGCTTAAAACAAATAGGGAGCGTGTAAGAGTAGATAAAATA 480
AGAP007639-RA      CCCAAGGATGCCAGAACGTTGCTTAAAACAAATAGGGAGCGTGTAAGAGTAGATAAAATA 480
AGAP003624-RA      CCCAAGGATGCCAGAACGTTGCTTAAAACAAATAGGGAGCGTGTAAGAGTAGATAAAATA 480
                   ************************************************************

AGAP000408-RA      GCAGGAGGAAAATATTGGTATAATGGAATACAACAGTGTTTTTCCAACAGTTTTAAAAAT 540
AGAP007639-RA      GCAGGAGGAAAATATTGGTATAATGGAATACAACAGTGTTTTTCCAACAGTTTTAAAAAT 540
AGAP003624-RA      GCAGGAGGAAAATATTGGTATAATGGAATACAACAGTGTTTTTCCAACAGTTTTAAAAAT 540
                   ************************************************************

AGAP000408-RA      CAAAGCATTCATTTGGACTCGATATTGATAAACATATCAATCGATGGGCTCCCTCTGTAT 600
AGAP007639-RA      CAAAGCATTCATTTGGACTCGATATTGATAAACATATCAATCGATGGGCTCCCTCTGTAT 600
AGAP003624-RA      CAAAGCATTCATTTGGACTCGATATTGATAAACATATCAATCGATGGGCTCCCTCTGTAT 600
                   ************************************************************

AGAP000408-RA      AAGAGTAGCCCTACTCAATTTTGGCCTATATTGATGAATATACATGAATTGCCAGACATC 660
AGAP007639-RA      AAGAGTAGCCCTACTCAATTTTGGCCTATATTGATGAATATACATGAATTGCCAGACATC 660
AGAP003624-RA      AAGAGTAGCCCTACTCAATTTTGGCCTATATTGATGAATATACATGAATTGCCAGACATC 660
                   ************************************************************

AGAP000408-RA      CCAGTGATGATCGTGGCTATTTTTTGTGGTTCTTCAAAGCCAGGCAGTATAGAAGAATTT 720
AGAP007639-RA      CCAGTGATGATCGTGGCTATTTTTTGTGGTTCTTCAAAGCCAGGCAGTATAGAAGAATTT 720
AGAP003624-RA      CCAGTGATGATCGTGGCTATTTTTTGTGGTTCTTCAAAGCCAGGCAGTATAGAAGAATTT 720
                   ************************************************************

AGAP000408-RA      TTAAATCCTTTTGTTGAAGACATTAACAAAGTCCAAGAAGATGGAATAATGATAAATGGA 780
AGAP007639-RA      TTAAATCCTTTTGTTGAAGACATTAACAAAGTCCAAGAAGATGGAATAATGATAAATGGA 780
AGAP003624-RA      TTAAATCCTTTTGTTGAAGACATTAACAAAGTCCAAGAAGATGGAATAATGATAAATGGA 780
                   ************************************************************

AGAP000408-RA      AAAAAAATAAAAGTAAAACTACGAGCAATTATCGCTGATTCTCCGGCTCGTGCGTTTATT 840
AGAP007639-RA      AAAAAAATAAAAGTAAAACTACGAGCAATTATCGCTGATTCTCCGGCTCGTGCGTTTATT 840
AGAP003624-RA      AAAAAAATAAAAGTAAAACTACGAGCAATTATCGCTGATTCTCCGGCTCGTGCGTTTATT 840
                   ************************************************************

AGAP000408-RA      AAAGGTAAGTTACAACATCCTTTCCATGATAAATGCTAA--------------------- 879
AGAP007639-RA      AAAGGTAAGTTACAACATCCTTTCCATGATAAATGCTAACTAACTTTTTGTTTTATTGCA 900
AGAP003624-RA      AAAGGTAAGTTACAACATCCTTTCCATGATAAATGCTAA--------------------- 879
                   ***************************************

AGAP000408-RA      ------------------------------------------------------------
AGAP007639-RA      GGAGTAGCTTATTTCAATGCCAAGCATGGTTGTTTAAAATGTACATGCCATGGAGAATTT 960
AGAP003624-RA      ------------------------------------------------------------
```

```
AGAP000408-RA       -------------------
AGAP007639-RA       AGTGAGCTATCGAAAACCG 979
AGAP003624-RA       -------------------
```

Ping-pong network 7:
Sequence Logo 3.5.0 (toolshed.g2.bx.psu.edu/repos/devteam/weblogo3/rgweblogo3/3.5.0)

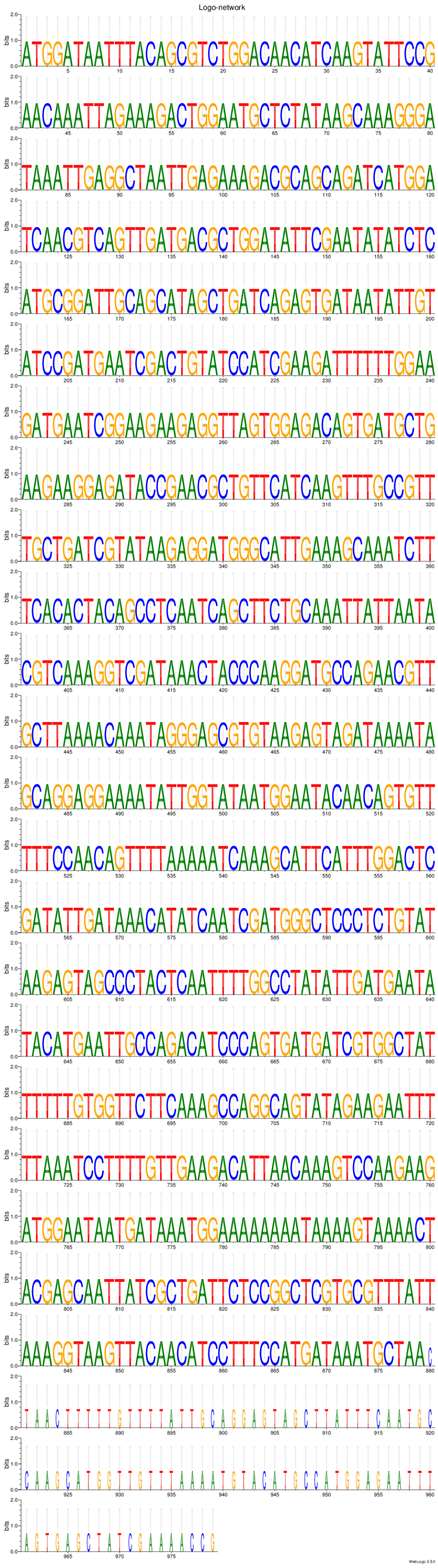

Ping-pong network 8:
CLUSTAL 2.1 multiple sequence alignment

```
AGAP001072-RA      AATGCTAACTCGTTATATCATTTATAGAATATGACTTCTATGTAGGATTGGTTAATGCTG 60
AGAP010287-RA      AATGCTAACTCGTTATATCATTTATAGAATATGACTTCTATGTTGGATTGGTTAATGCTG 60
                   ******************************************* ****************

AGAP001072-RA      AGTTTGATATTGCAGTCCCAGCTTCAGTGGACGAGGGTTCATGCCGCCCAGCAGCAGCAG 120
AGAP010287-RA      AGTTTGGTATTGCAGTCCCAGCTTCAGTGGACGAGGGTTCATGCCGCCCAGCAGCAGCAG 120
                   ****** *****************************************************

AGAP001072-RA      TGTCCAAACTGGGTTATTCCAGATGCTGCCCCTCGTGTGGGAAGTGGCCAGATCCCATGT 180
AGAP010287-RA      TGTCCAAACTGGGTTATTCCAGATGCTGCCCCTCGTGTGGGAAGTGGCCAGATCCCATGT 180
                   ************************************************************

AGAP001072-RA      TGCTTCGCGCATAACAGCTCGTCCTTTATTCGGAGTGGAACGGACACGGAATAATTCAGT 240
AGAP010287-RA      TGCTTTGCGCATAACAGCTCGTCCTTTATTCGGAGTGGAACGGACACGGAATAATTCAGT 240
                   ***** ******************************************************

AGAP001072-RA      GAAAGAGTATCGATGTGATGACACCCTTTTGACGATGTTGGTCAGGGAACCTGTGATGAT 300
AGAP010287-RA      GAAAGAGTATCGATGTGATGACACCCTTTTGACTATGTTGGTCAGGGAACTTGTGATGAT 300
                   ********************************* **************** *********

AGAP001072-RA      GTGGTGCAAAAAAAATGCTACCGGCATCACTTTCGTTCCGCTATTGCACTTGACCCGAGC 360
AGAP010287-RA      GTGGTGCAAAAAAA-TGCTACCGGCATCACTTTCGTTCCGCTATTGCACTTGACCCGAGC 359
                   ************** *********************************************

AGAP001072-RA      TAGGACCGCGACGGCGCTGACGAACTTGCGGCAGCGGGGATGATGCGAATCCCTCCTTCG 420
AGAP010287-RA      TAGGACCGCGACGGCGCTGACGAACTTGCGGCAGCAGGGGTGATGCGAATCCCTCCTTCG 419
                   *********************************** *** ********************

AGAP001072-RA      AAATTGCGGTGCTGAGATGATGATAAAGTAGAAGCTTGTGTGGCATGGTCTCGTCCTTAC 480
AGAP010287-RA      AAATTGCGGTGCTGAGATGATGATAAAGTAGAAGCTTGTGTGGCATGGTCTCGTCCTTAC 479
                   ************************************************************

AGAP001072-RA      TCAAACACTGCTTAAGCTACACCCAAGGTGACATGAATCATCGACCCCCACAGGAATAAG 540
AGAP010287-RA      TAAAACACTGCTTAAGCTACACCCAAGGTGACATGAATCATCGACCCCCACAGGAATAAG 539
                   * **********************************************************

AGAP001072-RA      GCTGCAGCCAGAACGGAATGAACATTTGTTTTCTTTGAACTGGAACGGCAACAGTGGCAG 600
AGAP010287-RA      GCTGCAGCCAGAACGGAATGAACATTTGTTTTCTTTGAACTGGAACGGCAACAGTGGCAG 599
                   ************************************************************

AGAP001072-RA      CATAGACGATCGTCTCGAACACGACGACTGCAACGAACCGACATGGAACCGCGGACAATA 660
AGAP010287-RA      CATAGACGTTCGTCTCGAACACGACGACTGCAACGAACCGACATGGAACCGCGGACAATA 659
                   ******** ***************************************************

AGAP001072-RA      ACTTAAATAACCACTATGCATAAGCACAAATATACTTATGGTTTATTTCCT 711
AGAP010287-RA      ACTTAAATAACCACTATGCATAAGCACAAATATACTTATGGTTTATTTCCT 710
                   ***************************************************
```

Ping-pong network 8:
Sequence Logo 3.5.0 (toolshed.g2.bx.psu.edu/repos/devteam/weblogo3/rgweblogo3/3.5.0)

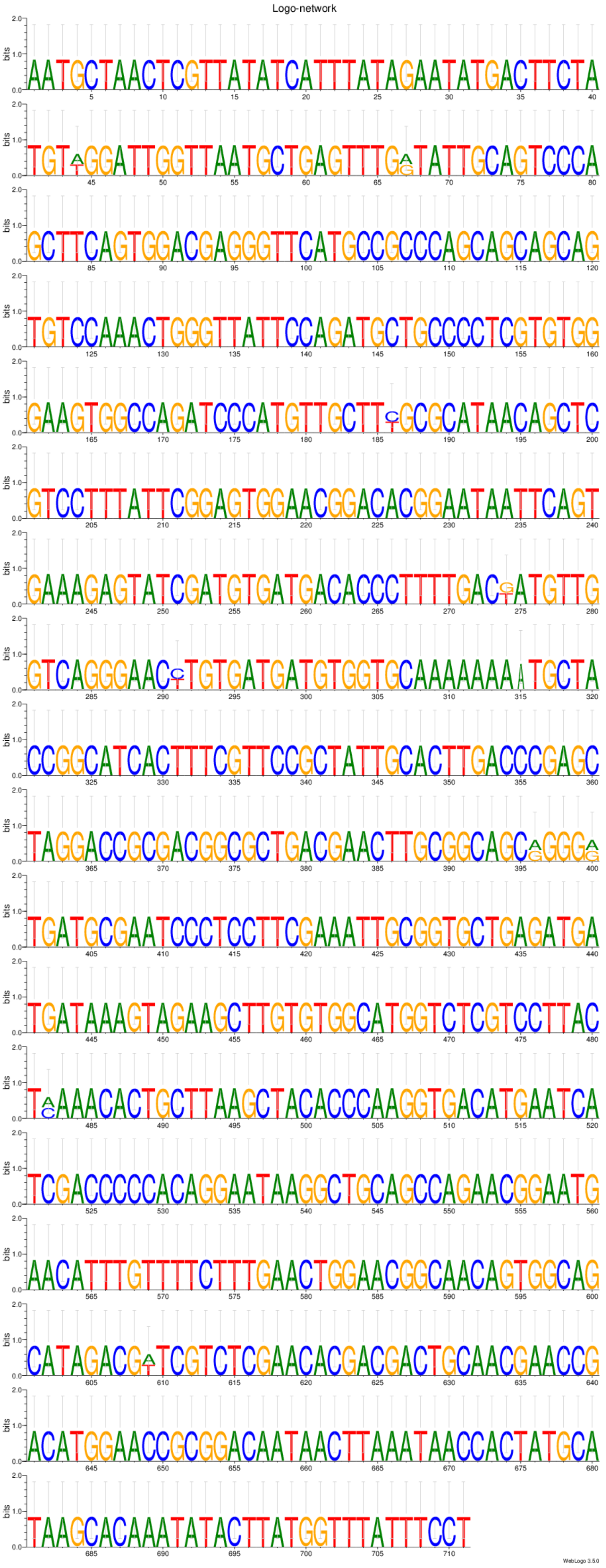

```
Ping-pong network 9:
CLUSTAL 2.1 multiple sequence alignment


AGAP007075-RA      CACTTTCATAGAGCAGGTTCGCTCTTCCCTCACCTCAACCGGGAATCCTTTGAAGAGATA 60
AGAP012821-RA      ------------------------------------------------------------


AGAP007075-RA      GATCGATCCTGCTTGCTGTGATGGATTCCGTGCTCGTAGGGCTGCTGCTATTCACCTGCA 120
AGAP012821-RA      ------------------------------------------------------------


AGAP007075-RA      TCACTTTTGCCCATCCCCAGCAGCAGCATTCCGTTCCGTCCATCTTTATTTACGATGATT 180
AGAP012821-RA      ------------------------------------------------------------


AGAP007075-RA      TTGAGCGGTGTGAAGCGACCGCATCGGTGTACTGCTATGCACGCACCATACTGCGTGTCG 240
AGAP012821-RA      ------------------------------------------------------------


AGAP007075-RA      ATCAATTTCCGGCCGGATTCGTAGTGCCACAAACGGAAAGCACGGATCGAATTGTCCACA 300
AGAP012821-RA      ------------------------------------------------------------


AGAP007075-RA      AGCATCGGGCGCAGTACCTAGAGCTGGGTATCTGTTTGCGGGACTGTGAGCAGGAAGTGA 360
AGAP012821-RA      ------------------------------------------------------------


AGAP007075-RA      AAAGCTTAAGTGCAGCCACCAGGAAGACACTCTTCCAGCCAGAAATTCCTGTGAATTTTA 420
AGAP012821-RA      ------------------------------------------------------------


AGAP007075-RA      CTTTCCTAATTCCGAATGAGCTGTTTCCCACGATGCCGGCCGATAAGCGTCGCTACGAAA 480
AGAP012821-RA      ------------------------------------------------------------


AGAP007075-RA      CGCTGGTGAATGTGTGCGTAAATCAGCGGCTGCGCACGCGTTACAACATTACCGGTTACA 540
AGAP012821-RA      ------------------------------------------------------------


AGAP007075-RA      CGGCCCTCGAGTACTGTCGACCCAAGCCCAAACAAACGGCCCGCCCCTACGATGCATTGG 600
AGAP012821-RA      ------------------------------------------------------------


AGAP007075-RA      AGGTGATCTTTCTTGCCGTCTCGGGGACGCTTGTCAGTACGCTCATCCTCACCACGCTGC 660
AGAP012821-RA      ------------------------------------------------------------


AGAP007075-RA      TGGACGTTTGTCGCGTTAACCAAGAAAATGCAATCGTCTCCGCTTTCTCCGTGCGCCGCA 720
AGAP012821-RA      ------------------------------------------------------------


AGAP007075-RA      ATTGGATGCGCCTGCGGGCCGATGCCGATTCGCCCCTGCACCGCGATCTGTTGTACATCG 780
AGAP012821-RA      ------------------------------------------------------------


AGAP007075-RA      ATGGGCTGCGCGTGCTCGTCAACCATCTGGTGATTGTGCTGCACAGCTTTCTGATCGCGA 840
AGAP012821-RA      ------------------------------------------------------------


AGAP007075-RA      GCGTTGCCCCGGCTCAGAACTACAGCGAGCTGGAGGACCTGGCGAACAATGTGCCGATGC 900
AGAP012821-RA      ------------------------------------------------------------


AGAP007075-RA      GCATCTACCTCTCCTCGAACGCGTACCTGGTGCAGATCTTCTTCACGATCGGTGGCTATC 960
AGAP012821-RA      ------------------------------------------------------------


AGAP007075-RA      TGCTGAGCGTTAACTTTCTGCGCGATGCCGACCGCGGCCCGATCGATGCGCGTTACGCCG 1020
AGAP012821-RA      ------------------------------------------------------------


AGAP007075-RA      GCAACAAGATTCTCAACCGGCTGGTGCGCCTGGTGCCCGTGTATGCGTTCTTTCTACTGT 1080
AGAP012821-RA      ------------------------------------------------------------


AGAP007075-RA      TCTCCGTTAGCCTTAACGTACGCTTCGATGTGAATGTGAACGGGTTTCGGCTGTTTACGG 1140
AGAP012821-RA      ------------------------------------------------------------


AGAP007075-RA      CAGAGAATGCCATTTGCCGTCAGAATTGGTGGACCAATGTGTTGTTTGTGAATAATTTTC 1200
AGAP012821-RA      ------------------------------------------------------------
```

```
AGAP007075-RA      TGTGGCCAAAGGAGCTTTGTTTGATGCACACCTGGTACTTGGCGGCTGATTTGCAACTGT 1260
AGAP012821-RA      ------------------------------------------------------------

AGAP007075-RA      TCCTAATGGCGATGGGTGTGCTGGTGCTGGTGCACCGGAGGCCAAAGAGTGTAGGGGTAG 1320
AGAP012821-RA      -----ATGGCGATGGGTGTGCTGGTGCTGGTGCACCGAAGGCCCAAGAGTGTTGGGGTAG 55
                        ******************************** ***** ******** *******

AGAP007075-RA      TGTTTTTGGTCGGAGTGGTAGTATCGTTTGCTGTTCCCGGGTATATAACGCACCAGCATA 1380
AGAP012821-RA      TGTTTTTGGTCGGAGTGGTAGTATCGTTTGCTGTTCCCGGGTATATAACGCACCAGCATA 115
                   ************************************************************

AGAP007075-RA      AGTTGCACCCTGTGCTGCCGGGTAAGCTTAGTGAAGCTAAATTCCTGACCATGTACGAGC 1440
AGAP012821-RA      AGTTGCACCCGATGCTGCCGGGTAAGCTGAGTGAAGCCAAGTTCCTGACCATGTACGAGC 175
                   **********  **************** ******** ** *******************

AGAP007075-RA      CATGGATAAGGCGCATTTATCTACCAAGCTATGCGAACACTGGCTGCTATCTGTACGGAG 1500
AGAP012821-RA      CTTGGATACGGCGCATTTATCTACCAAGCTATGCAAACACTGGCTGCTACCTGTACGGAG 235
                   * ****** ************************* ************** **********

AGAP007075-RA      TCATTGCCGGGTATCTGTACCATCGCACAAAGAACTACAAGATGCAGCTTGAACGATTTT 1560
AGAP012821-RA      TCATTGCCGGGTATCTGTACCATCGCACAAAGAACTACAAGTTGCAGCTTGAACGATTTT 295
                   ***************************************** ******************

AGAP007075-RA      GGCTCTATCGATTGATCAACGCATACGTAACACCGGTACTGGTCGCCGTGACGGTGTCTT 1620
AGAP012821-RA      GGCTCTATCGAATGATCAACGCATACGTAACACCGGTACTGGTCGCCGTGACGGTGTCTT 355
                   *********** ************************************************

AGAP007075-RA      CCTTCCTCTGGTACGTGATCGAAGTCCCAAAACCTAACCTCTGGGTATCGCTCTACAGTG 1680
AGAP012821-RA      CCTTCCTCTG------------------------------------------------TG 367
                   **********                                                **

AGAP007075-RA      CGCTTTACAGAAACATAATCGGCATCTTTGTGGCTGTGTGCTTTTTGCGCTCCATCGACA 1740
AGAP012821-RA      CACTTTACAGAAACATAATCGGCATCTTTGTGGCTGTGTGCTTTCTGCGCTCCATCGACA 427
                   * ****************************************** ***************

AGAP007075-RA      AACCTCCTGGCATTTTGCGTAGCATTCTCAGCTCCAAACTGCTGACCACACTCGGCAAGC 1800
AGAP012821-RA      ATCCTCCTGGCATTTTGCGTAGCATTCTCAGCTCCAAACTGCTGACCACACTCGGCAAGC 487
                   * **********************************************************

AGAP007075-RA      TTACCTACAGTGCGTACGTACTGCACGATGTGGTGATGCGGTTTTTGCTGTTGCGCGAAA 1860
AGAP012821-RA      TTACCTACAGTGCGTACGTACTACATGATGTGGTGATGCGGTTTTTGCTGTTGCGCGAAA 547
                   ********************** ** **********************************

AGAP007075-RA      ACTTCAACAGTGTGATCAACGTGCAAAAGTTTATAGCTTGGGTGTACATTGTGACGGGGG 1920
AGAP012821-RA      ACTTCAACAGTGTGATCAACGTGCAAAAGTTTATAGCTTGGGTGTACATTGTGACGGGGG 607
                   ************************************************************

AGAP007075-RA      TAGCCTTTGCCGGTGGGCTGGTCGTGTTTCTTGCCATTGAGCAGCCCATGATTCAGCTGA 1980
AGAP012821-RA      TAGCCTTTGCCGGTGGGTTGGTCGTGTTTCTTGCCATTGAGCAGCCCATGATACAGCTGA 667
                   ***************** ********************************** *******

AGAP007075-RA      TTAAACCGTACATAAGCCGAATGTGTCCTGTAGGGGTAAAAGCAAAGCAGAAGTAA 2036
AGAP012821-RA      TTAAACCGTACATAAGCCGAGTGTGCCCTGTAAGGGTAAAAGCAAAGCAGAAGTAA 723
                   ******************** **** ****** ***********************
```

Ping-pong network 9:
Sequence Logo 3.5.0 (toolshed.g2.bx.psu.edu/repos/devteam/weblogo3/rgweblogo3/3.5.0)

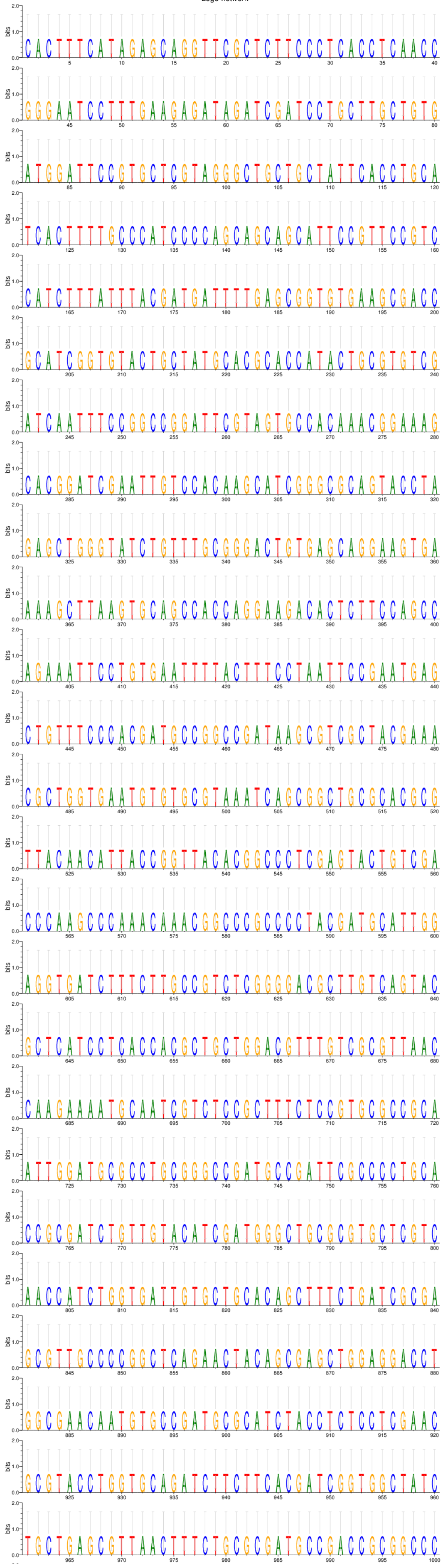

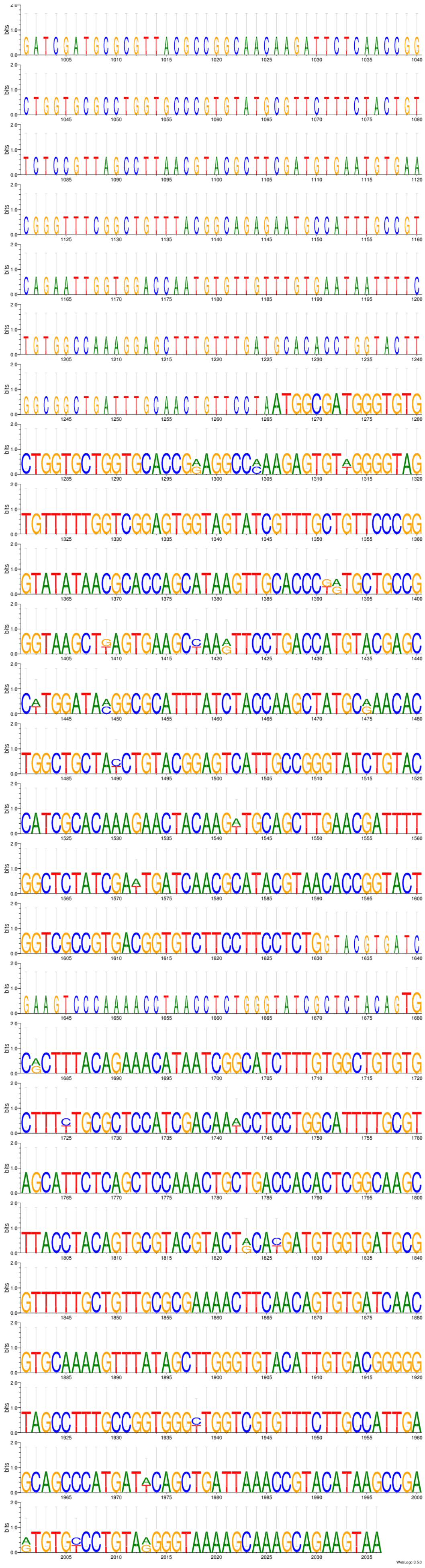
WebLogo 3.5.0

Ping-pong network 10:
CLUSTAL 2.1 multiple sequence alignment

```
AGAP012456-RA      ATGGTGCAGCACGAGCGCAACGTGATACATCGTGACATCCCGGAAAACTTGCAGCTGGGG 60
AGAP012876-RA      ---------CACGAGCGCAACGTGATACATCGTGACATCCCGGAAAACTTGCAGCTGGGG 51
                            ***************************************************

AGAP012456-RA      CACGGTGGTGATCTGAAGATAGCTGATTTCGGTTGGTCCGTGTACGAGCCGACCTTGTTC 120
AGAP012876-RA      CACGGTGGTGATCTGAAGATAGTCGATTTCGGTTGGTCCGTGTATGAGCCGACCCTGTTC 111
                   **********************  ******************** ********* *****

AGAP012456-RA      CGGACGCGGTGCGTTTCGCTCGACTATTTATCGCCCGAGATGGTACATGGTCAGCCGCAC 180
AGAP012876-RA      CGGACGCCGTGCGTTTCGCTCGACTATTTGTCGCCCGAGATAGTGCATGGTCAGCCGCAC 171
                   ******* ********************* *********** ** ***************

AGAP012456-RA      ACAAAAACTGTCGATCTATGGAATTTGGGCGTGCTGGCGTACAAGCTGCTGTGCGGTAAG 240
AGAP012876-RA      ATAAAAACTGTCGATCTATCGAATTTGGGCGTGCTGGCGTACAAGCTGCTGCGCGGTAAG 231
                   * ***************** ******************************* ********

AGAP012456-RA      GCCCCGTTTTTGGCGACCACGTATGAAGAATCGTACCGTAAAATTATGAAGCTGCAGTTT 300
AGAP012876-RA      GTCCCGGTTTTGGCGACCACGTATGAGGAAACGTACTATAAAATCATGAAGCTGCAGTTT 291
                   * **** ******************* *** *****  ****** ***************

AGAP012456-RA      AAGATGCCGCCAGATGTAACGAAGCCGGCGGTCCATCTGATCTCGCGACTGTTCGTTAAG 360
AGAP012876-RA      AAGATGCCGCCAGATATGACGAAGCCGGCGGCCCATCTGATCTCGCGACTGTTCGTTAAG 351
                   *************** * ************* ****************************

AGAP012456-RA      GATCTGGCCAGCCGTATGCCGCTGAAACATGTT--------------------------- 393
AGAP012876-RA      GATCTGGCCAGCCGTATGCCGCTGAAACATGTTGCGTCCATCCCTGGATTCTGGTGCACG 411
                   *********************************

AGAP012456-RA      ------------------
AGAP012876-RA      TGCACAAAAAGTAAATAG 429
```

Ping-pong network 10:
Sequence Logo 3.5.0 (toolshed.g2.bx.psu.edu/repos/devteam/weblogo3/rgweblogo3/3.5.0)

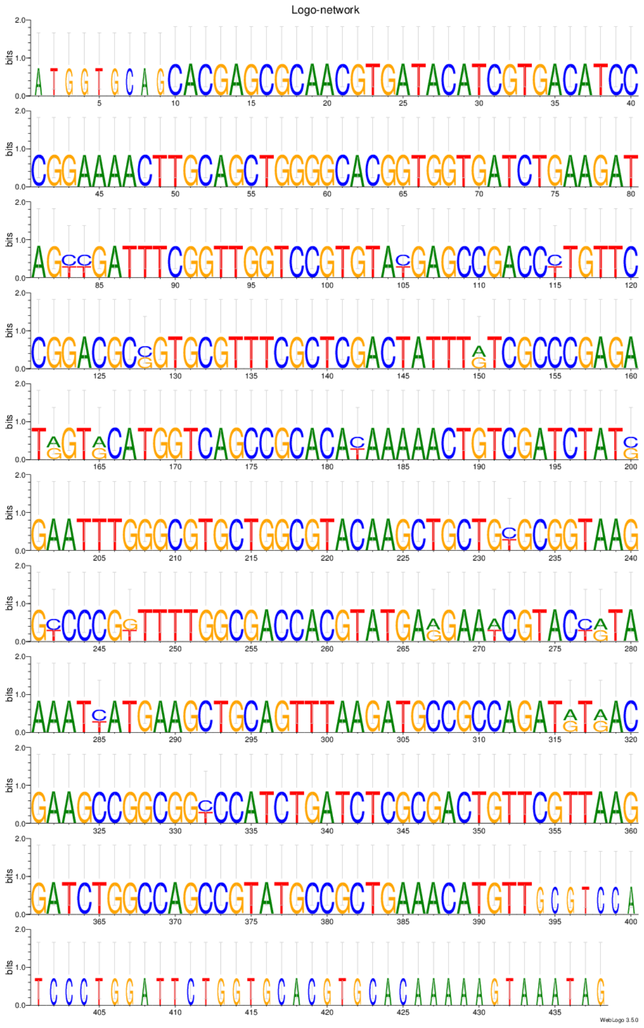

Ping-pong network 11:
CLUSTAL 2.1 multiple sequence alignment

```
AGAP007493-RA      AAAAAAAACAGGTGCTAAATACTCATGTGGCGTGTCCTACTTCCGCACGATCAGTCGCTT 60
AGAP013547-RA      ------------------------------------------------------------


AGAP007493-RA      GCAACGGAGGGGCGATGGCTTCGTGTTAGCAAATCGGTGGAATTTCTTTAGCTAATCGAG 120
AGAP013547-RA      ------------------------------------------------------------


AGAP007493-RA      AGCATTCGCCGTGATTAACTGCAACCAACCACCCAGTGAAGAGAGTTCGCGGTCATCATC 180
AGAP013547-RA      ---------------------------ATGGCGGTTTGTTTACATTTCGTTGCGCTTATC 33
                                              *   *    **   * * ****  *   * ***

AGAP007493-RA      ATCATGTTCCCGTGGTTCGATATTCGGTCGATCAATTTGATCAACTTTCTGTTCGCCAAC 240
AGAP013547-RA      ATCAGGTTTCCGTGGTTCGATATTCGGTCG------------AGCTTTCTTTTCGTCAAC 81
                   **** *** *********************            * ****** **** ****

AGAP007493-RA      CTCATCGTGACGGTCGTGGTGCTCGGTGGACTGTTGGCAACGATCGAAAAGCATCTGCCC 300
AGAP013547-RA      CTCATCGTGACGATCGTGGTGCTCGGTGGACTGTTGGCAACGATCGAAAAGCATCTCCCC 141
                   ************ ******************************************* ***

AGAP007493-RA      ACCGCCATCCGGCAGACGTTCCGGTATGGCAAGCATGCGCTGAAGGGGTCGCCGGACCGA 360
AGAP013547-RA      TCTGCCATCCGGCAGACGTTCCGG------------------------------------ 165
                    * *********************

AGAP007493-RA      TTGGTATCCCTGCTGGAGGTCCCGAAGGCGTGGTTCAAACATTTTTACGTCTTTGCTGCC 420
AGAP013547-RA      ------------------------------------------------------------


AGAP007493-RA      CTCTGGTCGGTGGCCGGATTTGCCGTCATGATGGAGACCTACCTCACTGGCCAGCCAGCG 480
AGAP013547-RA      ------------------------------------------------------CCAGCG 171
                                                                         ******

AGAP007493-RA      CGGGACTACGTAATCGCCTTCCTCGACACGATGGCAACCAACAAGCGCATGGTGCGCACC 540
AGAP013547-RA      CGGGATTACGTAATCGCCTTCCTCGATACGTTGGCAACCAACAAGCGTATGGTGCGCACC 231
                   ***** ******************** *** **************** ************

AGAP007493-RA      ACGCCTACCGAGACGATGGTTGCCATGACGCTGATCACGCTGCAGTGCTTGCGCCGGTTC 600
AGAP013547-RA      ACGCCTACCGAGACGATGGTTGCCATGGCGTTGATCACGCTGCAGTGTTTGCGCCGGTTC 291
                   *************************** ** **************** ************

AGAP007493-RA      TACGAGACCTGGTTCGTGCAGGTGTTCTCGAGCAAGCTGAAAATCAACCTGTCCGCGTAC 660
AGAP013547-RA      TACGAGACCTGGTTCGTGCAGGTGTTCTCGAGCAAGCTGAAAATCA-CCCACCAGTG--- 347
                   ********************************************** **   * * *

AGAP007493-RA      CTCGTCGGGTACATCCATTACTTCGGTACGATCGTGGCGATCCTAGCGCAGGCGGAAGGG 720
AGAP013547-RA      --------GTACGTGGAAAACTT--------TCACAACTATCCTAA-GCAAAGGAAAGCA 390
                           **** *  *  ****        **     * ******  ***   * ***

AGAP007493-RA      TTCACCCGCGCTGGTCCCGTCTCGCTGCCCACCAACGGATATCGGTTCGAGCCCAGCGTC 780
AGAP013547-RA      CTCGTCC-TGTTCGTTCTGTGA-------------------------------------- 411
                    **  **  * * ** * **

AGAP007493-RA      CGATTAGCCCTGTGCGTTGGCGTGTTTTGCTACGCCTGGTACCACCAGTACCTGTCGAAC 840
AGAP013547-RA      ------------------------------------------------------------


AGAP007493-RA      GTGATCCTGGCCAACCTGCGCAAGGACAAGGCGGGCAAAGTGGTGAGCCAGAAGCACAGC 900
AGAP013547-RA      ------------------------------------------------------------


AGAP007493-RA      CTGCCAACCGGCGACTACTTCGATGCTGTATCCTCGCCCCACATGTTCTTCGAGATCGTG 960
AGAP013547-RA      ------------------------------------------------------------


AGAP007493-RA      ATGTACGTCGTGCTGTTCTGTGTGCTGCACCGGAACAGTACGATGGTGTACGTGCTGCTG 1020
AGAP013547-RA      ------------------------------------------------------------


AGAP007493-RA      TGGGTCCTCTCGAATCAGCTGATGAACTCGTGGCTCACCCACCAGTGGTACGTGGAAAAC 1080
AGAP013547-RA      ------------------------------------------------------------


AGAP007493-RA      TTTCCCAACTATCCCAAGCAAAGGAAAGCGCTCGTACCGTTCGTTCTGTAAGACCGACGG 1140
AGAP013547-RA      ------------------------------------------------------------


AGAP007493-RA      TGAGGGTGGGACACAATATCATTCCGGGTGTTGGGTATTTTTCGATTTAGGAAAAACGGG 1200
AGAP013547-RA      ------------------------------------------------------------
```

```
AGAP007493-RA      TACAATCTGAATCGTTTCTGTTTGGTGCTAACCTTAATTGTAAGATCTTTACTGGTACTT 1260
AGAP013547-RA      ------------------------------------------------------------


AGAP007493-RA      TCGCACTTTCATAAAGTGAAATAAAACTGTTAATATGTCTCCTCT 1305
AGAP013547-RA      ---------------------------------------------
```

Ping-pong network 11:
Sequence Logo 3.5.0 (toolshed.g2.bx.psu.edu/repos/devteam/weblogo3/rgweblogo3/3.5.0)

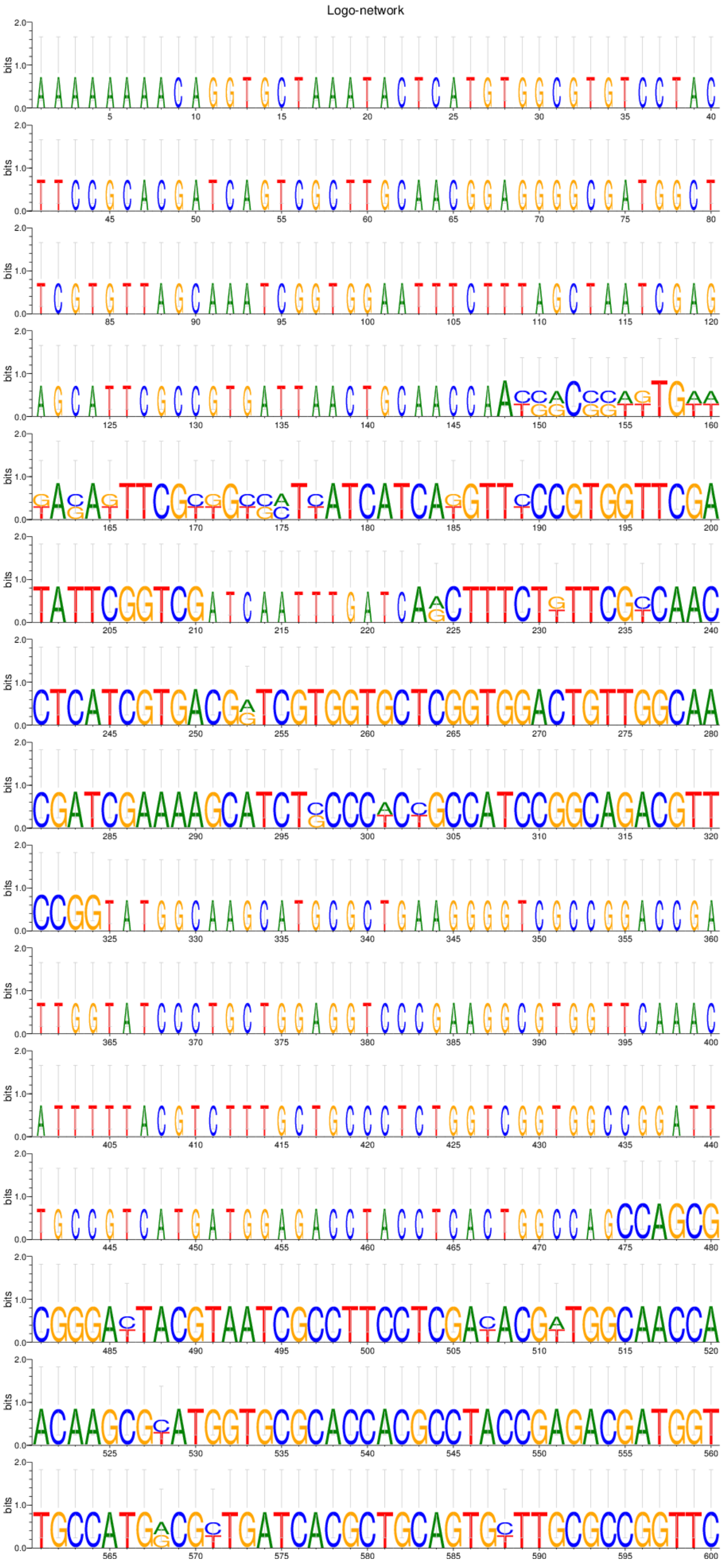

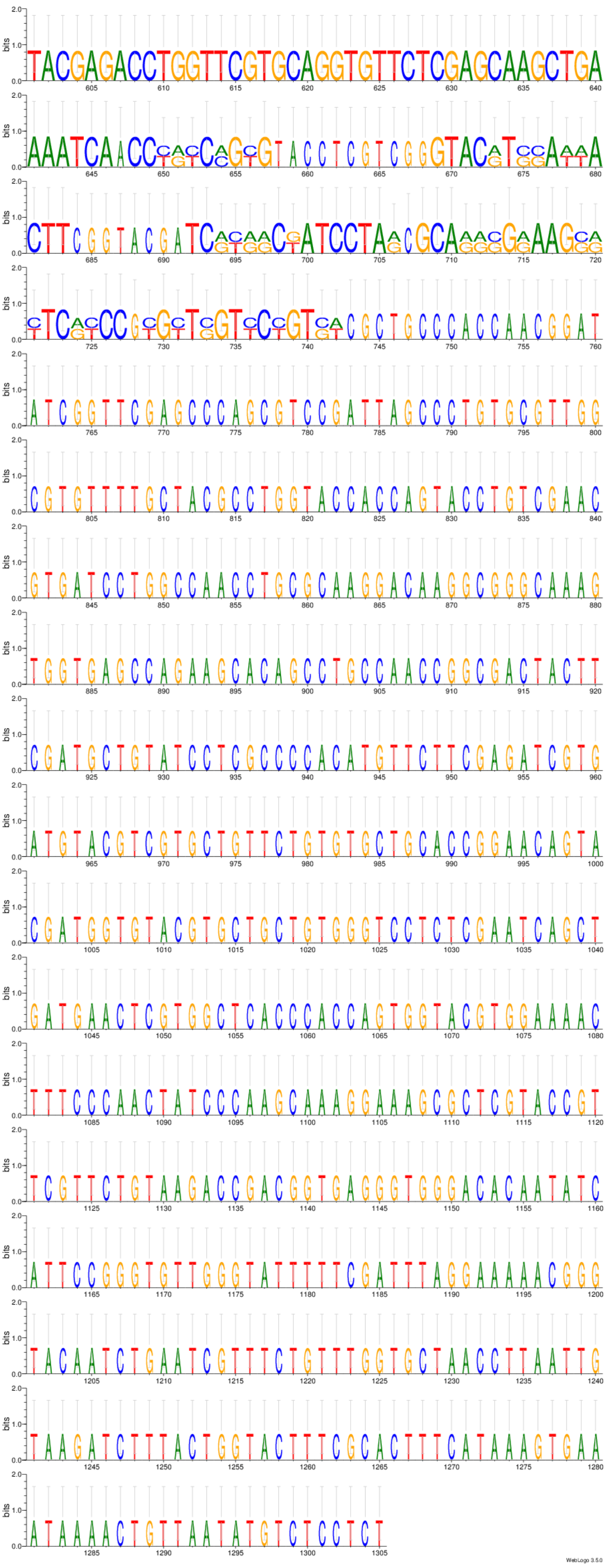
WebLogo 3.5.0

Ping-pong network 12, repeat A:
CLUSTAL 2.1 multiple sequence alignment of AGAP013040-RA and AGAP005338-RA

```
AGAP013040-RA      ATGACCAAGCGCGGTTTTGATTTGATTTTTGGTGGCTGCGAGCGGCATAACACCGTGGAA 60
AGAP005338-RA      ATGACCAAGCGCGGATTTGATTG------TGGTGGCTGCGAGCGGCATAACACCGTGGAA 54
                   ************** *******       *******************************

AGAP013040-RA      GATATGGTGCCTTGCGAAAAGTGCAGAAAGTGGTACCATTACGGGTGTGTCGGCGTCACT 120
AGAP005338-RA      GATATGGTGCCTTGCGAAAAGTGCAGAAAGTGGTACCATTACGGGTGTGTCGGCGTCACT 114
                   ************************************************************

AGAP013040-RA      GTGATGTCAAAATCTTGGTGCAATTTGTGTGTGATGAAGGTGGCAAAGTGCGAAGCTATG 180
AGAP005338-RA      GTGATGTTAAAATCTTGGTGCAATTTGTGTGTGATGAAGGTGGCAAAGTGCGAAGCTAAG 174
                   ******* ************************************************** *

AGAP013040-RA      GAGATGGGCCGTCTAAAGTCCTCTAAGGCAGCCATTGCGATGACTGAGGTGCCTGCTGAA 240
AGAP005338-RA      GAGATGGGCCGTCTAAAGTCCTCTAAGGAAGCCATTGCGATGACTGGGGTGCCTGCTGAA 234
                   **************************** ***************** *************

AGAP013040-RA      AGCTGCGAAGGCGACAAAAGAGCCAAAAGAGCAGCACATAGCGATGGCGAAGTCTCGGCG 300
AGAP005338-RA      AGCTGCGAAGGCGACAAAAGAGCAAAAAGAGCAGCACATAGCGAGGGCGAAGCCTCGGAG 294
                   *********************** ******************** ******* ***** *

AGAP013040-RA      AATAAGCTGGCTACAGAGGAAAAAGAGCCCACTACATTCCATGCTGCAAAAAGGCCTCCT 360
AGAP005338-RA      AATAAGCTGGCTACAGAGGAAAAAGAGCCCACTACATTCCATGCTGCAAAAAGGCCTCCT 354
                   ************************************************************

AGAP013040-RA      ACCACTAATTTACCGACAACAACGCAACTCAAAAGGTCTAGGCAAGCGGCACTCGAAAAG 420
AGAP005338-RA      ACCGCAAATTCATCCACAACAGCGCAACACAAAAGGTCTAGGCAAGCGGCACTCGAAAAG 414
                   *** * **** * * ****** ****** *******************************

AGAP013040-RA      CTTATGGAAGTGCAACAGCAAGAACGGGAGATGGCCATAAAAGAGTACGAGCTCAAAATG 480
AGAP005338-RA      CTTATGGAAGTGCAACAGCAAGAACGGGAGATGGCCATGAAAGAGTACGAGCTCAAAATG 474
                   ************************************** *********************

AGAP013040-RA      GCTAACCTCAAGCTCGAACATTTAAAGATTCGACTTCAGCTCGAGGAGGAAAAGGTTTCG 540
AGAP005338-RA      GCTAACCTCAAGCTCGAACATTTAAAGATTCGACTTCGGCTCGAGGAGGAAGAGGCATCG 534
                   ************************************* ************* ***  ***

AGAP013040-RA      ATTCGATCTGTGACCGATGCACGGATCAAGCATCAGCAGCAGCAGCATCAGCAGCAGCAT 600
AGAP005338-RA      ATTCGATCTGTGACCGATGCACGGATCAAGCATCAGCAGCAGC---ATCAGCAGCAGCAT 591
                   *******************************************   **************

AGAP013040-RA      CAGCAGCATCAGCAGCATCAGCAGCATCAGCAGCAGCAGCATCAGCAGCAGCAGCATCAG 660
AGAP005338-RA      CAGCAGCAGCATCAGCATCATCAGCATCAGCAGCAGCACCAGCATCAGCAACAGCACAAG 651
                   ******** ** ******** ***************** ** ** ***** *****  **

AGAP013040-RA      CAGCAGCAGCAT-CAGCAGCAGC-AGCATCAGCTGCAGCAGCAGCATCAGCTGCAGCAGC 718
AGAP005338-RA      CAGAAACAGTGTACGATAGCGGTGAGTGATGGTGAAATTGGTGAGCTCGCTAACGTCGAA 711
                   *** * ***  * *   *** *  **     *    *   *     **     *  *

AGAP013040-RA      ATCATCAGCATCATCAGCATCATCAGCATCAGCAGCATCAGCAGCAGCAGCAGCAGCAGC 778
AGAP005338-RA      ATGGTAAATGTTG-CGACGAAACCG--TTCGACACCCTTGGCGATTGA------------ 756
                   **  * *   *   *  *   * *    **  ** * *  **    *

AGAP013040-RA      AGCAGCAGCAGCATCAGCATCAGCATCAGCATCAGCATCAGCAACAGCACAAGCAGAAAC 838
AGAP005338-RA      ------------------------------------------------------------


AGAP013040-RA      AGTGTACGATAGCAGTGAGTGATGGTGAAATTGGTGAGCCCGCTAACGTCGAAATGGTAA 898
AGAP005338-RA      ------------------------------------------------------------


AGAP013040-RA      ATGTGAATCCTGCGACAGTGGCAGTCGTAGACATGTCAGTTTTTGCCAAGTATCCTGAGA 958
AGAP005338-RA      ------------------------------------------------------------


AGAP013040-RA      TGGCGCGGCGATCCACGCAATCACATAACCTTCCGTCGGATCACGTACACTCCCTACTAA 1018
AGAP005338-RA      ------------------------------------------------------------


AGAP013040-RA      ACTGCCATGTGCGACACAACACATTTCTCCATGAGACGGTCCAATCTGGAGGAACAGAAT 1078
AGAP005338-RA      ------------------------------------------------------------


AGAP013040-RA      CATTGACAAAAACTTGTGCGTCGTGA 1104
AGAP005338-RA      --------------------------
```

Ping-pong network 12, repeat A:
Sequence Logo 3.5.0 (toolshed.g2.bx.psu.edu/repos/devteam/weblogo3/rgweblogo3/3.5.0) for AGAP013040-RA and AGAP005338-RA

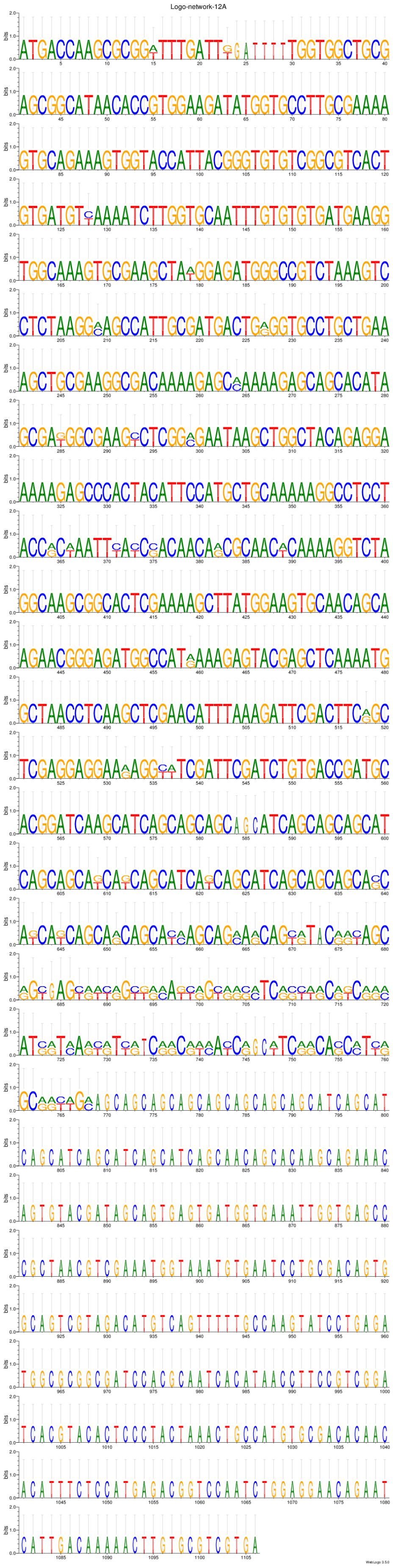

Ping-pong network 12, repeat B:
CLUSTAL 2.1 multiple sequence alignment of AGAP013040-RA positions 938-1104 and AGAP011262-RA positions 446-608

```
AGAP013040-RA_pos938-1104      TTTTTGCCAAGTATCCTGAGATGGCGCGGCGATCCACGCAATCACATAAC 50
AGAP011262-RA_pos446-608       TTTTTGCCAAGTATCCTGAGATGGCGCGACGATCGACGCAATCACATAAC 50
                               **************************** ***** ***************

AGAP013040-RA_pos938-1104      CTTCCGTCGGATCACGTACACTCCCTACTAAACTGCCATGTGCGACACAA 100
AGAP011262-RA_pos446-608       CTTCCGTCAGATCACGTACACTCCCTACTAAACTGCCATGTGCGACACAA 100
                               ******** *****************************************

AGAP013040-RA_pos938-1104      CACATTTCTCCATGAGACGGTCCAATCTGGAGGAACAGAATCATTGACAA 150
AGAP011262-RA_pos446-608       TACATTTCTCCATGAGACGGTCCAATCTGGAGGAATAGAATCGTTGACAA 150
                                ********************************** ****** *******

AGAP013040-RA_pos938-1104      AAACTTGTGCGTCGTGA 167
AGAP011262-RA_pos446-608       AAACTTGTGCGTC---- 163
                               *************
```

Ping-pong network 12, repeat B:
Sequence Logo 3.5.0 (toolshed.g2.bx.psu.edu/repos/devteam/weblogo3/rgweblogo3/3.5.0) for AGAP013040-RA positions 938-1104 and AGAP011262-RA positions 446-608

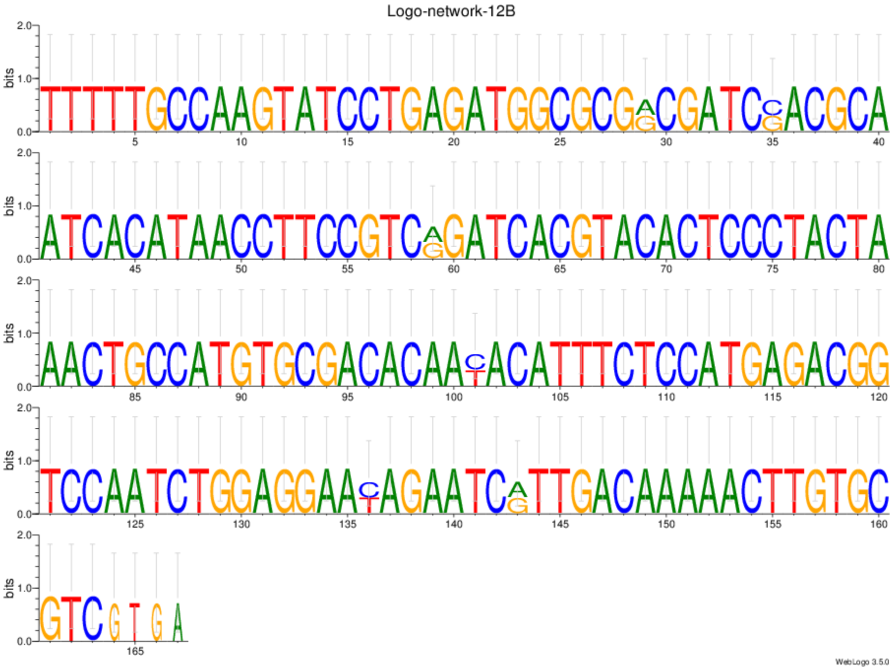

```
Ping-pong network 13:
CLUSTAL 2.1 multiple sequence alignment


AGAP001089-RA      --------------------------GAGTGTGCGAATCATGACTACAATGTTTACAATC 34
AGAP012592-RA      --------------------------GAGTGTGCGAATCATGACTACAATGTTTACAATC 34
AGAP010306-RA      TGTCTTTACACGCATTGTTATTGTGTGAGTGTGCGAATCATGACTACAATGTTTACAATC 60
                                             **********************************

AGAP001089-RA      CAAGAGCTGTACAGTAAATTAAATAAGATTCGGTGCTTTCAAATAAAATTGGATAGAAAA 94
AGAP012592-RA      CAAGAGCTGTACAGTAAACTAAATAAGATACGGTGCTTTCAAATAAAATTGGATCGAAAA 94
AGAP010306-RA      CAAGAGCTGTACAGTAAACTAAATAAGATTTGGTACTTTCAAATAAAATTGGATCGAAAA 120
                   ****************** **********  *** ******************* *****

AGAP001089-RA      TTGAAAAAGTTTTGTAATTTGCATTACATACAGTCCATCACGGTGCCGCATCGTACCAGC 154
AGAP012592-RA      TTGAAAAAGTTTCGTAATTTGCATTACATACAGTGCATCACGGTGCCGCATCGTACCAGC 154
AGAP010306-RA      TTGAAAAAGTTTCGTAATTTGCATTACATACAGTGCATCACGGTGCCGCATCGTACCAGC 180
                   ************ ********************* *************************

AGAP001089-RA      AGGTCGATAAATGTTGTGAATGAAGGAACGGCGG-TATATTAGGAATGGTTGTTTGAGCA 213
AGAP012592-RA      AGGTCGATAAATGTTGTGAATGAAGGAACGGCGG-TATATTAGGAATGGTTGTTTGAGCA 213
AGAP010306-RA      AGGTCGATAAATTTT--------AGGGGCGGTGTCTATTCTACGAACAATTCATATCGCC 232
                   ************ **        ***  *** *  ***  ** ***   **  *   **

AGAP001089-RA      GCACGATGGCAGCGAAGCG-GTGAACCAACGCCACTTTGTTGGGCGGTGGAAGTAAATGT 272
AGAP012592-RA      GCACGATGGCAGCGAAGCG-GTGAGCCAACCCCACTTTGTTGGGCGGTGGCAGTAAATGT 272
AGAP010306-RA      GGTGATTAGCTACGAGGTAAGCGGCATAGGTGTGCTTCTTCATGCCGTATGATTAAAAAT 292
                   *     * **  *** *   * *    *      ***  *   ** **   * ****  *

AGAP001089-RA      AG--TAGCCACTGATA-ACCG----ACAAGAATATTTTGGTGATGATGAAGTGTACCTCT 325
AGAP012592-RA      AG--TAGCCACTGATA-ACCG----ACGAGAATATTTTGGTGTTGATGAAGTGTACCTCT 325
AGAP010306-RA      CCCTTCGCCACTGGGACACCGTGTGACGGAACCGTTCAAAACTAGCTTGAAGCTGCGCTA 352
                       * *******  * ****    **   *   **        * *  *   * *

AGAP001089-RA      GTTTGGAAGGTCTATTC-GACGAA--------CAATTCATATCG----CCGGTGATTAGC 372
AGAP012592-RA      GTTTGGAAGGTCTATTC-GACGAA--------CAATTCATATCG----CCGGTGATTAGC 372
AGAP010306-RA      GCTTGTGAAGTCTCTCCCGTCGAAGGTGTGCGCAAGCTACTTTGTACCCCGGCTGCTGGT 412
                   * ***  * **** * * * ****        ***   *  * *    ****    * *

AGAP001089-RA      TACGAGGTAA--GCGGCATAGGTATGCTTCTTCATGCCGTATGATTAA--ACATCCCTTC 428
AGAP012592-RA      TACGAGGTAA--GTGGCATAGGTGTTCTTGTTCATGCCGTATGATTAA--AAATCCCTTC 428
AGAP010306-RA      GATCTAGGAACGGCAGCTCAGG-ATTTGCAAGCATTTAGTTTACTGGACGGGAGCAAATC 471
                    *    * **  *  **  ***  *       ***   ** *  *  *    * *   **

AGAP001089-RA      GCCACTGGGACACCGTGTGACGGAACCGTTCAAAACTAGCTT------------------ 470
AGAP012592-RA      GCCACTGGGACACCGTGTGACGGAACCGTTTAAAACTAGCTT------------------ 470
AGAP010306-RA      ATTATCTAAGCCAGGAGCTACGGCGGCATACCGAGTGGTCTCGTTTTCGTTGACGGCTGC 531
                      *      *   * *  ****   * *    *     **

AGAP001089-RA      ------------------------------------------------------------
AGAP012592-RA      ------------------------------------------------------------
AGAP010306-RA      TGGAACAAAAATGCCAAAAGATGTACGGTGGAATGCAGAAACGCTATCTGACGTTCGGGC 591


AGAP001089-RA      ------------------------------------------------------------
AGAP012592-RA      ------------------------------------------------------------
AGAP010306-RA      GGGGAAGGAGCAGCAGGAACTTGAAGAAGTTGTCTTGCGCTAGCCAAGAGAGGAGTCAAG 651


AGAP001089-RA      ------------------------------------------------------------
AGAP012592-RA      ------------------------------------------------------------
AGAP010306-RA      TCATAGACACCATTAACAATATATCGTTGCCAAATCGTACACGGACGCAACGCCGCAGCT 711


AGAP001089-RA      ------------------------------------------------------------
AGAP012592-RA      ------------------------------------------------------------
AGAP010306-RA      GACCCTCAGTTTTGCTTTTGGATTTGTTACTCGTGCTGTAAGATGTGTTCCTCGTTTTTA 771


AGAP001089-RA      ------------------------------------------------------------
AGAP012592-RA      ------------------------------------------------------------
AGAP010306-RA      CATATTCTCTTACGAAATCTTATAAACTATACCGGTTTCCATTCGAAGAAAGAGGACTAC 831


AGAP001089-RA      ------------------------------------------------------------
AGAP012592-RA      ------------------------------------------------------------
AGAP010306-RA      ATTTAAATTATGCCAACGATGGATCGTTTTGGACAGATTGACTATTTTGCCGTGAAATCT 891


AGAP001089-RA      ------------------------------------------------------------
AGAP012592-RA      ------------------------------------------------------------
AGAP010306-RA      CACATCGCCTCCTAGTTTGTACCAACATACACACATCGATATCGTTAATCGACATTGTAG 951
```

```
AGAP001089-RA      --------------------------------------------------
AGAP012592-RA      --------------------------------------------------
AGAP010306-RA      GTATTCAGATTATATAAACTGTACGAACTACATGTTCGTCGTTCCTCTA 1000
```

Ping-pong network 13:
Sequence Logo 3.5.0 (toolshed.g2.bx.psu.edu/repos/devteam/weblogo3/rgweblogo3/3.5.0)

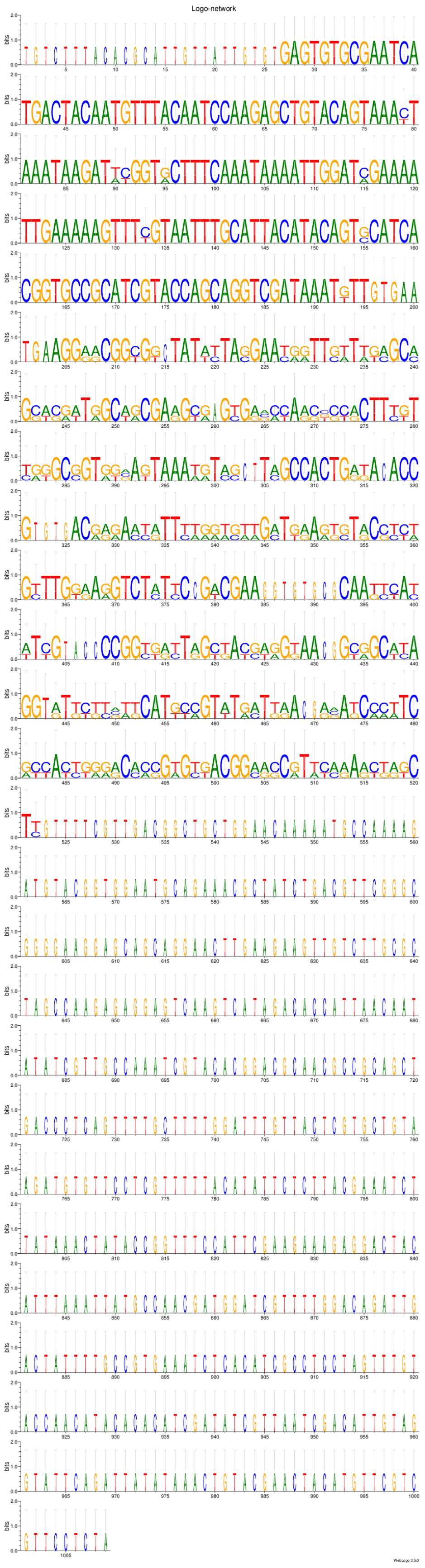

Ping-pong network 14:
CLUSTAL 2.1 multiple sequence alignment

```
AGAP012450-RA      ATGCCATTCTCAATAGTGCAAACCCGTGGGCCTAAAGGGTACGCTGAATCAAGCATTGTT 60
AGAP012715-RA      ATGCCATTCTCAATAGTGCAAACCCGTGGGCCTAAAGGGTACGCTGAATTAAGCATTGTT 60
AGAP012588-RA      CTGCCATTCTCAATAGTGCAAACCCGTGGGCCTAAAGGGTACGCTGAATTAAGCATTGTT 60
                    ************************************************ **********

AGAP012450-RA      CCTGATTCCTGGGTTCAAGGCACTAGCCATAAAAAAA---ACATAT-TTTGGCCAAACGT 116
AGAP012715-RA      CCTGATTCCTGGGTTCAAGGCACTAGCCATAAAAAAACATATCTGT-TTTGGCCAAACGT 119
AGAP012588-RA      CCTGATTCCTGGG-------CACTAGCCATAAAAAAC---ATATCTGTTTGGCCAAACGT 110
                   *************       ****************    *  * * *************

AGAP012450-RA      TAAAGGAAACATACAACATCTAATTACGAATGATAATAGTGTTCCGGAAGAAGGATGGAT 176
AGAP012715-RA      -AAAGGAAACA--CAACATCTAATTACGAATGATAATAGTGTTCCGGAAGAAGGATGGAC 176
AGAP012588-RA      TAAAGGAAACATACAACATCTAATTACGAATGATAATAGTGTTCCGGAAGAAGGATGGAC 170
                    **********  ********************************************** 

AGAP012450-RA      GAAACAGGAGTGTCAGATTAAGCGGCAAGGATTATCATACGAAGCTGCTGATTGCACTAT 236
AGAP012715-RA      GAAACAGGAGTGTCAGATTAAGCGGCAAGGATTATCATACGAAGCTGCTGATTGCACTAT 236
AGAP012588-RA      GAAACAGAAGTGTCAGATTAAGCGGCAAGGATTATCATACGAAGCTGCTGATTGCACTAT 230
                   ******* ****************************************************

AGAP012450-RA      AATGATCATATCCGGCGAATCATCTACAGATTCCGGAAAGGAACCGATATTAGCTAGTTG 296
AGAP012715-RA      AATGATCATGTCCGGTGAATCATCTACAGATTCCGGAAAGGAACCGATATTAGCTAGTTG 296
AGAP012588-RA      AATGATCATGTCCGGCGAATCATCTACAGATGCTGGAAAGCAACCGATATTAGCTAGTTG 290
                   ********* ***** *************** * ****** *******************

AGAP012450-RA      CAGAGAAACAACTTCATACGCTGATATGTTTTCAACAAGAATTGTGGAAAACTCTTTAAC 356
AGAP012715-RA      CAGAGAAACAACTTCATACGCTGATATGTTTTCAACAAGAATTGTGGAAAACTCTTTAAC 356
AGAP012588-RA      CAGAGAAACAACTTCATACGCTGATATGTTTTCAACAAGAATTGTGGAAAACTCTTTAAC 350
                   ************************************************************

AGAP012450-RA      TCCAATCCGCGTCGAGAATGAAATCGACATCTCCTCACCACCTACGTCGCCTCTTACACC 416
AGAP012715-RA      TCCAATCCGCGTCGAGAATGAAATCGACATCTCCTCACCACCTACGTCGCCTCTTACACC 416
AGAP012588-RA      TCCAATCCGCGTCGAGAATGAAATCGATATCTCCTCACCACCTACGTCGCCTCTTACACC 410
                   *************************** ********************************

AGAP012450-RA      AAGTTTGTTAGAAGTGGATGAATATGTTGGTTCAGTTGAGGTAGTCCCAGAAGAAGTTTC 476
AGAP012715-RA      AAGTTTGTCAGAAGTGGATGAATATGTTGGTTCAGTTGAGGTAGTCCCAGAAGAAGTTTC 476
AGAP012588-RA      AAGTATGTCAGAAGTGGATGAATATGTTGGTTCAG-----GTAGTCCCAGAAGAAGTTT- 464
                   **** *** **************************     ******************* 

AGAP012450-RA      ACGTGACAAGACGATAAAAGAAATAATAGAAGCCCAAACTAATATGATGATAGAGCTTCG 536
AGAP012715-RA      ACGTGACAAGACGATAAAAGAAATACTAGAAGCCCAAACTAATATGATGATAGAGCTTCG 536
AGAP012588-RA      ACGTGACAAGACGATAAAAGAAATACTAGAAGCCCAAACTAATATGTTGATAGAGCTTCG 524
                   ************************ ********************* *************

AGAP012450-RA      TAATGAGCAAAAGCAGATTTTAAAAGAACAAAAGAAAATAGTGAGTCGTATCGGCATAAT 596
AGAP012715-RA      TAATGAGCAAAAGCAGATTTTAAAAGAACAAAAGAAAATAGTGAGTCGTATCGGCATAAT 596
AGAP012588-RA      TAATGAGCAAAAGCAGATTTTAAAAGAACAAAAGAAAATAGTGAGTCGTATCGGCATAAT 584
                   ************************************************************

AGAP012450-RA      CGAAGTACAATTAAATACTTTACTTAATCATACCGAAGTAAGTACTACTAATCTTACTGG 656
AGAP012715-RA      CGAAGTACAATTAAATACTTTACTTAATCATACCGTAGTAAGTACTACTAATCTCACTGG 656
AGAP012588-RA      CGAAGTACAATTAAATACTTTACTTAATCATACCGAAGTAAGTACTTCTAATCTTACTGG 644
                   *********************************** ********** ******* *****

AGAP012450-RA      TTTTGACCATCCATTCAAAGACAATGCCGAAGATCTGGAAAAATTTGAAAAAGATTTAGA 716
AGAP012715-RA      TTTTGACCATCCATTCGTAGACAATGCCGAAGATCTGGAAAAATTTGAAAAAGATTTAGA 716
AGAP012588-RA      TTTTGACCATCCATTCGTAGAGAATGCCGAAGATCTGGAAAAATTTGAAAAAGATTTAGA 704
                   ****************  *** **************************************

AGAP012450-RA      TGAGGAAGAATATTATATCCAAATAGTAAGCTTATTGAAACAAAAAATTAATGATAAAGA 776
AGAP012715-RA      TGAGGAAGAATATTATACCCAAGTAGTAAGCTTATTGAAACAAAAAATTATTGATAAAGA 776
AGAP012588-RA      TGAGGAAGAATATTATACCCAAGTAGTAAGCTTACTGAAACAAAAAAATATTGATAAAGA 764
                   ***************** **** *********** ************ ** *********

AGAP012450-RA      TATAAACAACAGAATGTTCGCAACCCTTGATGCCCTTTTCGATAGGAGTTTCTTAACAAA 836
AGAP012715-RA      TATAAACAACAGAATGTTAGCAACCCTTGATGCCCTTTTCGATAGGAGTTTCTTAACAAA 836
AGAP012588-RA      TATAAACAACAGAATGTTAGCAACCCTTGATGCCCTTTTCGATACGAGTTTTTTAACAAA 824
                   ****************** ************************* ****** ********

AGAP012450-RA      GTGCACATGGACCGGTATTTCTAAATCAGGAACTAAAATAGCAATGCATTCATTTAAAAA 896
AGAP012715-RA      GTGCACATGGACAGGTATTTCTAAATCAGGAACTAAAATAGCAATGCATTCATTTAAAAA 896
AGAP012588-RA      GTGCACATGGACAGGTATTTCTAAATCAGGAACTAAAATAGCAATGCATTCATTTAAAAA 884
                   ************ ***********************************************

AGAP012450-RA      TGTAGTAAATCTTTTTAAATGCGTAGGAAGTACTAACCTTGTGCCAGTCACAGATGAAAC 956
AGAP012715-RA      TGTAGTAAATCTTTTTAAATGCGTAGGAAGTACTAACCTTGTGCCAGTCACAGATGAAAC 956
AGAP012588-RA      CGTAGTAAATCTTTGTAAATGCGTAGGAAGTACTAACCTTGTGCCAGTCACATATGAAAC 944
                    ************* ************************************* *******
```

```
AGAP012450-RA      GGTGCAAGGTTTCTTCATGAATAGATTGAAACACGCATTAGAACGATCTAAGGCCAAAGG 1016
AGAP012715-RA      GGTGCGAGGTTTCTTCATGAATAGATTGAAACACGCATTAGAACGATCTAAGGCCAAAGG 1016
AGAP012588-RA      GGTGCAAGGTTTCTTCACGAATAGATTGAAACACGCATTAGAACGATCTAAGGCCAAAGG 1004
                   ***** *********** ******************************************

AGAP012450-RA      TCTACGCAAATCAACTAGCCGTAAAAGAAAATTAATATAA 1056
AGAP012715-RA      TCCACGCAAATCAACTAGCCGTAAAGGAAAATTAATATAA 1056
AGAP012588-RA      TCTACGCAAATCAACTAGCCGTGAAAGAAAATTAATATAA 1044
                   ** ******************* ** **************
```

Ping-pong network 14:
Sequence Logo 3.5.0 (toolshed.g2.bx.psu.edu/repos/devteam/weblogo3/rgweblogo3/3.5.0)

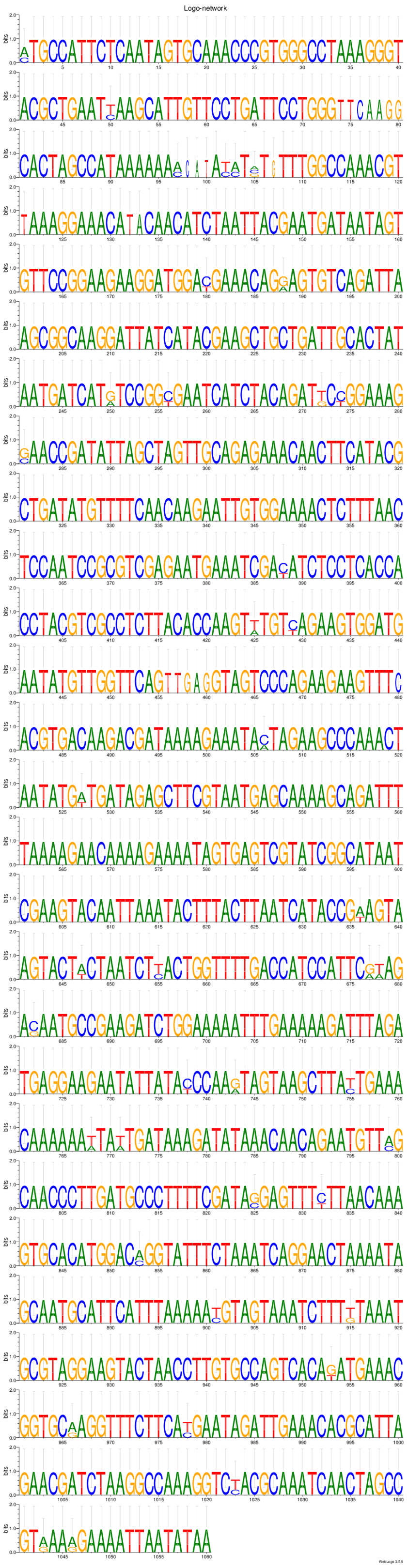

Ping-pong network 15:
CLUSTAL 2.1 multiple sequence alignment

```
AGAP012455-RA      ------------------------------------------------------------
AGAP012701-RA      ATACCGTCGGAGCCGTGTATTGCGGCGTCTGCATATGGTGCACAAATGCCGTCATCGATT 60


AGAP012455-RA      ------------------------------------------------------------
AGAP012701-RA      GCAACAATGCCAAGTTCGACTGGCTCATTATACACCATGACGCCACCAACTTCAGTTGCA 120


AGAP012455-RA      ------------------------------------------------------------
AGAP012701-RA      GTCACGTCTTCGATTGTGTCTTCAACAAGAATGGCCACAGCAACTTTGCTTCCATCGGGG 180


AGAP012455-RA      ------------------------------------------------------------
AGAP012701-RA      ACATTCATTAGAACGGTTGCACCATCCACGCCTGCGCAGACAACATATGTTGTAGCTTCA 240


AGAP012455-RA      ------------------------------------------------------------
AGAP012701-RA      TCGACTGCGACTACATCAACGGTAGCATCATCTCATTACATCCATTCAGCATGGCGTGCA 300


AGAP012455-RA      ------------------------------------------------------------
AGAP012701-RA      CCCCCACCGACACCCGCACCCATTTTTTCGATGCAGCCGACCTCAATCAAGTTCCCTTCT 360


AGAP012455-RA      GCACCTGTGGTGC--ATGGTTCA-CCTGTTGTGCCTTCG-ACTTCAACACGA---TGCCC 53
AGAP012701-RA      GGGACCATAATACGGATGGTTCAACCTGGAGTAAATCCGCAAACCAATCCAACTGTGCCT 420
                   *   *  *  * *  ******** ****  **   * ** *   ***  * *   ****

AGAP012455-RA      GCGTCCT--ATCGGCACGGTTTTACGGGCTAATATACCGT-CGGAGCCGTGTATTGCGGC 110
AGAP012701-RA      GTGCCTTCAATTGGAATGCCTTCCACAGCAAGACTTCCATGTGGAAGAACGGTTCAGATC 480
                   * * * *  ** ** * *  **     ** *     ** *  ***     *  *     *

AGAP012455-RA      GTCTGCATATGGTGTACAAATGCCGT---CATCGATTGCAACAATGCCAAGTTCGACTGG 167
AGAP012701-RA      GGCCAACTAGCAT-TGAATGTGCCGGGTGCATCAGTTATGCATTCGTCTGGTACGGCTA- 538
                   * *    **   * *  *  *****    ****  **        * *  ** ** **

AGAP012455-RA      CTCATTATACACCACGACGCCACCGAC---TTCAGTTG--CAGTCA-CGTCTTCGATTGT 221
AGAP012701-RA      --CGATTTCGACCGCGCAACCATCGACCAATCCAACCAATCGACCAGCATTGGTGTCTGT 596
                     *  * *  *** **   *** ****   * **      *   ** * *    *  ***

AGAP012455-RA      GTCTTCAACAAGAATGGCCACAGCAACTTTGCTTCCATCGGGGATATTCATTAGAACGGT 281
AGAP012701-RA      GCCTTCGCTGGGAACGTCAACTGCTCCTATGCTTCCTTATGGAACCACAATCCGTACCAT 656
                   * ****     *** * * ** **  ** ******* *  ** *     **  * **  *

AGAP012455-RA      TGCACCATCCACGCCTGCGCAGACAACATATGTTGTAGCTTCATCGACTGCGACTACATC 341
AGAP012701-RA      TGTACCATCTGCAACCGTT------ACATCTCCGTCGACTACAGTAACAGCAACAGC-CC 709
                   ** ******  *  * *        **** *       ** **   ** ** **  *  *

AGAP012455-RA      AACGCCAGGCACTTCTTACTCGGACGATAAATCAGCTTTGACTTCTACAGTGTATGCAAC 401
AGAP012701-RA      A--GTCAGGCACTTCTTACTCGGACGATACATCAGCTTCGACTTCTACAGTGTATGCAAC 767
                   *  * ************************ ******** *********************

AGAP012455-RA      TTCGTTGCAAATGGCTGCATCATCACTTCATTCACCGGTGTCAGCGCTTCGACCTTCCGT 461
AGAP012701-RA      TTCGTTGCAAATGGCTGCATCATCACTTCATTCACCGGTGTCAGCGCTTCGACCTTCCGT 827
                   ************************************************************

AGAP012455-RA      TAAACCACTGTCAGGAACGACTAAGGCAACGCCAATCGCATCTTATTCATCTGGTAGAAG 521
AGAP012701-RA      TAAACCACTGCCAGGAACGACTAAGGCAACGCCAATCGCATCTTATTCATCTGGTAGAAG 887
                   ********** *************************************************

AGAP012455-RA      TACAGCACCTTCCATTCACATTACTTCAATAGAATTGCTTCCTTCCTTGACCATGACACC 581
AGAP012701-RA      TACAGCACCTTCCATTCACATTACTTCAATAGAATTGCTTCCTTCCTTGACCATGACACC 947
                   ************************************************************

AGAP012455-RA      ACCAGAATCACCACAAATAAAATCAATCGGGTCAGATTCGGTCACATCGCAGCTTGCCGT 641
AGAP012701-RA      ACCAGAGTCACCACAAATAAAATCAATCGGGTCAGATTCGGCCACATCGCAGCTTGCCGT 1007
                   ****** ********************************** ******************

AGAP012455-RA      TTCCCGGTGTACGCAGCCATCAACATCCGGTATACTCTTAAAAAATTCATCGGTTGTGCC 701
AGAP012701-RA      ATCCCGTTGTACGCAGCCATCAACATCCGGTATACTCATAAAAGATTCATCGGTTGTGCC 1067
                    ***** ****************************** ***** ****************

AGAP012455-RA      TACAAAATCAAAAAGTGCAACAATTACAGAAAACTTACCTACTGCTGCCATAGTGTCGTC 761
AGAP012701-RA      TACAAAATCAAAAAGTGGAACAATTACAGAAAACTTACCTACTGCTGCCATAGTGTCGTC 1127
                   ***************** ******************************************

AGAP012455-RA      GGAAACATCTCAAAATCTGGACTCGAGCATCAGCAGCGTGGAGTCTATG----------- 810
AGAP012701-RA      GGAAACATCTCAAAATCTGGACTCGAGCATCAGCAGCCTGGAGTCTATGGTAGATATGTT 1187
                   ************************************* ***********
```

```
AGAP012455-RA       ------------------------------------------------------
AGAP012701-RA       ACCAGATTTCGATAGCAATAATGATACGATCCCAAAAAACCAGGAACACATT 1239
```

Ping-pong network 15:
Sequence Logo 3.5.0 (toolshed.g2.bx.psu.edu/repos/devteam/weblogo3/rgweblogo3/3.5.0)

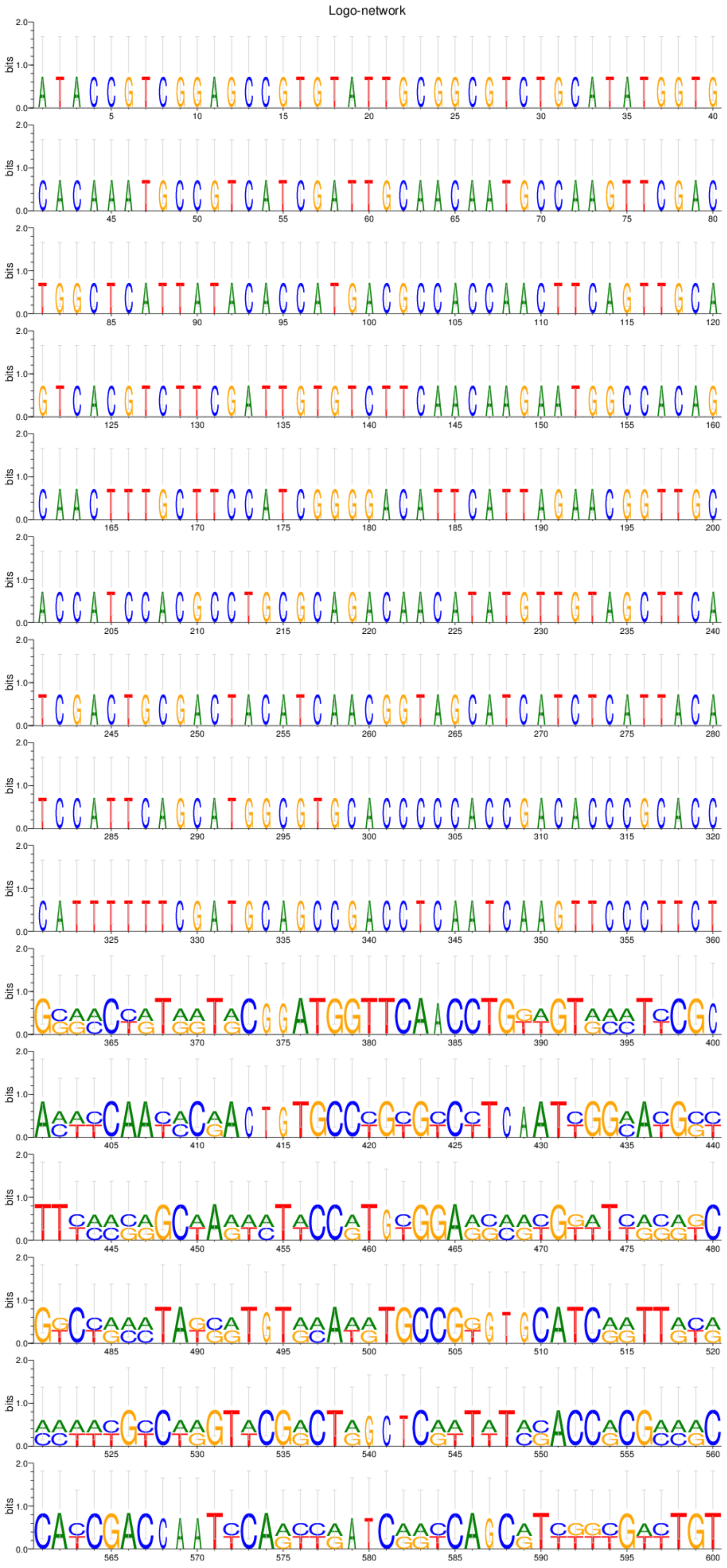

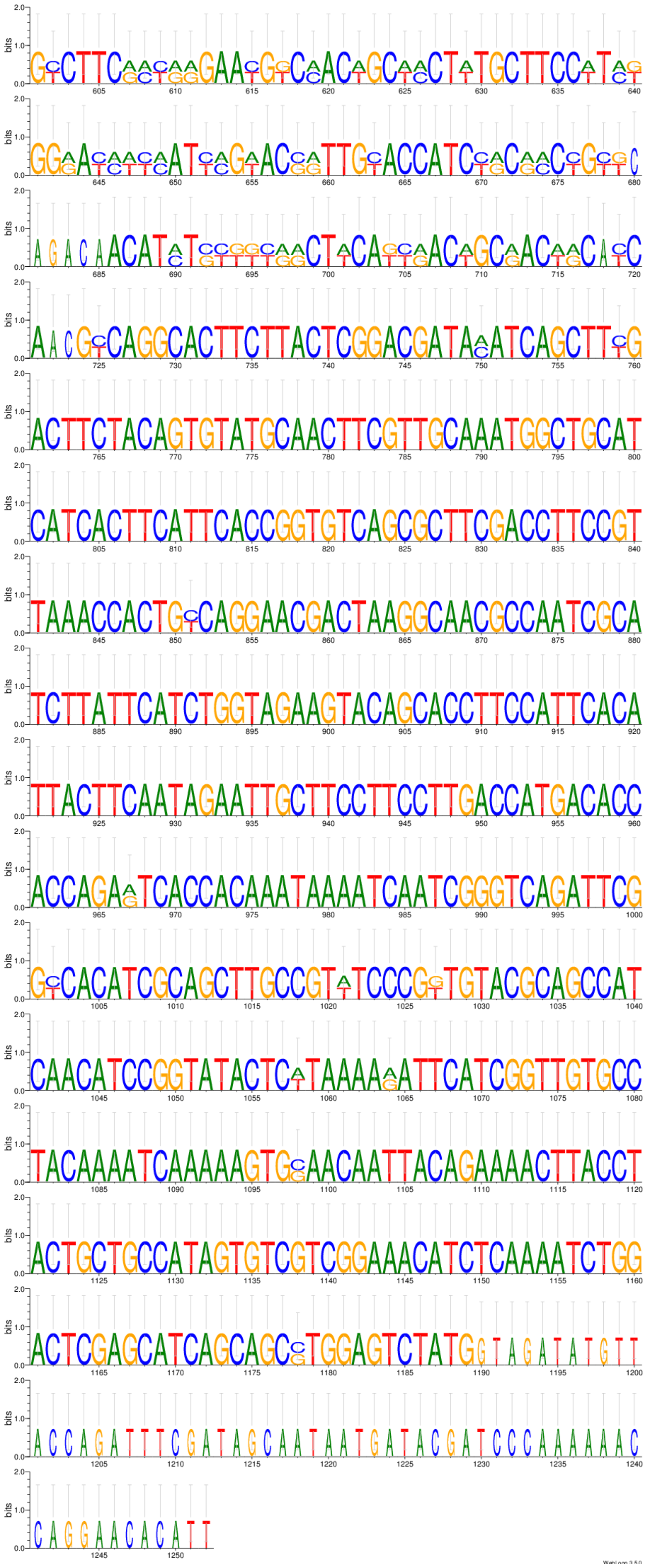
bits
WebLogo 3.5.0

```
Ping-pong network 16:
CLUSTAL 2.1 multiple sequence alignment


AGAP012560-RA      ------------------------------------------------------------
AGAP012710-RA      ATGGCACCCAAAACCAGTGGAAAAGCTGCGAAGAAGTCTGGCAAGGCCCAGAAAAATATT 60


AGAP012560-RA      ---------NACAAGAAAAAGAAGCGCAAGACTCGCAAGGAAAGCTACGCTATTTACATC 51
AGAP012710-RA      TCCAAGTCCGACAAGAAAAAGAAGCGCAAGACCCGCAAGGAAAGCTACGCTATTTACATC 120
                             ********************** ***************************

AGAP012560-RA      TACAAAGTGTTGAAGCAAGTCCACCCGGATACTGGCATCTCTTCGAAGGCCATGAGCATC 111
AGAP012710-RA      TACAAAGTGTTGAAGCAAGTCCACCCGGATACTGGCATCTCTTCGAAGGCCATGAGCATC 180
                   ************************************************************

AGAP012560-RA      ATGAACAGTTTCGTCAACGATATCTTCGAACGCATTGCTGCTGAGGCATCCCGCTTGGCG 171
AGAP012710-RA      ATGAACAGTTTCGTCAACGATATCTTCGAACGCATTGCTGCTGAGGCATCCCGCTTGGCG 240
                   ************************************************************

AGAP012560-RA      CACTACAACAAGCGTTCGACGATCACGTCCCGCGAAATCCAAACCGCTGTTCGTCTGCTG 231
AGAP012710-RA      CACTACAACAAGCGTTCGACGATCACGTCCCGCGAAATCCAAACCGCTGTTCGTCTGCTG 300
                   ************************************************************

AGAP012560-RA      CTGCCTGGTGAGCTTGCCAAGCACGCCGTCTCCGAAGGAACGAAGGCTGTCACAAAGTAC 291
AGAP012710-RA      CTGCCTGGTGAGCTTGCCAAGCACGCCGTCTCCGAAGGAACGAAGGCTGTCACAAAGTAC 360
                   ************************************************************

AGAP012560-RA      ACCAGCTCGAAGTAA 306
AGAP012710-RA      ACCAGCTCGAAGTAA 375
                   ***************

Ping-pong network 16:
Sequence Logo 3.5.0 (toolshed.g2.bx.psu.edu/repos/devteam/weblogo3/rgweblogo3/3.5.0)
```

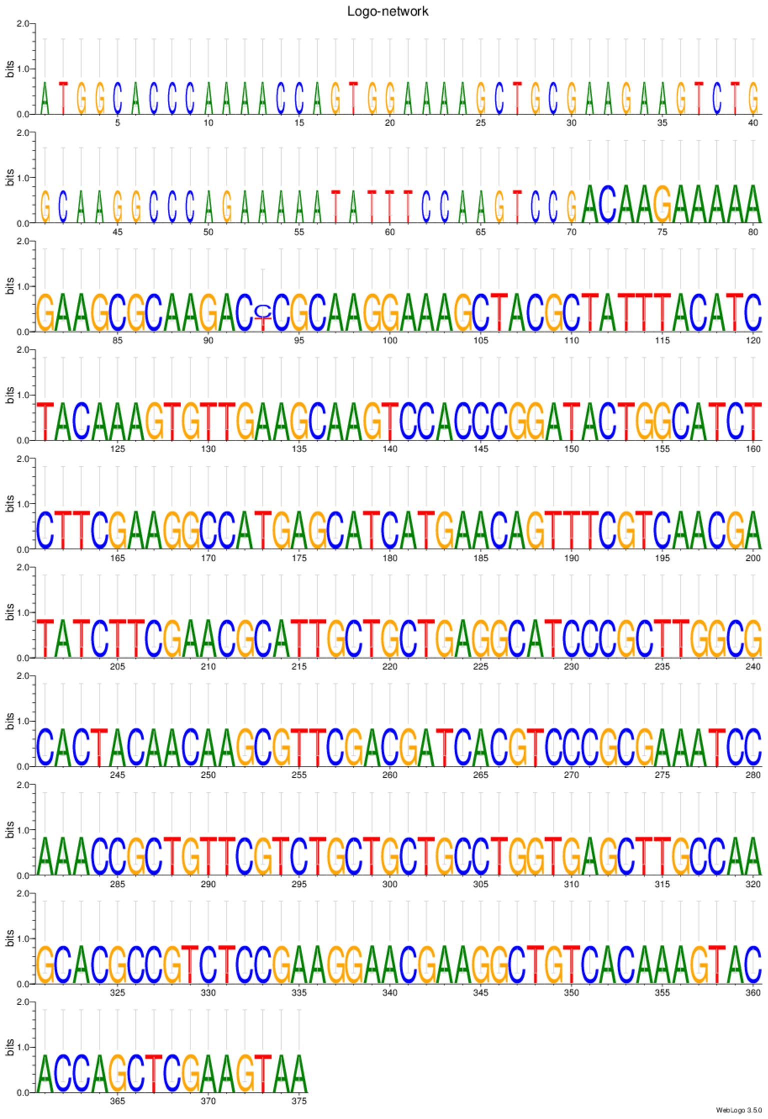

Ping-pong network 17:
CLUSTAL 2.1 multiple sequence alignment

```
AGAP013541-RA      ------------------------------------------------------------
AGAP027986-RA      ATGGATTTCATGCAGTATTTACACGGAGTGGAACGGGAGAAAGCAAACATTCCCGTAGCG 60
                                                                               

AGAP013541-RA      ----------------------------------------------------------AT 2
AGAP027986-RA      AGTCGGACGGCAGGACGCAATAGTTCCTTCACAACGTCCTTTGGCTCGCTCGTAATGTGT 120
                                                                              *

AGAP013541-RA      GGATTTTATGCAGTATTTACACGGAGTGGAACGGGAGAAAGCAAACCCGTGCGTCACGGG 62
AGAP027986-RA      GAAGCTGTAGCAGAATCTTCAC---GTGAGGCGGGCACGAGCAG----GTGCGTCACGGG 173
                   * *  *   **** ** * ***   ***   ****    ****     ************

AGAP013541-RA      CCGTGACGCGACACCAAGAATGCAGGTGCGGGACCACATCGCGACAGATTCGATCCATGA 122
AGAP027986-RA      CCGTGACGCGACACCAAGAATGCAGGTGCGGGACCACATCGCGACAGATTCGATCCATGA 233
                   ************************************************************

AGAP013541-RA      CTCCAGGGCTGCAGTAATGACTACCGGCAACATAGGTGAGACCTTTTCCAACTATCGGGA 182
AGAP027986-RA      CTCCAGGGCTGCAGTAATGACTACCGGCAACGTAGGTGAGACCTTTTCCAACTATCGGGA 293
                   ******************************* ****************************

AGAP013541-RA      CGTAGCGTATCGTGTCTATCGGGAATTGGTGCTTCAATTTACGCAAGCGGAAGCGTTGCG 242
AGAP027986-RA      CGTAGCGTATCGTGTCTATCGGGAATTGGTGCTTCAATTTACGCAAACGGAAGCGTTGCG 353
                   ********************************************** *************

AGAP013541-RA      TCTGCAAAACGCTGAGACTCAGCGTAAGATCCGCGAATCACAGAAAAGACAGCAACGAAT 302
AGAP027986-RA      TCTGCAAAACGCTGAGACTCAGCGGAAGATCCGCGAATCACAGAAAAGACAGCAACGAAT 413
                   ************************ ***********************************

AGAP013541-RA      AAAGCAACTAGAGCGTGAGTTAAATGATATCCGTGAGCTTGAAAAGGATGAGGAACGGTG 362
AGAP027986-RA      AAAGCAACTAGAGCGTGAG----------------------------------AACGGTG 439
                   *******************                                  *******

AGAP013541-RA      CCGGGCAACAGATCGTTCGGGCGCACCGTATCCTGATCAATTTGGCTACGGGCATCACGA 422
AGAP027986-RA      CCCTGGAACGG-----------------TATCCTGATCAATTTGGCTACGGGCATCACGA 482
                   **  * *** *                 ********************************

AGAP013541-RA      GCCGCACGGGGACATGTTCCGGGGTCAAACCCCTTCCCCGCCGTGTATTGACCTCGACCG 482
AGAP027986-RA      GCCGCACGGGGACATGTTCCGGGGTCAAACCCCTTCCCCGCCGTGTATTGACCTCGACCG 542
                   ************************************************************

AGAP013541-RA      CAGTGGTGATGCGAATCGGACGCACGTCAGTGGCGACACAACTACACACACATGCCCTGA 542
AGAP027986-RA      CAATGGTGATGCGAATCGGACGCACGTCAGTGGCGACACAACTACACACACATGCCCTGA 602
                   ** *********************************************************

AGAP013541-RA      TAAACGTCATAACCGCGCAGTGATCCACAACACACACAATCTAAAATCGTTCACTAATAC 602
AGAP027986-RA      TAAACGTCATATCCGCGCAGTGATCCACAACACACACAATCTAAAATCGTTCACTAATAC 662
                   *********** ************************************************

AGAP013541-RA      TACCAACGCGACATTACACACAAACACTGTCTTAGAAGACACGTACATCGAGGCTCATCA 662
AGAP027986-RA      TACCAACGCGACATTACACACAAACACTGTCTTAGAAGACACGTACATCGAGGCTCATCA 722
                   ************************************************************

AGAP013541-RA      CCGTTTCTGGCTGCGTCCTGCGGGGTTGAAAGATGCTACAAGTGGCGGTGAGGCCACCTT 722
AGAP027986-RA      CCGTTTCTGGCTGCGTCCTGCGGGGTTGAAAGATGCTACAAGTGGCGGTGAGGCCACCTT 782
                   ************************************************************

AGAP013541-RA      TGAAAGACAGGAATTGATCGAGGAAAGCGAAGTTTTAGAACCATCGGAGTCAGGTGCTTA 782
AGAP027986-RA      TGAAAGACAGGAATTGATCGAGGAAAGCGAAGTTTTAGAACCATCGGAGTCAGGTGCTTA 842
                   ************************************************************

AGAP013541-RA      CTTCGAACCCTGCGTTGCAATCGAGCTCGAGGAAGTCATGCCACGACTGAGCTTGAAACC 842
AGAP027986-RA      CTTCGAACCCTGCGTTGCAATCGAGCTCGATGAAGTCATGCCACGACTGAGCTTGAAACC 902
                   ****************************** *****************************

AGAP013541-RA      ATGGTCTGAGGTGCTGCAGCTAGGAGAACGAAGCACTGGAGGGGAGACATACTCGCGCGA 902
AGAP027986-RA      ATGGTCTGAGGTGCTGCAGCTAGGAGAACGAAGCACTGGAGGGGAGACATACTCGCGCGA 962
                   ************************************************************

AGAP013541-RA      CGGCTCGAACCATGTCCCACGTTCGTCAAGGAACAGAATTTCTATCCCGGAATCTAGAGA 962
AGAP027986-RA      CGGCTCGAACCATGTCCCACGTTCGTCAAGGAACGGAATTTCTATCCCGGAATCCAGAGA 1022
                   ********************************** ******************* *****

AGAP013541-RA      GATTGGGAGGATGGAGTCTTCTGTGTCGTCGAGAAGGGATCCGGTCTGGTCGATCGTGAG 1022
AGAP027986-RA      GATTGGGAGGATGGAGTCTTCTCTGTCGTCGAGAAGGGATCCGGTCTGGTCGATCGTGAG 1082
                   ********************** *************************************

AGAP013541-RA      GGATCTTGCCACTAAATGTTGCCTGCTGCAGTGCCACATTGGACTACCTCATGGAAAAAT 1082
AGAP027986-RA      GGATCTTGCCACTAAATGTTGCCTGCTGCAGTGCCACATTGGACTACCTCATGGAAAAAT 1142
                   ************************************************************
```

```
AGAP013541-RA      GAACGGCTTGTGGCGGTATGTACGGCTGAAACAAAAGCTTAATTTAATTTCAGCAGCAGA 1142
AGAP027986-RA      GAACGGCTTGTGGCGGTATGTACGGCTGAAACAAAAGCTTAATTTAATTTCAGCAGCAGA 1202
                   ************************************************************

AGAP013541-RA      TAAAGGGAAGTCAGGATACTCATGCCAAATCATCCGTGCGTATCGTGGTACCATCCGGTG 1202
AGAP027986-RA      TAAAGGGAAGTCAGGATACTCATGCCAAATCATCCGTGCGTATCGTGGTACCATCCGGTG 1262
                   ************************************************************

AGAP013541-RA      CTGGGAAACGTTTATAGACAGCTCTGCGA-GTACA-GTAT-CGGACAGAAAGATAGGGCA 1259
AGAP027986-RA      CTGGGAAACGTTTATAGACAGCTCTGCGAAGTATGTGTGTGCGGTCCGGTTGCAGAAGCT 1322
                   ***************************** ***   ** * *** * *   *     ** 

AGAP013541-RA      TTGGTTTTTTAACGCACCGTGCACCCGTTGGGCCAAGTTTATGTGTGGTTTGTATGTGTT 1319
AGAP027986-RA      GTGA-------------------------------------------------------- 1326
                    **

AGAP013541-RA      TGTGTTTGTGTGTGTGTGTATGTGTGTGTGTATGTGTGTGTGTGTGTGTATGTGTGTG 1379
AGAP027986-RA      ------------------------------------------------------------

AGAP013541-RA      TGTGTGTATGTGTGTGTGCATATGTGTGTGTGTGTGGATGTATGTTTGA 1428
AGAP027986-RA      -------------------------------------------------
```

Ping-pong network 17:
Sequence Logo 3.5.0 (toolshed.g2.bx.psu.edu/repos/devteam/weblogo3/rgweblogo3/3.5.0)

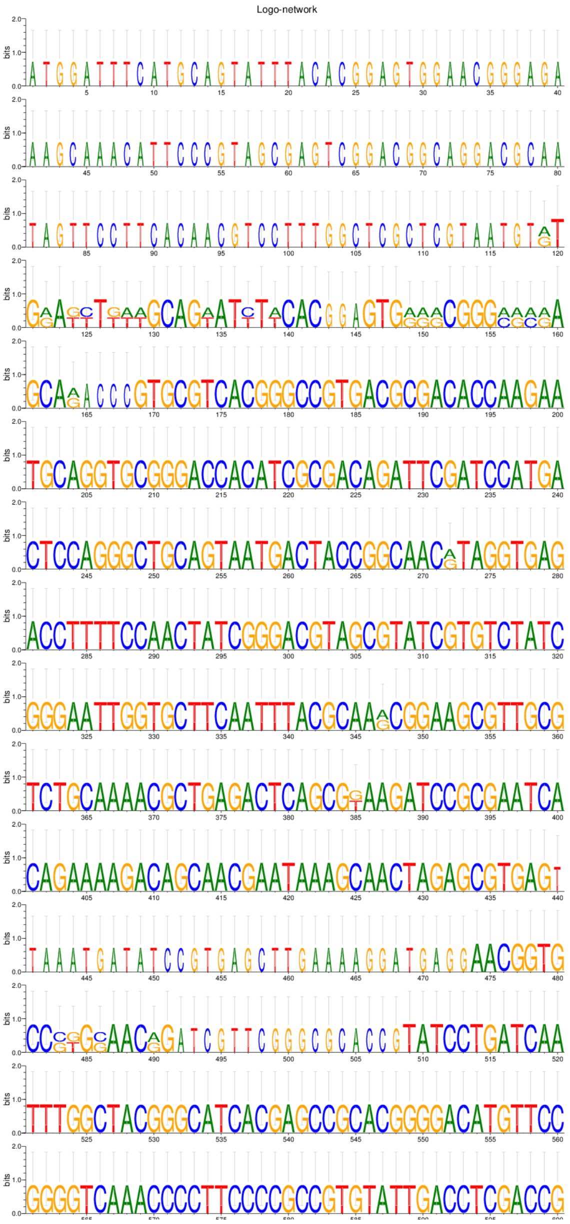

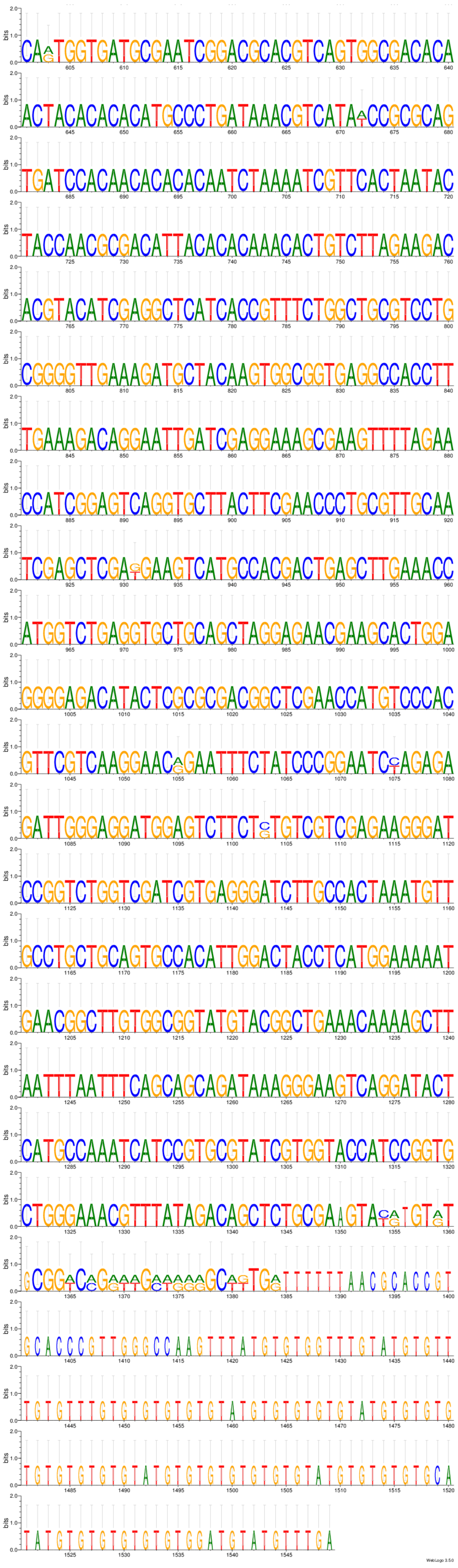
bits
WebLogo 3.5.0
