## Supplementary material for "Conserved small nucleotidic elements at the origin of concerted piRNA biogenesis from genes and lncRNAs": multiple sequence alignments SI-MSA2

### Supplemental Information: Multiple sequence alignments and consensus sequences of *An. gambiae, Ae. aegypti,* mouse and human snetDNAs

#### Multiple alignment and consensus of *An. gambiae* snetDNAs matching to 2R_lncs_14 positions 1163-1213:

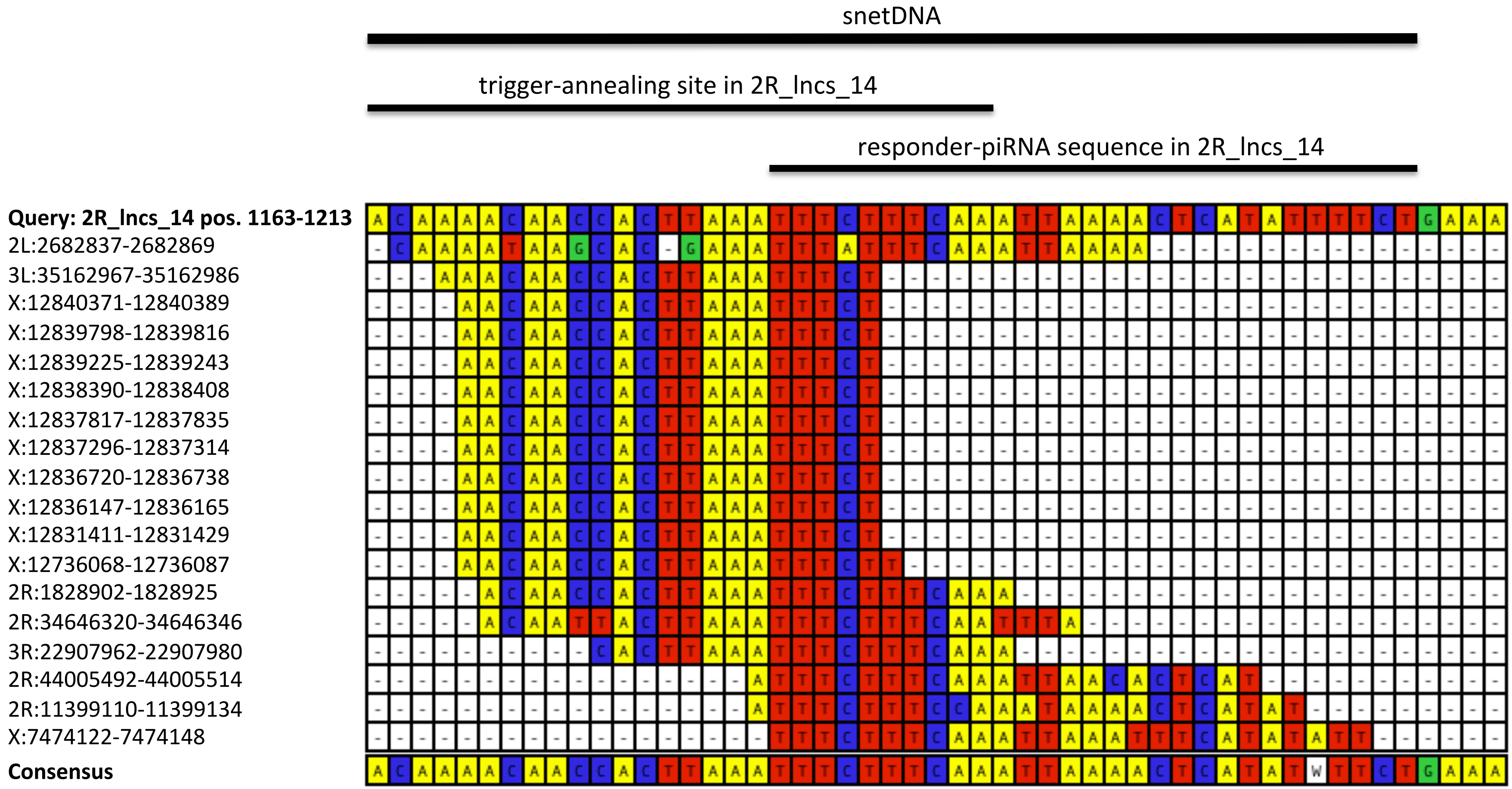

#### Multiple alignment and consensus of *Ae. aegypti* snetDNAs matching to 2R_lncs_14 positions 1163-1221:

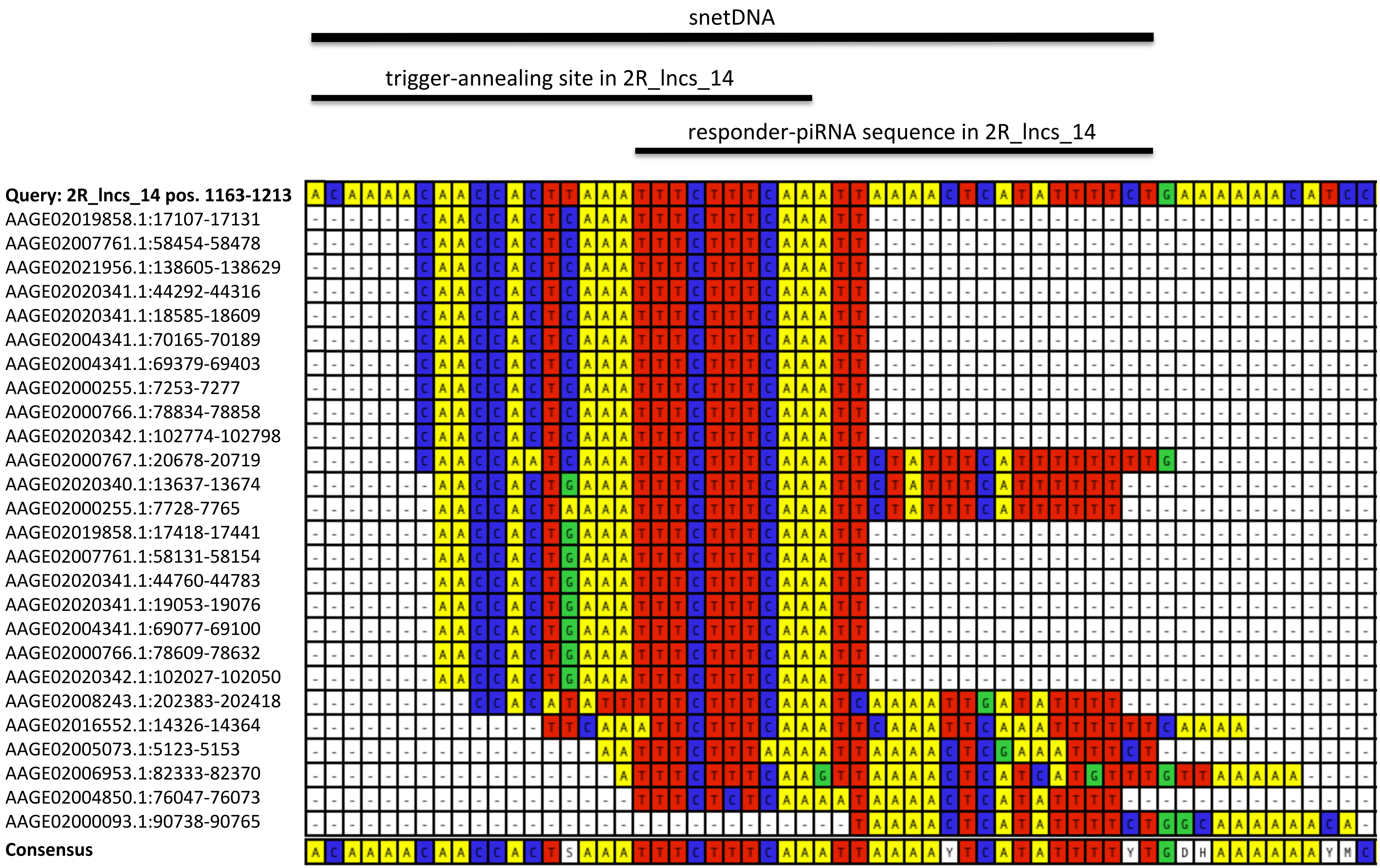

### Multiple alignment and consensus of mouse snetDNAs matching to 2R_lncs_14 positions 1163-1231:

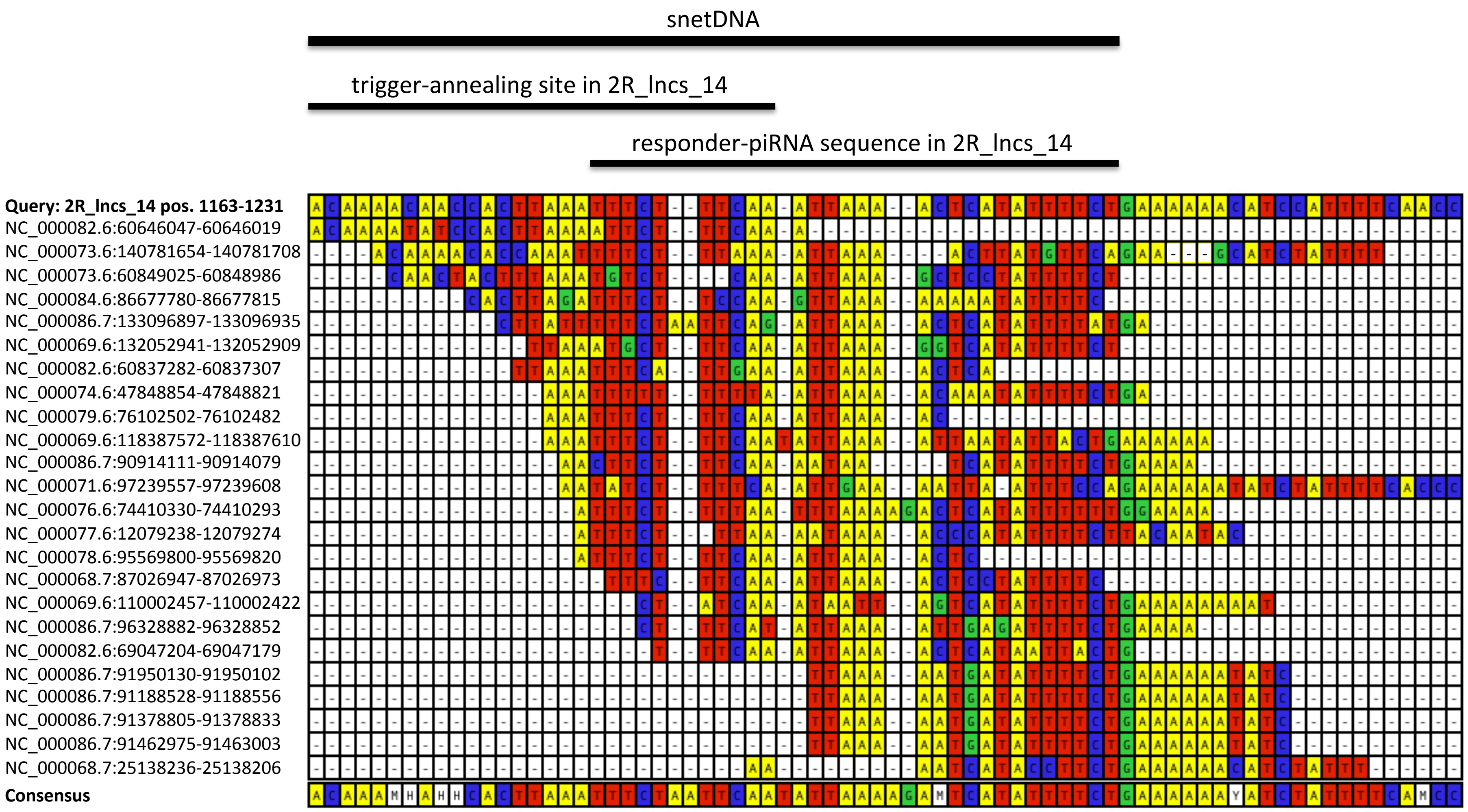

### Multiple alignment and consensus of human snetDNAs matching to 2R_lncs_14 positions 1141-1227:

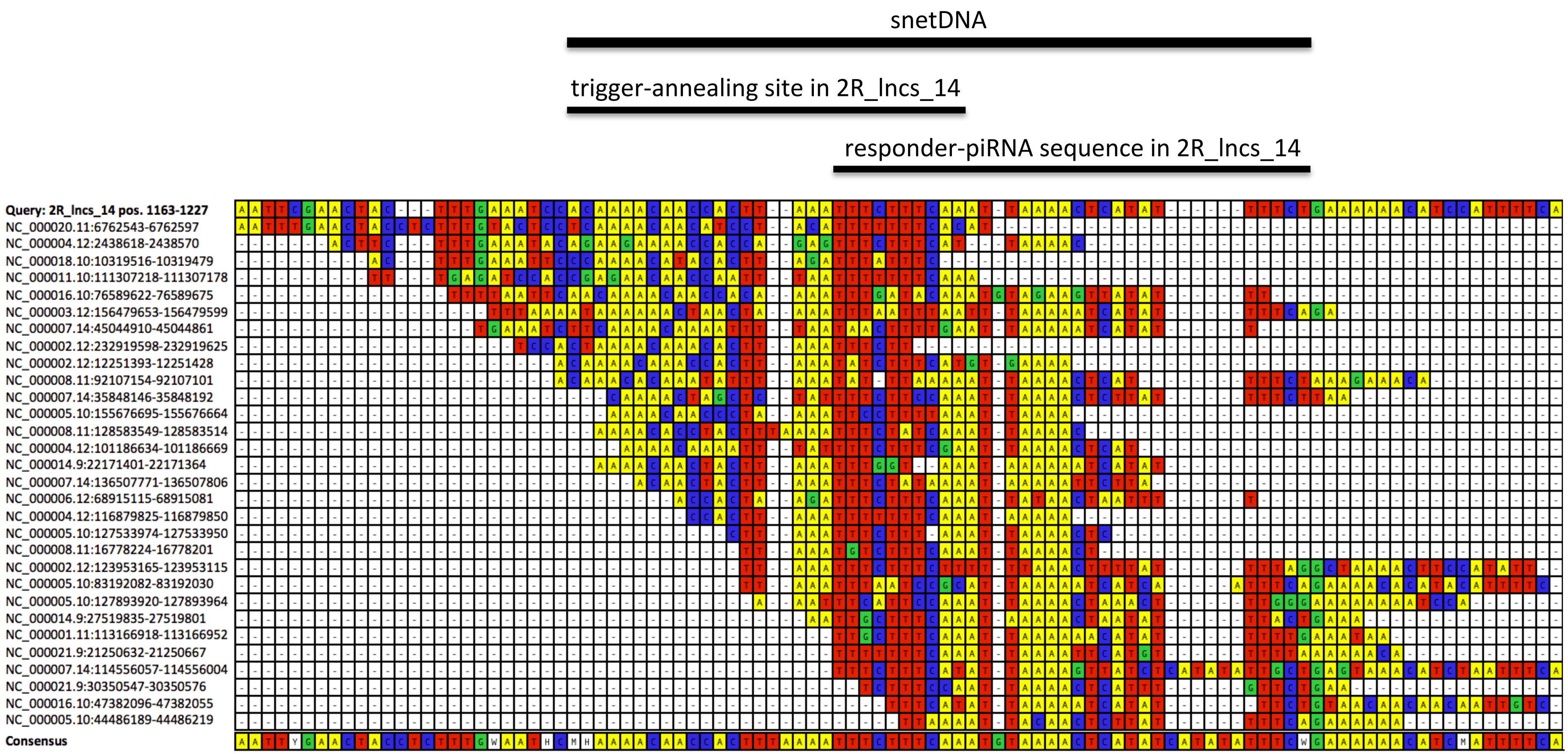

### Multiple alignment and consensus of *An. gambiae* snetDNAs matching to AGAP011923-RA positions 231-284:

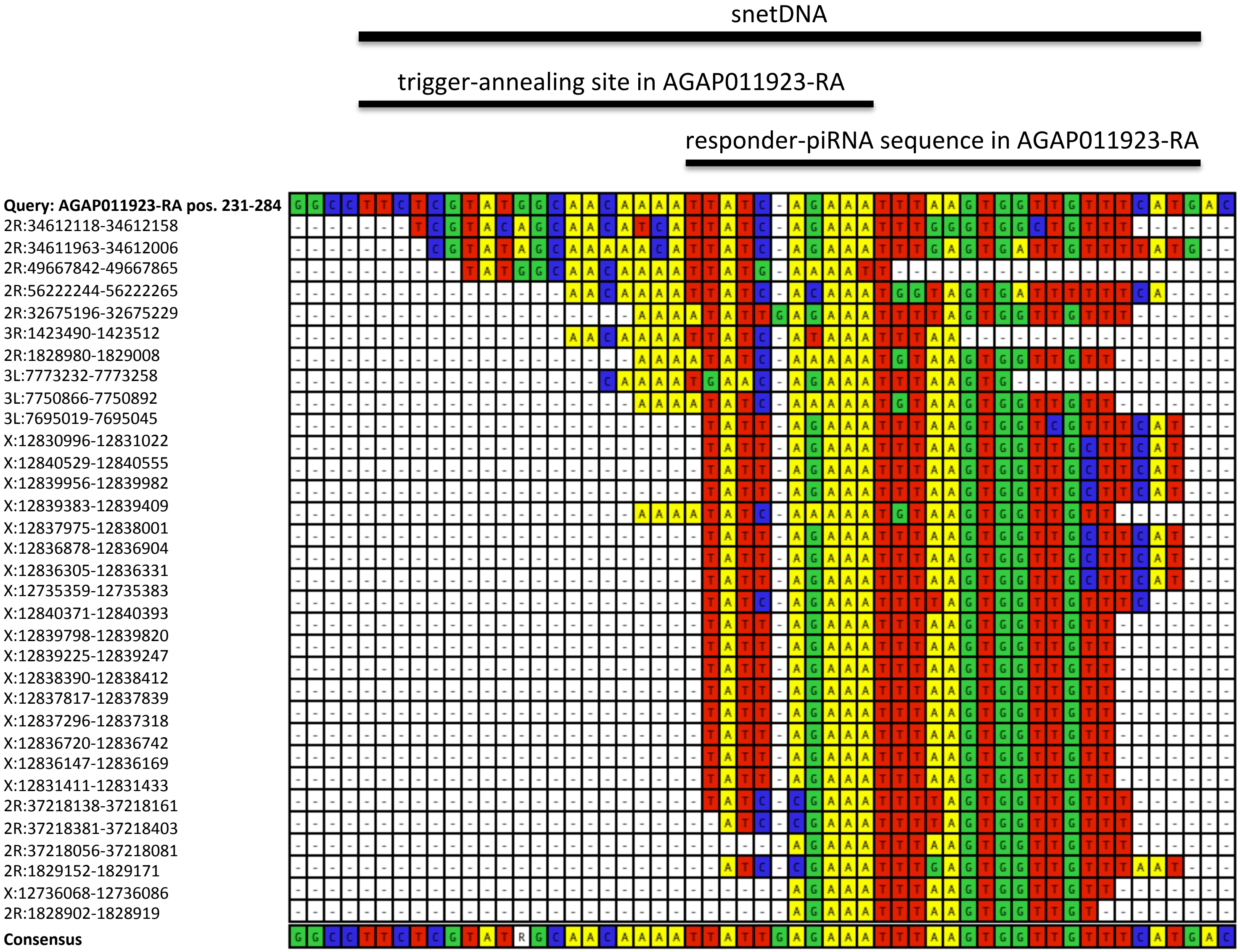

### Multiple alignment and consensus of *Ae. aegypti* snetDNAs matching to AGAP011923-RA positions 231-284:

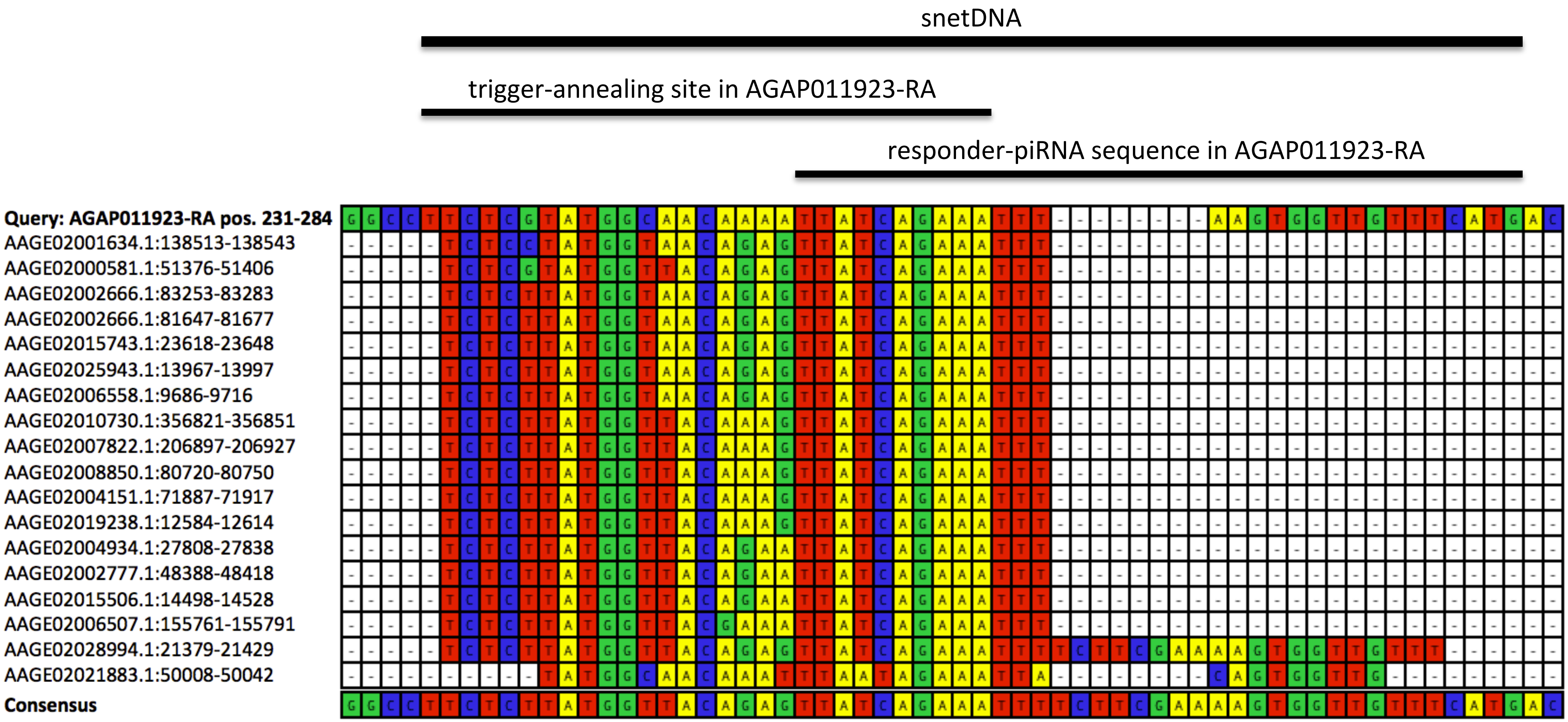

### Multiple alignment and consensus of mouse snetDNAs matching to AGAP011923-RA positions 231-284:

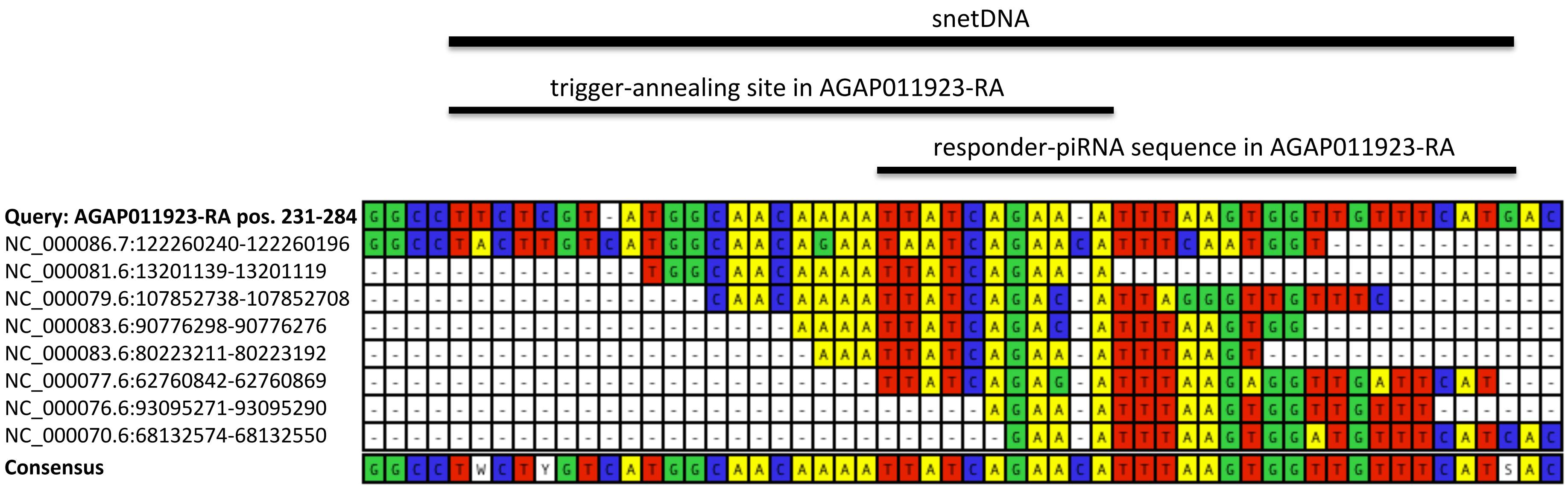

#### Multiple alignment and consensus of human snetDNAs matching to AGAP011923-RA positions 231-290:

snetDNA

trigger-annealing site in AGAP011923-RA

responder-piRNA sequence in AGAP011923-RA

| | |
|---|---|
| **Query: AGAP011923-RA pos. 231-290** | GGCCTTCTCGTATGGC-AACAA-AATTATCAGAAATTTAAGTGGTTGTTTCATGACGATGAT--- |
| NC_000001.11:215425757-215425793 | --CCTTCAAGTATGGCTAACACAAAATATCAGAAATTTA-------------------------- |
| NC_000004.12:19540137-19540103 | -----TCTTGTTTGGA-AACAA-AATTATGAGAAATCAAAGT----------------------- |
| NC_000002.12:10443999-10444050 | ---------GTATAGA-AATAT-AATAATCAGAAATTTAAGT--TTGTTTGATAAGAATCCACAT |
| NC_000009.12:26175790-26175821 | ---------GTATGCC-CACAATAATTATCAGAAATTGAAGT----------------------- |
| NC_000023.11:113736746-113736708 | ----------TGGCAA-AATCA-ACTTATCAAAAATGTAAGTGTTTGTTTC-------------- |
| NC_000010.11:60478615-60478595 | -----------------AACAA-AATTATCAGAAATTTA-------------------------- |
| NC_000006.12:101351131-101351097 | --------------------AA-AGTAATCAGAATTTTAAGTGTTTGTTACATTAC--------- |
| NC_000002.12:197087120-197087090 | ---------------------A-AATTATTATAGGTTTAAGTGGTTGTTTCAT------------ |
| NC_000020.11:48021859-48021893 | ---------------------A-AATTGTCAGAAATTTAAGTGTGTGTGTTCTTGAC-------- |
| NC_000014.9:43878706-43878739 | ----------------------------TCAGAAATTTAACTAGTTGATTCATGACCAAGAT--- |
| **Consensus** | GGCCTTCTHGTATGGMTAACAAWAATTATCAGAAATTTAAGTGKTTGTTTCATKACVATGATCAT |
